# Functional Biosynthetic Stereodivergence in a Gene Cluster via a Dihydrosydnone *N*-oxide

**DOI:** 10.1101/2024.04.29.591611

**Authors:** Jiajun Ren, Anugraha Mathew, Maria Rodriguez Garcia, Tobias Kohler, Olivier Blacque, Anthony Linden, Leo Eberl, Simon Sieber, Karl Gademann

## Abstract

Chirality features a critical role in the biochemistry of life and often only one enantiomeric series is observed (homochirality). Only few natural products have been obtained as racemates, e.g. the quorum-sensing signal valdiazen produced by *Burkholderia cenocepacia* H111. In this study, we investigated its biosynthetic gene cluster and discovered that both the enantiomerically pure (*R*)–fragin and the racemic valdiazen are obtained from the same pathway. This stereodivergence is based on the unusual heterocycle dihydrosydnone *N*-oxide intermediate, as evident from gene knockout, stable isotope feeding experiments, and mass spectrometry experiments. Both non-enzymatic racemisation via keto-enol tautomerisation and enzyme-mediated dynamic kinetic resolution were found to be crucial to this stereodivergent pathway. This novel mechanism underpins the production of configurationally and biologically distinct metabolites from a single gene cluster. Our findings highlight the intricate design of an intertwined biosynthesis pathway, providing a deeper understanding of microbial secondary metabolism related to microbial communication.

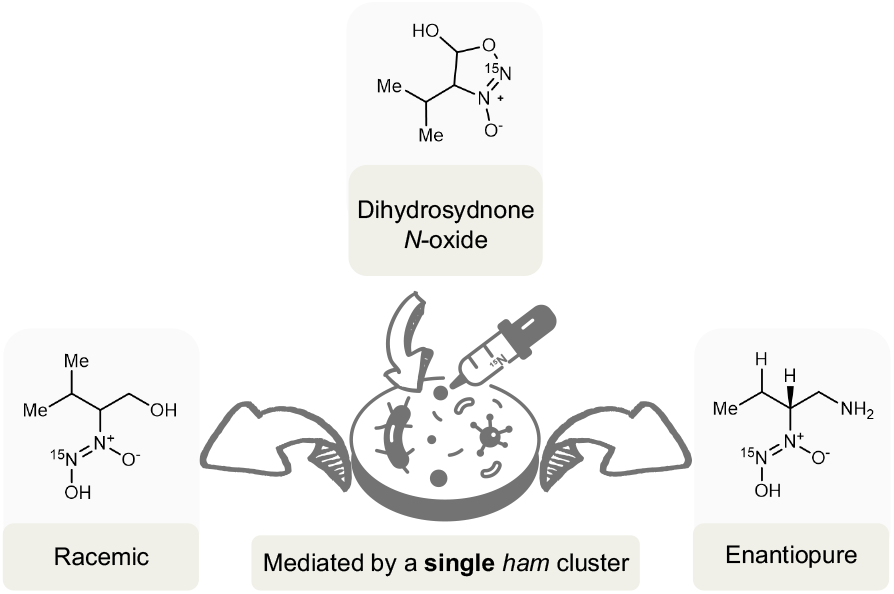

## Introduction

Biochemical homochirality describes the dominance of one enantiomer over the other across the entire biosphere on Earth. For instance, the primary metabolism of living organisms relies almost exclusively on L–amino acids to make proteins while using right-handed carbohydrates (D–monosaccharides) to make polysaccharides and DNA (**Fig 1A**).^1^ Although the origin of biochemical homochirality remains controversial,^2^ building macromolecules with one enantiomer eliminates the diastereoisomer problem,^3^ and renders biological processes more efficient.^4–7^ However, racemic secondary metabolites are sometimes produced in a homochiral biological environment, in particular regarding alkaloids and terpenoids.^8^ Different racemisation mechanisms have been documented including both enzymatic and non-enzymatic pathways.^8^ The dihydrodaidzein racemase involved in the biosynthesis of equol is an example of enzymatic pathways.^9^ Non-enzymatic pathways typically involve a reactive biosynthetic intermediate that can undergo dimerisation, cyclisation or tautomerisation (**Fig 1B**).^8,10^ In this study, we demonstrate both non-enzymatic racemisation and enzyme-mediated dynamic kinetic resolution occur in parallel and are encoded by the *ham* gene cluster in *Burkholderia cenocepacia* H111.^11^

**Fig 1.**
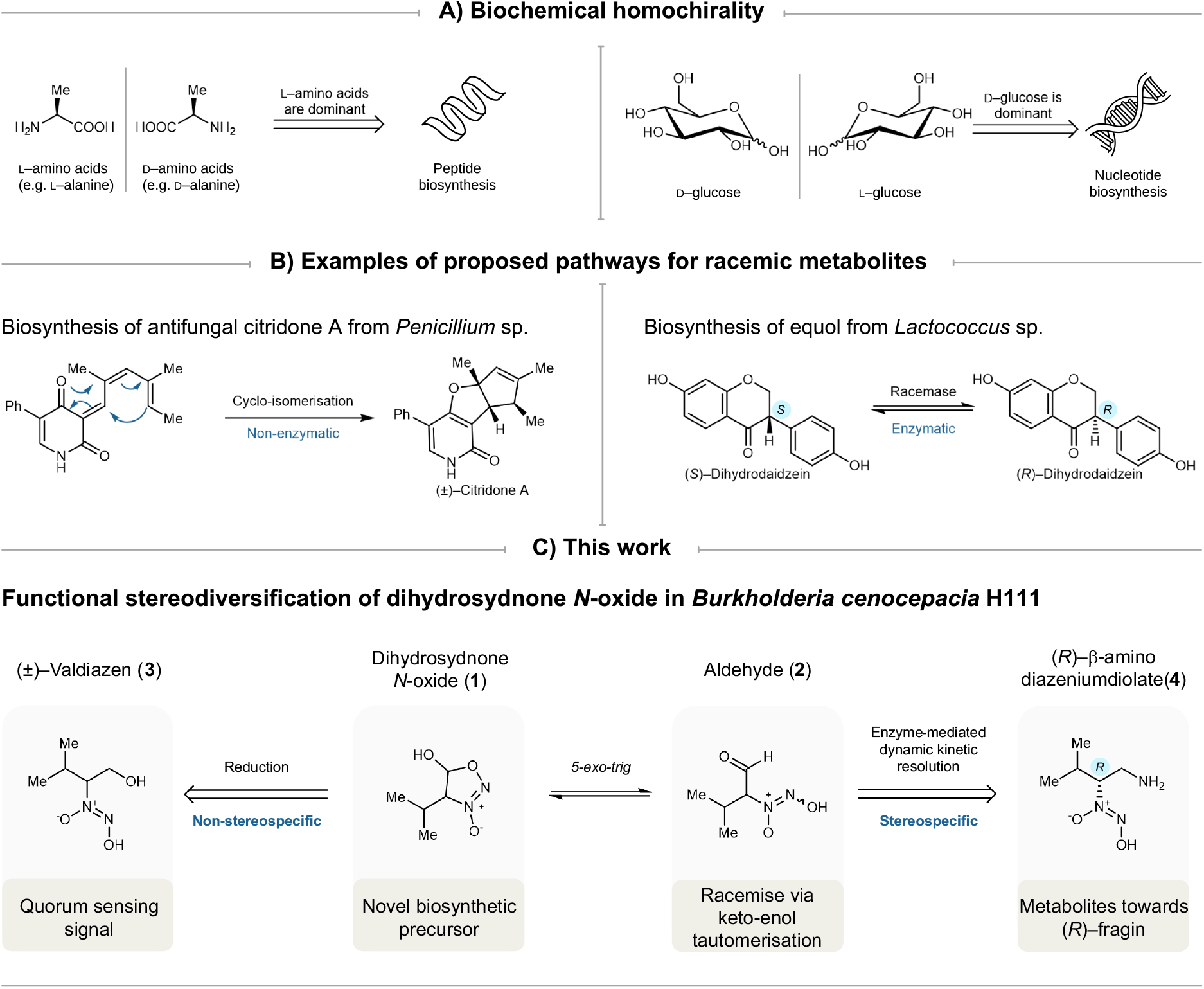
**A)** Homochiral preferences in the biosynthesis of peptides and nucleotides.^1^ **B)** Two proposed biosynthetic pathways resulting in racemic metabolites.^9,10^ **c)** Dihydrosydnone *N*-oxide is a key intermediate at the junction of the biosynthesis of valdiazen and fragin in *Burkholderia cenocepacia* H111.

The *ham* cluster is rather unique in producing two secondary metabolites with different stereogenic configurations and distinct biological functions in contrast to other diazeniumdiolate gene clusters.^12–20^ Antifungal fragin features the (*R*)–configuration opposite to its L–valine precursor, whereas the quorum-sensing signal valdiazen was found as a racemate.^11^ This suggested that racemisation and dynamic kinetic resolution occur in the biosynthesis of the two molecules. Moreover, valdiazen regulates the production of fragin and autoregulates its own production via a positive feedback loop, i.e. valdiazen induces its own and fragin biosynthesis.^11^ In this study, we provide evidence that the diverse stereoconfigurations of secondary metabolites produced by the *ham* cluster might be the result of a finely tuned interplay between enol-keto tautomerisation and enzyme-mediated dynamic kinetic resolution, centred around the novel heterocycle dihydrosydnone *N*-oxide **1** (**Fig 1C**).

## Results

### Discovery of dihydrosydnone-*N* oxide as an intermediate

We hypothesised the existence of a common aldehyde intermediate **2** (**Fig 2**) for the biogenesis of (*R*)– fragin and (±)–valdiazen, based on previous findings from our group and others.^11,13–16,19–26^ The *ham* cluster consists of seven genes distributed in two oppositely oriented operons.^11,26^ As demonstrated by isotopically labelled feeding experiments, the biosynthesis starts with loading L–valine onto the HamD nonribosomal peptide synthetase (NRPS) complex.^26^ HamC, which shares homology with diiron *N*-oxygenase AurF,^27^ was found to be responsible for the *in vitro* nitrogen oxidation of valine, while the substrate was bound to the NRPS complex.^11,23,28^ The cupin domain of HamB was suggested nonessential for the production of valdiazen,^11^ and that HamA (PvfA) and HamE (PvfD) were nonessential for the production of hydroxylamine intermediates towards valdiazen and pyrazine *N*-oxide derivatives.^22^ From the second operon, HamG was predicted to be an aminotransferase, and HamF a starter condensation domain,^11^ and both of them are indispensable for the production of fragin.^11^ The NRPS HamD contains a reductase instead of the more typical thioesterase domain. A similar NRPS architecture leads to an aldehyde intermediate in the biosynthesis of myxochelins^21,29^ and myxalamid S.^30^ The C(α)-proton in aldehyde **2** is expected to have an increased acidity due to the presence of both electron-withdrawing aldehyde and diazeniumdiolate functional groups. In addition, we reason that either the aminotransferase HamG or the condensation domain HamF will determine the configuration of fragin. Consequently, tautomerisation and racemisation of such an aldehyde could be central for the stereodivergence in this biosynthetic gene cluster.

**Fig 2.**
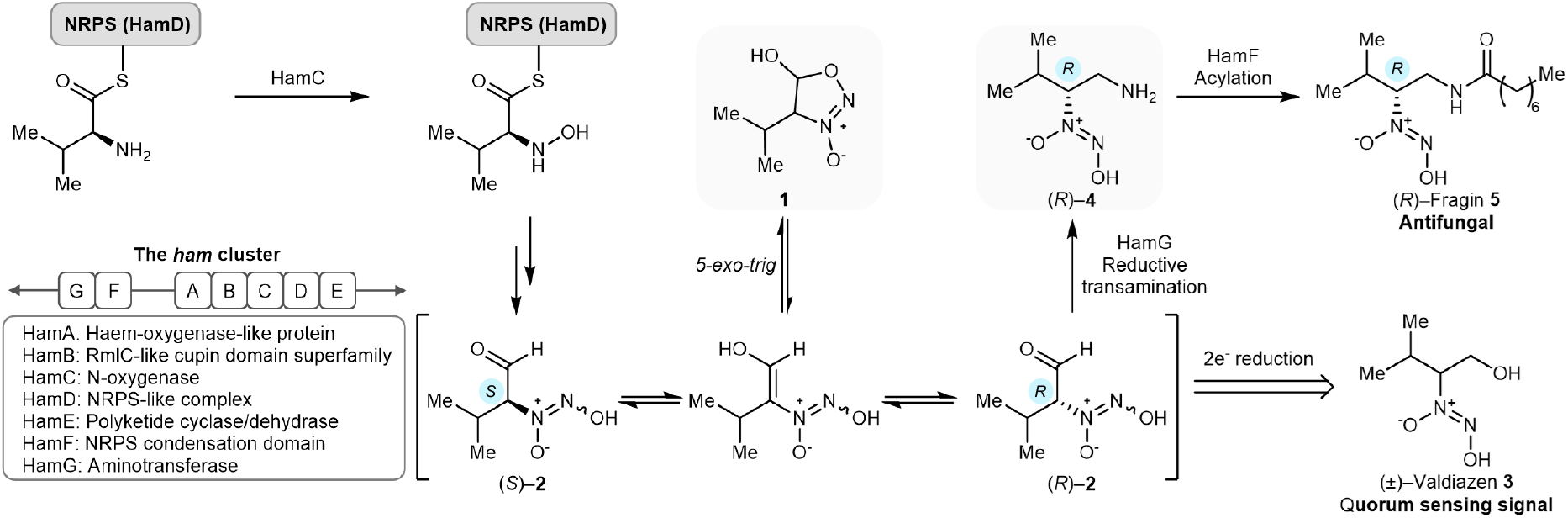
Proposed biosynthesis of (*R*)–fragin **5** and (±)–valdiazen **3** via dihydrosydnone *N*-oxide **1** mediated by the *ham* cluster in *B. cenocepacia* H111.

In order to address these hypotheses, we first synthesised standards and ^15^N-labelled tracers for feeding experiments and mass spectrometry analyses. Methyl 2,2-dimethoxyacetate **6** was transformed to ketone **7** via the respective Weinreb amide (**Fig 3A**). Ketone **7** was then reacted with hydroxylamine to oxime **8** as a mixture of (*E*/*Z*)-isomers in 81% yield. The oximes were further reduced and nitrosylated using ^15^N-sodium nitrite to give ^15^N-diazeniumdiolate **9**. Subsequent acid hydrolysis at 50 °C surprisingly did not lead to the aldehyde, but instead to hemiacetal ^15^N-**1**. This novel heterocycle dihydrosydnone *N*-oxide **1** was fully characterised including ^1^H–NMR spectroscopy (characteristic doublet at 5.95 ppm in methanol-*d*_*4*_) and unambiguously by X-ray crystal structure analysis (**Supplementary Table 3**). Interestingly, only one diastereoisomer of racemate **1** could be detected by 1H–NMR spectroscopy. The heteroatom-rich molecular architecture of **1** constitutes the first example of a dihydrosydnone *N*-oxide. The corresponding oxidised form, sydnone *N*-oxide (Traube’s anion), was discovered in 1895.^31^ Since then, only a few derivatives of this aromatic family were synthesised mostly using NO gas.^31–36^ Notably, molsidomine containing a sydnone imine core has been approved as a vasodilating drug for angina pectoris.^37,38^ Other similar heterocycles (**Supplementary Table 1**) including 4,5-dihydro-1,2,3-oxadiazole,^39^ and 5-hydroxy-3-methyloxadiazolinium perchlorate have been synthesised.^40^ When comparing their bond lengths, it is interesting that the sydnone *N*-oxide anion has the longest N-N bond while molsidomine has the shortest C-C bond in the cycle (**Supplementary Table 1**).

**Fig 3.**
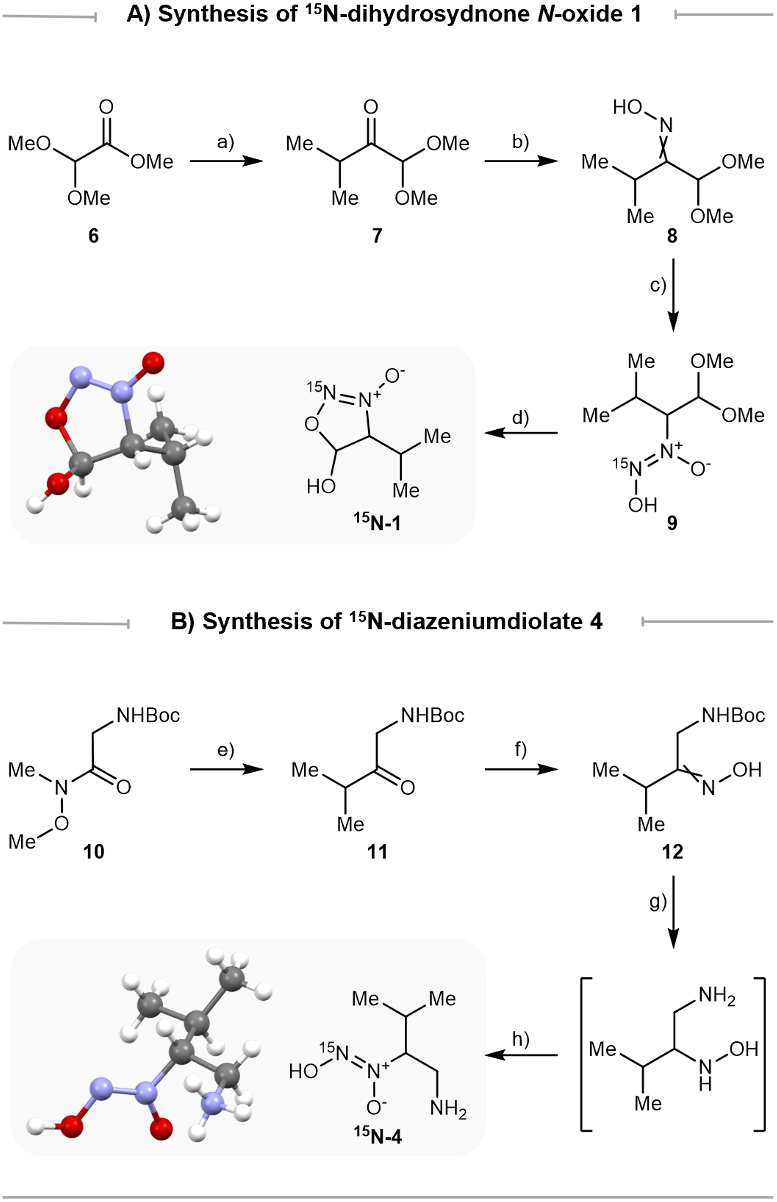
Synthesis of ^15^N-labelled tracers and X-ray crystal structures of unlabelled compounds **1** and **4**. Colour code: blue = nitrogen; red = oxygen; grey = carbon; white = hydrogen. **A)** Synthesis of ^15^N-dihydrosydnone *N*-oxide **1**: a) AlMe_3_, *N*-methoxy-methanamine·HCl, 4.5 h, then isopropyl magnesium bromide, 2 h, 56% over 2 steps; b) NH_2_OH·HCl, NaOAc, EtOH/H_2_O, 70 °C, 3 h, 81%; c) NaCNBH_3_, HCl, EtOH, 1 h, then 1 M aq. HCl, Na^15^NO_2_, 1 h, 55% over 2 steps; d) 10% aq. HCl, 50 °C, 3 h, 46%. **B)** Synthesis of ^15^N-diazeniumdiolate **4**: e) isopropyl magnesium bromide, 2 h, 49%; f) NH_2_OH·HCl, KOAc, EtOH/H_2_O, 70 °C, 3 h, 89%; g) NaCNBH_3_, HCl, EtOH/H_2_O, 1 h, then TFA, DCM, 1 h; h) 1 M aq. HCl, Na^15^NO_2_, 30 mins, 21% over 3 steps.

The ^15^N-labelled amine analogue **4** was synthesised to interrogate the role of the predicted condensation domain HamF. The commercially available Weinreb amide **10** was reacted with a Grignard reagent to the ketone **11**, which was subsequently transformed to the mixture of oximes **12** (**Fig 3B**). A three-step reaction sequence involving oxime reduction, Boc deprotection and nitrosylation with ^15^N-sodium nitrite led to the amine ^15^N-**4**. The racemic **4** was crystallised from a methanol/ether mixture and its molecular structure was confirmed by X-ray crystal analysis. (**Supplementary Table 4**).

### Dihydrosydnone *N*-oxide 1 is an intermediate at the junction between the biosynthesis of valdiazen and fragin

With both isotopically labelled compounds in hand, feeding experiments were performed in *B. cenocepacia* H111 mutants *ΔhamC, D*, and *G*, to investigate whether ^15^N-**1** and ^15^N-**4** can be incorporated into the biosynthetic pathway. To detect both fragin enantiomers, a separation method was developed on an ultra-high-performance liquid chromatography (UHPLC) instrument connected to a high-resolution mass spectrometer (HRMS). A screening of columns was performed, and the chiral Lux^®^ i-Amylose-3 (*Phenomenex*) was selected for its best separation performance. With the developed method (**see Method**), both fragin enantiomers were differentiated by directly injecting the cell-free supernatant. All feeding experiments were performed in biological triplicates (**Supplementary Table 2**). We observed stereoselective (*R*)–^15^N-fragin production when supplementing racemic dihydrosydnone *N*-oxide ^15^N-**1** in a *ΔhamC* and *ΔhamD* mutant. In contrast for the *ΔhamG* mutant, compound **4** was instead required to produce (*R*)–^15^N-fragin (**Fig 4A and B**). As expected, in the double mutants *ΔhamCG* and *ΔhamDG*, supplementation of dihydrosydnone *N*-oxide ^15^N-**1** did not result in fragin production (**Supplementary Fig 1**). To fully ascertain the stereogenic outcomes of ^15^N-fragin production, cell-free supernatants of all mutant cultures were spiked with enantiomerically pure (*R*)– and (*S*)–fragin, respectively (**Fig 4C and Supplementary Fig 2**). A clear distinction between analytical fragin and ^15^N-labelled fragin could be observed by high resolution mass spectrometry (HRMS), in addition to a characteristic loss of ^15^NO for ^15^N-labelled fragin detected by HRMS/HRMS (**Fig 4D**).

**Fig 4.**
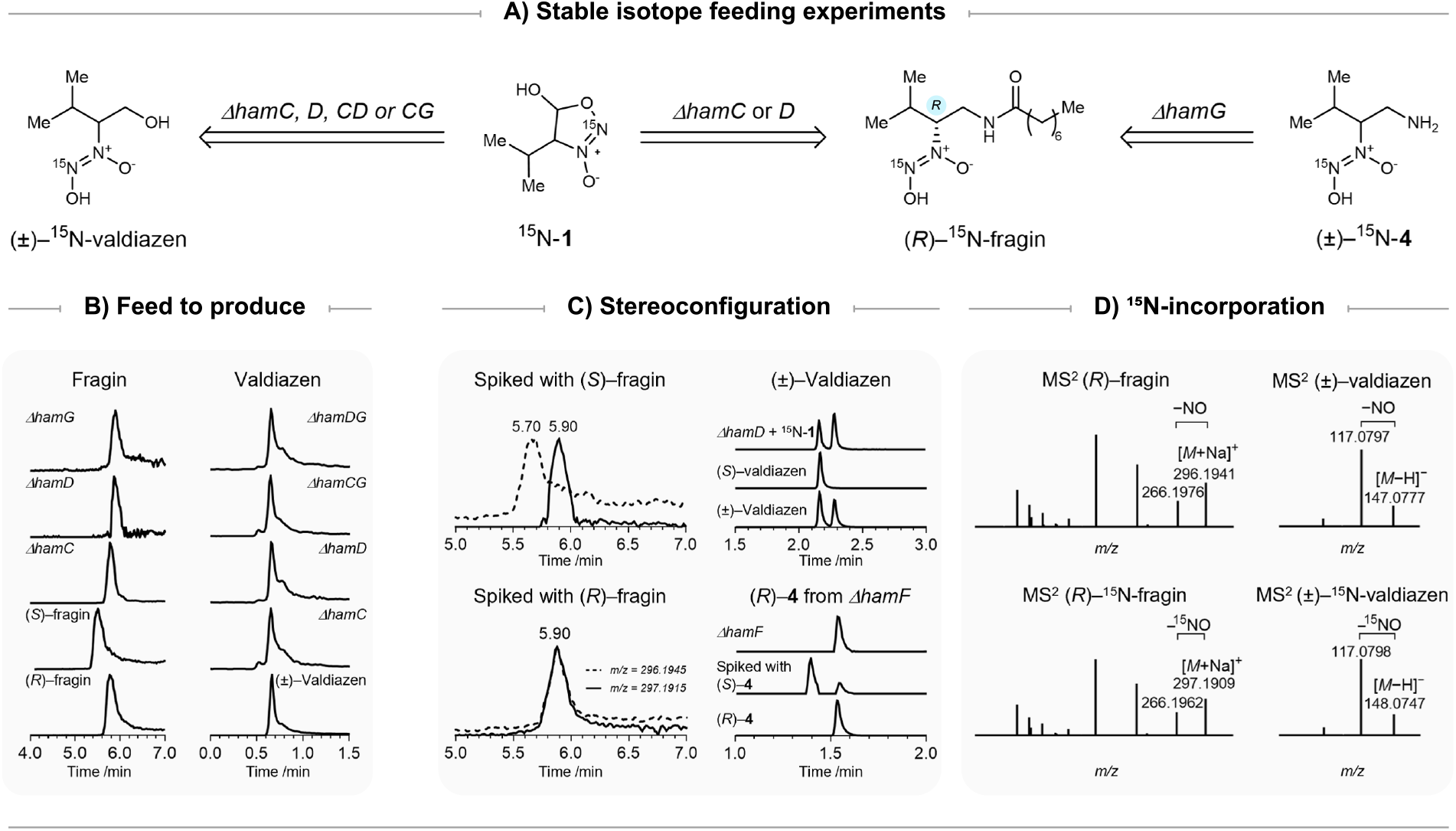
**A)** Dihydrosydnone *N*-oxide **1** and β-amino diazeniumdiolate **4** are competent biosynthetic intermediates for the production of fragin and valdiazen. **B) Feeding experiments**: Extracted-ion chromatogram (EIC) traces showed ^15^N-valdiazen production and the stereoselective (*R*)–^15^N-fragin production by feeding ^15^N-labelled intermediates in mutants related to *hamC, hamD*, and *hamG*. **C) Configuration**: The configuration of fragin produced in feeding experiments was confirmed by spiking with unlabelled (*R*)– or (*S*)–fragin. One example set of EIC traces from *ΔhamD* feeding experiments is shown here. FDAA-derivatisation and spiking experiments confirmed the configuration of valdiazen and compound **4. D)** ^15^**N-incorporation**: The incorporation of ^15^N-labels was confirmed by HRMS/HRMS spectra.

Similar feeding experiments were conducted to deconvolute valdiazen biosynthesis. Due to the detection challenges of valdiazen, each supernatant sample was extracted before the analysis by UHPLC-HRMS in negative mode using an NH_4_HCO_3_ buffer to increase sensitivity. As expected, supplying ^15^N-**4** to *ΔhamG* mutant did not give any valdiazen (**Supplementary Fig 3**). To our surprise, valdiazen was produced in all mutants tested when fed with dihydrosydnone *N*-oxide ^15^N-**1** (**Fig 4B**). The characteristic loss of ^15^NO on valdiazen can also be seen in HRMS/HRMS in negative mode (**Fig 4D**). We then extended the feeding experiments to the remaining genes of the *ham* cluster (**Supplementary Fig 4** and **Supplementary Table 2**). When providing dihydrosydnone *N*-oxide ^15^N-**1**, *ΔhamA, ΔhamB*, and *ΔhamE* mutants were all capable of synthesising (*R*)–^15^N-fragin and ^15^N-valdiazen. Notably, after spiking with analytical (*R*)–fragin, the amount of endogenous unlabelled fragin in *ΔhamB* was so high that the signal of its naturally occurring ^13^C-isomer ([*M*+Na]^+^ = 297.1978) dominated the signal of the ^15^N-labelled fragin ([*M*+Na]^+^ = 297.1915). Nevertheless, spiking with analytical (*S*)–fragin indirectly confirmed the (*R*)–configuration of the labelled fragin produced by *ΔhamB* (**Supplementary Fig 2**). Although HamB was previously found to be essential for the antifungal activity of *B. cenocepacia* H111,^11^ we detected the presence of natural fragin in *ΔhamB* mutant by UHPLC-HRMS. We hypothesised that the production of fragin was much lower in *ΔhamB* compared to the wild type, which led to the low antifungal activity in a disc-diffusion assay.^11^ The presence of valdiazen in all strains was also puzzling since it was postulated that the reductase domain of NRPS HamD was responsible for the two-electron reduction at the last biosynthetic step.^21,41–43^ These observations led to the hypothesis that the reduction of aldehyde/hemiacetal to the alcohol is mediated by an enzyme outside the *ham* cluster.

### Enantioseparation of valdiazen and β-amino diazeniumdiolate 4

Valdiazen was challenging to separate on chiral stationary phases with conventional UHPLC methods, due to its highly polar, volatile, and sensitive nature. Similar challenges were encountered with amine analogue **4**, as a valid biosynthetic intermediate for fragin production. To examine the configuration of valdiazen and compound **4** in supernatants, Marfey’s reagent (FDAA) was chosen to derivatise the diazeniumdiolate before UHPLC. FDAA is a well-known reagent that reacts with primary amines via nucleophilic aromatic substitution allowing enantioseparation of optical isomers such as amino acids.^17,44^ We speculate that diazeniumdiolate is nucleophilic towards alkyl and aryl halides under mild basic conditions.^26,45^ To our delight, pre-column derivatisation with FDAA enabled chiral separation of valdiazen and compound **4**. Consistent with the results from our previous study,^11 15^N-valdiazen derived from the feeding experiments was indeed produced as a racemate (**Fig 4C**). FDAA was also capable of capturing unlabelled **4** in the supernatants from *ΔhamF* cultures without exogenous isotopically labelled tracers. By spiking with FDAA-derivatised enantiomeric pure (*R*) and (*S*)–**4**, we demonstrated that only (*R*)–**3** was produced *in vivo* (**Fig 4C**).

### Homochirality in quorum sensing signalling

We tested the efficacies of valdiazen and β-amino diazeniumdiolate **4** to serve as a signal molecule by measuring β-galactosidase activities of a transcriptional fusion of the *hamA* promoter to *lacZ* in a *ΔhamD* mutant (**Fig 5A**). Both metabolites induced the *hamA* promotor, with valdiazen showing generally higher activity compared to compound **4**. To our surprise, the (*S*)–enantiomers of both metabolites showed significantly higher β-galactosidase activities than the (*R*)–enantiomers. In the case of compound **4**, the observed discrepancy in activity could be explained by the fact that HamF directs (*R*)–**4** to the biosynthesis of (*R*)–fragin in a *ΔhamD* mutant. Feeding valdiazen to a *ΔhamD* mutant did not restore fragin production, indicating that valdiazen does not serve as a direct precursor in the biosynthesis of fragin (**Fig 5B**). Analysis of the FDAA-derivatised cell-free supernatants showed no racemisation of the supplied valdiazen (**Fig 5B**). However, at present we cannot rule out the possibility that the transporter required for the uptake of valdiazen prefers (*S*)– over the (*R*)–enantiomer. In any case, our results show that (*S*)–valdiazen is more efficient to induce transcription of the *hamA* promoter than (*R*)–valdiazen.

**Fig 5.**
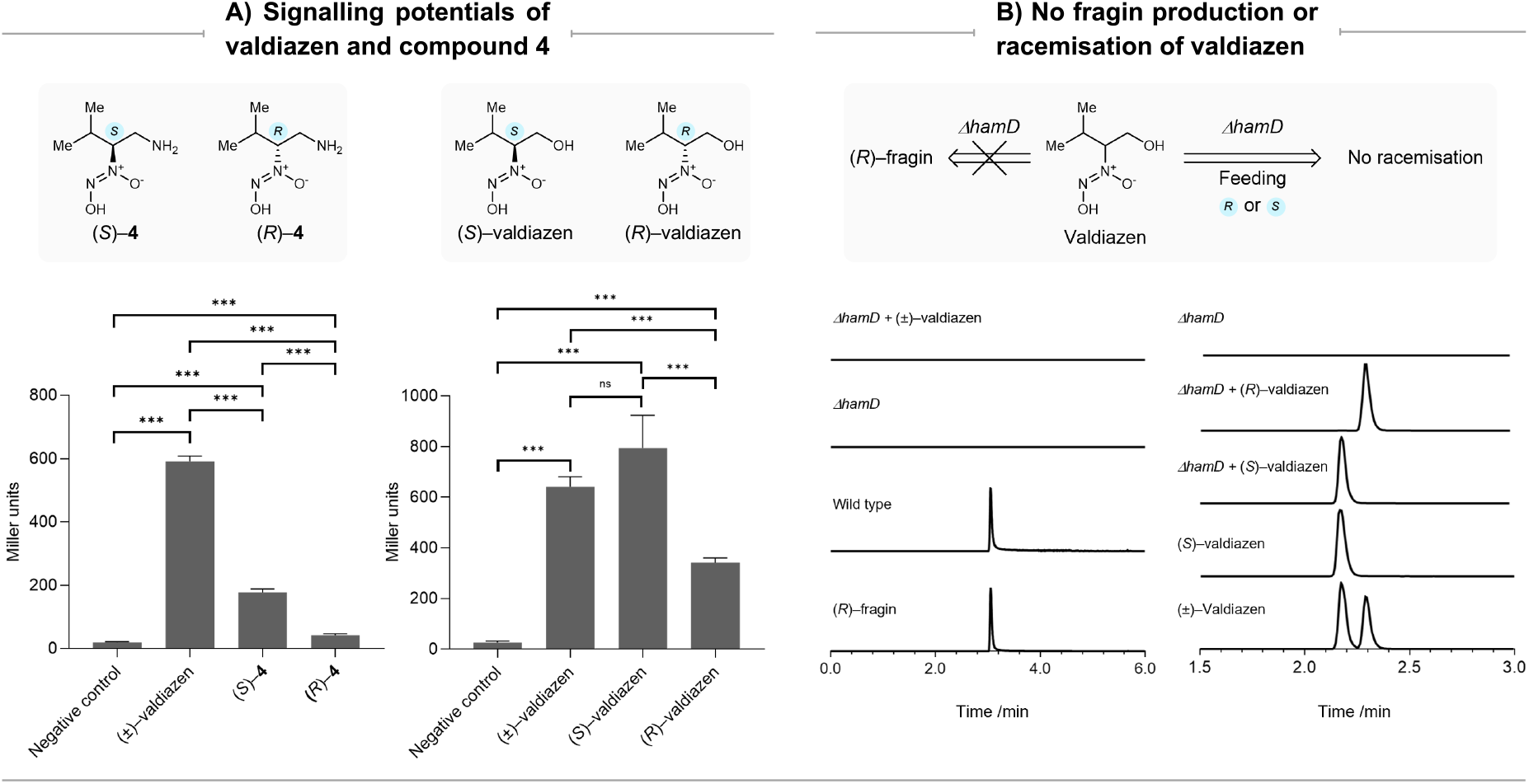
**A)** Valdiazen and β-amino diazeniumdiolate **4** showed distinct signalling potentials in inducing the *hamABCDE* operon promoter in a β-galactosidase assay. **B)** EIC traces confirmed that valdiazen is not a direct biosynthetic precursor in a *ΔhamD* mutant: The left set of EIC traces confirmed that no fragin was produced when feeding (±)–valdiazen to a *ΔhamD* mutant. The right set of EIC traces showed that valdiazen enantiomers fed in a *ΔhamD* mutant did not racemise.

## Discussion

In summary, our study provided evidence that *B. cenocepacia* H111 leveraged keto-enol tautomerisation and enzyme-mediated dynamic kinetic resolution to produce (±)–valdiazen and (*R*)– fragin via pathways encoded by the *ham* gene cluster. By using synthetic chemistry and gene disruption, we deconvoluted the late steps of (*R*)–fragin and (±)–valdiazen biosynthesis and suggested a mechanism of stereodiversification via the intermediate dihydrosydnone *N-*oxide **1**. The dihydrosydnone *N-*oxide **1** and β-amino diazeniumdiolate **4** were identified as biosynthetically relevant intermediates in the production of (*R*)–fragin and (±)–valdiazen. Our results also suggest that the racemisation during the biosynthesis was not mediated by racemases or epimerases, but rather via keto-enol tautomerisation. The racemic intermediate then underwent enzyme-mediated dynamic kinetic resolution, where only the (*R*)–enantiomer can be recognised by HamG and reductively aminated to give (*R*)–**4**. Finally, acylation of (*R*)–**4** by HamF results in (*R*)–fragin. HamF was also capable of mediating enzymatic kinetic resolution when providing racemic **4**. Further studies are required to investigate the molecular interactions between the enzymes and their respective substrates.

Research of the past three decades provided overwhelming evidence that most bacteria are capable of communicating with each other by the aid of small signal molecules. In most cases, these signalling systems are used to monitor the density of the population to control social behaviours, a phenomenon commonly referred as quorum sensing (QS).^46,47^ *N*-acyl homoserine lactones (AHL) is a QS signal found in Gram-negative bacteria.^48^ Both L– and D–AHLs have been identified, with D–AHLs being generally much less efficient in inducing the quorum-sensing responses.^49–51^ Similar to classic AHL-based QS systems, we discovered that both stereoisomers of valdiazen were synthesised, although (*S*)–valdiazen displayed significantly higher activity than its enantiomeric counterpart. We were intrigued that the signal molecules were not produced in a stereospecific manner, given the superior biological activity of one of the enantiomers. We hypothesised that the less potent stereoisomer of the autoinducer could in fact act as a partial agonist.

This study utilised both chemical synthesis and biosynthesis experiments in wild type and mutant bacteria. By synthetic chemistry, tool molecules were developed to tackle intricate biosynthetic questions. In the meantime, the pursue of biological questions led to the discovery of a new chemical scaffold, dihydrosydnone *N*-oxide. This unique heterocycle may serve as a new NO-releasing motif. Moreover, NRPS clusters with a C-terminal reductase have also been identified in other bacteria, such as *Mycobacterium tuberculosis* and *Bacillus brevis*.^42,52–54^ Our study may therefore serve as a blueprint for the identification of intermediates that represent branching points for the biosynthesis of bioactive compounds with different configuration.

## Methods

### Bacterial strains and plasmids used in this study

Strains and plasmids used in this study are listed in **Supplementary Table 6**.

### Construction of *Burkholderia cenocepacia* H111 deletion mutants

Construction of H111 strains with single gene deletions in *hamC* and *hamD* has been described in our previous study.^11^

Single deletion mutant H111 *ΔhamG* and double deletion mutants H111 *ΔhamCG* and H111 *ΔhamDG* were constructed using the pGPI-SceI/pDAI system.^55^ Briefly, the upstream and downstream homology regions flanking the region to be deleted (*hamC* and *hamD* genes) were cloned into pGPI-SceI::TetAR and the plasmid is transferred to the *hamG* mutant of H111. Correct integration of the plasmid into the host genome of *hamG* mutant was verified by PCR. The plasmid pDAI::Gm^R^, which carries the I-SceI nuclease, was transferred into the target strain, which resulted in a double-strand DNA break at its recognition site, linearising the chromosome and resulting in a second homologous recombination event. Conjugants were selected on PIA Gm plates and screened μM by PCR using the check primers. Colonies were patched on PIA and PIA Gm20 to isolate colonies that are cured of the pDAI.

### Stable isotope feeding experiments

The composition of ABG minimal medium has been described in our previous study.^11^

Different mutants of H111 were grown in ABG minimal medium containing 0.005 % yeast extract with agitation (220 rpm) at 37 °C for 24 hours, after which 50 μM ^15^N–intermediate was added to the cultures and subsequently incubated for 48 hours. Bacterial cultures were centrifuged at 5000 rpm with an Eppendorf Centrifuge 5804R and supernatants were filtered with a Millipore Steritop Vacuum Bottle Top Filter 0.22 μm system to remove all the bacterial cells. The ^15^N–intermediates used in this study include ^15^N–dihydrosydnone *N-*oxide **1**, and ^15^N–β-amino diazeniumdiolate analogue **4**.

### Assessment of promotor activity in liquid culture

Construction of the transcriptional lacZ fusions with the putative promoters of the *hamABCDE* operon has been previously described.^11^

Promoter activity of transcriptional lacZ fusions by β-galactosidase assays was performed as previously described with minor modifications.^56^ Briefly, bacterial cells were grown overnight in ABG minimal medium and synthetic valdiazen or β-amino diazeniumdiolate **4** dissolved in methanol was added to the cultures, where indicated. Bacterial cells were harvested, resuspended in 2 mL Z-buffer and 1 mL bacterial suspension was used to measure the optical density (OD_600_). Cells were lysed by adding chloroform (25 μL) and sodium dodecyl sulphate (SDS; 25 μL, 0.01%) to the residual 1 mL of bacterial suspension and briefly vortexed. The resulting mixture was incubated for 10 min at 28 °C. Subsequently, the reaction was initiated by adding *o*-nitrophenyl-β-galactosidase (ONPG; 200 μL, 4 mg/mL), vortexed briefly and incubated at room temperature. The reaction was stopped by the addition of 500 μL 1 M aqueous Na_2_CO_3_. Cell debris was removed by centrifugation (10 min, 15000 rpm) and the absorbance at 420 nm (OD_420_) was measured. The β-galactosidase activity was calculated using the following equation: Activity [Miller units] = (1000 × OD_420_) / (time [min] × volume [mL] × OD_420_), where time is the incubation time measured in minutes and volume is the number of cells used in mL.

### Chiral HPLC separation and detection of fragin

The supernatants were transferred in Eppendorf tubes, centrifuged (5 min, 20238 rcf) and part of the liquid was transferred in UHPLC vials. The analysis was performed on a Vanquish Horizon UHPLC system. The separation was achieved using a Lux^®^ i-Amylose-3 column (*Phenomenex*, 50 × 2.0 mm, 3 μm) with a flow at 0.3 mL/min, a temperature of 45 °C, an injection volume of 3 μL, and a solvent system composed of A (H_2_O + 0.1% HCO_2_H) and B (MeOH + 0.1% HCO_2_H). The column was equilibrated for 0.5 min at 40% B, the gradient of the run started with isocratic conditions with 40% B over 0.5 min, increased to 90% B in 6.5 min, followed by an increase to 100% B in 1 min, 100% B was held for 1 min, and the gradient decreased to 40% B in 0.4 min, the final conditions were held for 0.6 min. For the results obtained from *ΔhamC* mutants, the supernatant was concentrated, and dissolved in MeOH before analysis.

The ion source parameters of the HRMS (Exploris 240, *ThermoFischer*) in positive mode were set as follows: spray voltage 3.5 kV; capillary temperature 325 °C; sheath gas 50 L min^−1^; aux gas 10 L min^− 1^; sweep gas 1 L min^−1^; and vaporizer temperature 340 °C. Full scan analysis was set up at a resolution of 60000 with a scan range of 100–1500 *m*/*z* in positive mode and a ddMS_2_ was set up with a resolution of 30000, a scan width of 1 Da, a HCD collision energy at 10,30, and 50% and was triggered for the targeted masses at 296.1945 *m*/*z* for [fragin + Na]^+^ and 297.1915 *m*/*z* for [^15^N-fragin + Na]^+^. The EASY-IC™ internal calibration system was used at the beginning of each run. (*R*) or (*S*)–fragin were identified by comparison of their retention times with an analytical standard and by analysis of their MS/MS spectra. To further confirm the configuration of fragin, spiking experiment was performed by adding a solution of the (*R*) and (*S*)–fragin to each supernatant respectively before UHPLC separation. The separation and detection procedures were performed for *ΔhamA, B, C, D, E, CG*, and *DG* fed with ^15^N-**1** and with *ΔhamG* fed with ^15^N-**4**. Experiments involving supernatants were repeated in triplicates using the biological triplicates generated from the respective feeding experiments. All samples were centrifuged (5 min, 20238 rcf) and transferred in HPLC vials before analysis.

### Achiral HPLC separation and detection of fragin

To improve the sensitivity of the detection, a more sensitive method to detect fragin was developed using an achiral column. Briefly, the supernatants were transferred in Eppendorf tubes, centrifuged (5 min, 20238 rcf) and part of the liquid was transferred in UHPLC vials. The analysis was performed on a Vanquish Horizon UHPLC system. The separation was achieved using an EVO C18 column (*Phenomenex*, 50 × 2.1 mm, 1.7 μm) with a flow at 0.4 mL/min, a temperature of 40 °C, an injection volume of 1 μL, and a solvent system composed of A (H_2_O + 0.1% HCO_2_H) and B (MeCN + 0.1% HCO_2_H). The column was equilibrated for 0.5 min at 5% B, the gradient of the run started with isocratic conditions with 5% B over 0.5 min, increased to 95% B in 3.5 min, and was kept at 95% B for 1 min. The ion source parameters of the HRMS (Exploris 240, *ThermoFischer*) in positive mode were set as follows: spray voltage 3.5 kV; capillary temperature 325 °C; sheath gas 50 L min^−1^; aux gas 10 L min^− 1^; sweep gas 1 L min^−1^; and vaporizer temperature 340 °C. Full scan analysis was set up at a resolution of 60000 with a scan range of 100–1500 *m*/*z* in positive mode and a ddMS_2_ was set up with a resolution of 30000, a scan width of 1 Da, a HCD collision energy at 10, 30, and 45% and was triggered for the targeted masses at 244.2145 *m*/*z* for [fragin – NO + H]^+^ and 274.2125 *m*/*z* for [fragin + H]^+^. The EASY-IC™ internal calibration system was used at the beginning of each run. Fragin was identified by comparison of the retention time of the analytical standard (3.05 min) and by analysis of its MS/MS spectra. The separation and detection procedures were performed for WT, *ΔhamD*, and *ΔhamD* fed with (±)–valdiazen. Experiments involving supernatants were repeated in triplicates using the biological triplicates generated from the respective experiments.

### Achiral HPLC separation and detection of valdiazen

For valdiazen extraction, cell-free supernatant was adjusted to a pH value of 11 with 10 M aq. NaOH and extracted twice with 0.5 volumes dichloromethane. The dichloromethane phases were discarded, and the water phase was subsequently adjusted to pH value of 5 with 10 M aq. HCl and extracted twice with 0.5 volumes of dichloromethane. The two dichloromethane phases were combined, dried using anhydrous magnesium sulphate (Sigma-Aldrich, Switzerland), filtered, and concentrated *in vacuo*. The concentrated extracts were stored at −20 °C. The extracts were dissolved in MeOH (1 mL), transferred in Eppendorf tubes, centrifuged (5 min, 20238 rcf) and part of the liquid was transferred in UHPLC vials.

The analysis was performed on a Vanquish Horizon UHPLC system. The separation was achieved using a Luna^®^ Omega PS C18 column (*Phenomenex*, 50 × 2.1 mm, 1.6 μm) with an injection volume of 1 μL and a solvent system composed of A (H_2_O + 0.5 mM NH_4_CO_3_) and B (MeCN). Two analytical methods (A and B) were used and, for both methods, the column was equilibrated for 0.5 min at the starting conditions. **Method A**: The run started with a flow of 0.3 mL/min and isocratic conditions were kept at 0% B over 0.49 min, the flow was increased to 0.4 mL/min over 0.01 min, the gradient increased from 0 to 95% B in 3 min, 95% B was held for 2 min, the gradient decreased to 0% B in 0.1 min, and the final conditions were held for 1 min. **Method B**: The flow was 0.4 mL/min and the gradient of the run started with an increase from 0 to 95% B in 3.5 min, 95% B was held for 2 min, the gradient decreased to 0% B in 0.1 min, and the final conditions were held for 1 min. With method A, valdiazen was detected at 0.67 min and with the method B valdiazen was detected at 0.53 min. The ion source parameters of the HRMS (Exploris 240, *ThermoFischer*) in negative mode were set as follows: spray voltage 2.5 kV; capillary temperature 325 °C; sheath gas 50 L min^−1^; aux gas 10 L min^−1^; sweep gas 1 L min^−1^; and vaporizer temperature 325 °C. Full scan analysis was set up at a resolution of 60000 with a scan range of 100–1500 *m*/*z* in negative mode and a ddMS_2_ was set up with a resolution of 30000, a scan width of 0.4 Da, a HCD collision energy at 10, 30, and 50% and was triggered for the targeted masses at 147.0775 *m*/*z* for [valdiazen – H]^−^ and 148.0745 *m*/*z* for [^15^N-valdiazen − H]^−^. The EASY-IC™ internal calibration system was used at the beginning of each run. Valdiazen was identified by comparison of the retention time of the analytical standard and by analysis of its MS/MS spectra. The separation and detection procedures were performed for *ΔhamA, B, C, D, E, CG*, and *DG* fed with ^15^N-**1**. Some samples were diluted in H_2_O before analysis due to the high concentration of the targeted compounds. Experiments involving supernatants were repeated in triplicates using the biological triplicates generated from the respective feeding experiments. All samples were centrifuged (5 min, 20238 rcf) and transferred in HPLC vials before analysis.

### Chiral HPLC separation and detection of valdiazen via FDAA pre-column derivatisation

An aliquot (10 mL) of the supernatant from the cultures was concentrated under a gentle flow of nitrogen for 16 hours at 40 °C. The residue was resuspended in MiliQ water (100 μL), treated with 1% FDAA in acetone (200 μL), 1 M aq. sodium bicarbonate solution (40 μL) and heated at 40 °C for 1 hour. The mixture was cooled to r.t. and treated with 2 M aq. HCl (20 μL). The sample was then concentrated under a gentle flow of nitrogen for 1 hour to remove acetone. The derivatisation was performed using the analytical standards (*S*)–valdiazen and (±)–valdiazen, and with the supernatants (10 mL) of the *ΔhamD* fed with ^15^N-**1**. Experiments involving supernatants were repeated in triplicates using the biological triplicates generated from the respective feeding experiments. For spiking experiments, samples from supernatants were mixed with FDAA-treated synthetic analytical samples. All samples were centrifuged (5 min, 20238 rcf) and transferred in HPLC vials before analysis.

The analysis was performed on a Vanquish Horizon UHPLC system. The separation was achieved using an EVO C18 column (*Phenomenex*, 50 × 2.1 mm, 1.7 μm) with a flow at 0.4 mL/min, a temperature of 40 °C, an injection volume of 1 μL, and a solvent system composed of A (H_2_O + 0.5 mM NH_4_CO_3_) and B (MeCN). The column was equilibrated for 0.5 min at 5% B, the gradient of the run started to increase from 5% to 95% B over 4 min, and was kept at 95% B for 2 min.

The ion source parameters of the HRMS (Exploris 240, *ThermoFischer*) in negative mode were set as follows: spray voltage 2.5 kV; capillary temperature 325 °C; sheath gas 50 L min^−1^; aux gas 10 L min^− 1^; sweep gas 1 L min^−1^; and vaporizer temperature 340 °C. The EASY-IC™ internal calibration system was used at the beginning of each run. A full scan with a resolution of 60000, a scan range of 140–1000 Da, and RF lens of 70% were used in negative mode and a ddMS_2_ was set up with a scan width of 0.4 *m*/*z*, at a resolution of 30000, and with an HCD collision energy at 15, 30, and 50%). Valdiazen-FDAA derivatives (*m*/*z* 399.12699 for [valdiazen-FDAA – H]^-^ and *m*/*z* 400.12402 for [^15^N-valdiazen-FDAA – H]^-^) were identified by comparison of the retention time of the analytical standard (2.17 min for *S* and 2.28 min for *R*) and by analysis of their MS/MS spectra. The separation and detection procedures were performed for *ΔhamD* fed with ^15^N-**1**. Experiments involving supernatants were repeated in triplicates.

### Chiral HPLC separation and detection of the amine analogue 4 via pre-column derivatisation with FDAA

An aliquot (10 mL) of the supernatant from *ΔhamF* culture was concentrated under a gentle flow of nitrogen for 16 hours at 40 °C. The residue was resuspended in MiliQ water (100 μL), treated with 1% FDAA in acetone (200 μL), 1 M aq. sodium bicarbonate solution (40 μL) and heated at 40 °C for 1 hour. The mixture was cooled to r.t. and treated with 2 M aq. HCl (20 μL). The sample was then concentrated under a gentle flow of nitrogen for 1 hour to remove acetone. The same condition was used for synthetic enantiomeric pure amine analogues **4** (*R* and *S*) at 5 μmol scale which were used as analytical standards and spiking experiments later. The sample was then diluted with MeCN up to a volume of 1 mL, vortexed and further diluted 2000 times. For spiking experiments, samples from supernatants were mixed with FDAA-treated synthetic analytical samples. All samples were centrifuged (5 min, 20238 rcf) and transferred in HPLC vials before analysis. Experiments involving supernatants were repeated in triplicates using the biological triplicates generated from the respective feeding experiments.

The analysis was performed on a Vanquish Horizon UHPLC system. The separation was achieved using an EVO C18 column (*Phenomenex*, 50 × 2.1 mm, 1.7 μm) with a flow at 0.4 mL/min, a temperature of 30 °C, an injection volume of 3 μL, and a solvent system composed of A (H_2_O + 0.1% HCOOH) and B (MeCN + 0.1% HCOOH). The column was equilibrated for 0.5 min at 5% B, the gradient of the run started to increase from 5% to 95% B over 4 min, and was kept at 95% B for 2 min. The ion source parameters of the HRMS (Exploris 240, *ThermoFischer*) in positive mode were set as follows: spray voltage 3.5 kV; capillary temperature 325 °C; sheath gas 50 L min^−1^; aux gas 10 L min^−1^; sweep gas 1 L min^−1^; and vaporizer temperature 340 °C. The EASY-IC™ internal calibration system was used at the beginning of each run. A full scan with a resolution of 240000, a scan range of 400.1–400.2 Da, and RF lens of 70% were used in negative. Amine **4** FDAA derivatives (*m*/*z* 400.15752 for [amine **4**-FDAA + H]^+^) were identified by comparison of the retention time of derivatized analytical standards (1.37 for *S* and 1.54 min for *R*). For an optimal detection of the peak, the mass tolerance was set up to 3 ppm. The separation and detection procedures were performed for *ΔhamF*. Experiments involving supernatants were repeated in triplicates.

### Data analysis and plot generation

The UHPLC-HRMS data were visualized with FreeStyle (*ThermoFischer*), and the chromatograms and spectra were exported as CSV files. GraphPad Prism (version 10.1.2) was used to plot the chromatographic data points in the relevant range of retention time from a single feeding experiment. The intensity data from the chromatograms were normalised with the highest intensity set to 100% before being plotted as a line graph. Overlaid plots comparing negative control and positive results were plotted using intensity without normalisation. MS/MS graph were plotted directly from source data. Student’s t-tests were performed using original algorithm package from GraphPad Prism (version 10.1.2).

## Acknowledgment

We thank Prof. Dr. Roderich Süssmuth for helpful discussion on bacterial biosynthesis. We thank the Swiss National Science Foundation (Grant No. 186410) for financial support.

## Competing interests

The authors declare no competing interests.

## Author Contributions

K.G., L.E. and S.S. conceived and supervised the project. J.R. designed the major synthetic routes and J.R., T.K. and S.S. synthesised the compounds involved in this study. A.M. designed biological assays and performed most of the assays. M.G. performed stable isotope feeding experiments. S.S. designed and performed analytical experiments related to mass spectrometry analysis. J.R. performed precolumn derivatisation. O.B. and A.L. performed X-ray single crystal experiments and the analysis of these data. J.R., A.M., T.K., O.S.S, L.E., and K.G., analysed the data. J.R. wrote the manuscript with substantial contributions from A.M., S.S., K.G, and L.E. All authors revised the manuscript.

## Notes

### Competing Interest Statement

The authors have declared no competing interest.

