## Supporting information for "Functional Biosynthetic Stereodivergence in a Gene Cluster via a Dihydrosydnone *N*-oxide"

### General procedure

The synthesis of (*S*)–fragin, (*R*)–fragin, (*S*)–valdiazen, (*R*)–valdiazen were performed as described in our previous publication.[1]

Unless otherwise stated, all chemicals were of reagent grade and purchased from Sigma-Aldrich-Merck, Acros Organics, Honeywell, Fluorochem or TRC. Solvents for reactions were of analytical grade. Evaporation of solvents *in vacuo* was performed with a rotary evaporator equipped with a water bath at 40 °C and indicated pressure. **Thin layer chromatography** (TLC): reaction control with TLC were performed on Merck TLC plates, silica gel 60 F254 on aluminium with the indicated solvent system; the spots were visualized by UV light (254 nm) or $KMnO_4$ stain. **Flash Chromatography**: Silica gel column chromatography was performed using silica gel 60 (230–400 Mesh) purchased from Sigma-Aldrich and the compounds were eluted with the solvent mixture indicated. **Solid-phase extraction** (SPE) **column**: Discovery® DSC-18 SPE tube for semi-purification were purchased from Sigma-Aldrich-Merck, and used following their guidelines; **Ultra-high-performance liquid chromatography coupled to mass spectrometry** (UHPLC-MS) for **reaction control**: reaction controls with UHPLC-MS were performed on *Ultimate 3000 LC instrument* (*Thermo Fisher Scientific*) coupled to a triple quadrupole *Quantum Ultra EMR MS* (*Thermo Fisher Scientific*) using a reversed-phase column (*Kinetex®* EVO C18; 1.7 µm; 100 Å, 50 × 2.1 mm; *Phenomenex*), heated to 35 °C. The LC was equipped with an *HPG-3400RS* pump, a *WPS-3000TRS* autosampler, a *TCC-3000RS* column oven and a *Vanquish DAD* detector (all *Thermo Fisher Scientific*). The following solvents were used as eluents: $H_2O$+0.1 % $HCO_2H$ (A), MeCN+0.1 % $HCO_2H$ (B). The MS was equipped with an H-ESI II ion source. The source temperature was 250 °C, the capillary temperature 270 °C and capillary voltage 3500 V, and datasets were acquired at resolution 0.7 on Q3 in centroid mode. **Infrared spectra** (IR): IR spectra were recorded on *SpectrumTwo FT-IR Spectrometer* (*Perkin–Elmer*) equipped with a *Specac Golden GateTM ATR* (attenuated total reflection) accessory. The samples were applied as neat samples or as films. **Melting points** (m.p.): Melting points were determined using the *Büchi* B-545 apparatus in open capillaries and are uncorrected. **Nuclear magnetic resonance spectra** (NMR): $^{1}$H-NMR spectra were recorded using the indicated deuterated solvents at 298 K on the instruments *AVII* or *III-500* (500 MHz with Cryo-BBO, TXI, BBI or BBO probe) or *AVII-400* (400 MHz with QNP, BBO or BBFO probe); $^{13}$C-NMR spectra were recorded in the indicated solvents and on the same instruments. **High resolution electrospray ionization mass spectrometry for compound characterization** (HR-ESI-MS): HRMS for the characterization of synthetic compounds were measured on *Dionex Ultimate 3000* UHPLC system (*ThermoFischer Scientifics*, Germering, Germany) connected to a QExactive MS with a heated ESI source (*ThermoFisher Scientific*, Bremen, Germany); onflow injection of 1 µL sample ($c$ = ca. 50 µg $mL^{-1}$ in the indicated solvent) with an *XRS* auto-sampler (*CTC*, Zwingen, Switzerland); flow rate 120 µL $min^{-1}$; ESI: spray voltage 3.0 kV, capillary temperature 280 °C, sheath gas 30 L $min^{-1}$, aux gas 8 L $min^{-1}$, s-lens RF level 55.0, aux gas temperature 250 °C ($N_2$); full scan MS in the alternating (+)/(−)-ESI mode; mass ranges 80–1200 *m*/*z*, 133–2000 *m*/*z*, or 200–3000 *m*/*z* at 70000 resolution (full width half-maximum); automatic gain control (AGC) target of $3.00\times10^{6}$; maximum allowed ion transfer time (IT) 30 ms; mass calibration to <2 ppm accuracy with *Pierce®* ESI calibration solns. (*ThermoFisher Scientific,* Rockford, USA); lock masses: ubiquitous erucamide (*m*/*z* 338.34174, (+)-ESI) and palmitic acid (*m*/*z* 255.23295, (–)-ESI). **Ultra-high-performance liquid chromatography coupled to High resolution electrospray ionization mass spectrometry for the detection of fragin and valdiazen** (UHPLC-HR-ESI-MS): The samples were measures on a Vanquish Horizon UHPLC system (*ThermoFischer*) equipped with a quaternary pump, an autosampler, a Diode Array Detector, a Split Sampler HT, a Binary Pump H, and Column Compartment. The UHPLC system used was connected to a HRMS (Exploris 240, *ThermoFischer*) instrument. **Specific optical rotation**: Specific optical rotations were recorded by *Jasco P-2000 Polarimeter* with a path length of 1 dm using the 589.3 nm D-line of sodium. Measurements were recorded at the indicated temperature (in °C) and concentration (in g/100 mL) in the indicated solvent. **Preparative high performance liquid chromatography (preparative HPLC)**: Purification was made using prominence modular HPLC instrument (*Shimadzu*) coupled to an *SPD-20A* UV/Vis detector (*Shimadzu*) with a reversed-phase column (*Phenomenex Synergi™* 10 µm Hydro-RP 80 Å, 250 mm × 21.2 mm). The LC was equipped with a *CBM-20A* system controller, *LC-20A* solvent delivery unit, a *DGU-20A* degassing unit, *FRC-10A* fraction collector (all units from *Shimadzu*). The conditions used were indicated in the experimentals.

### Single crystal X-ray diffraction

The measurements were made at 160 K on a *Rigaku Oxford Diffraction Synergy/Hypix* diffractometer (**1**, **4**) and on a *Rigaku Oxford Diffraction SuperNova/Atlas* area detector diffractometer (**17**) using Cu radiation ($\lambda$ = 1.54184 Å) and *Oxford Instruments Cryojet XL* coolers. The selected suitable single crystals were mounted in oil on cryo-loops. Pre-experiment, data collection, data reduction and absorption correction were performed with the program suite *CrysAlisPro*.[2] Using *Olex2*,[3] the structure were solved with the *SHELXT*[4] small molecule structure solution program and refined with the *SHELXL* program package[5] by full-matrix least-squares minimization on $F^2$. *PLATON*[6] was used to check the results of the X-ray analyses. CCDC 2351242 (**1**), 2351243 (**4**) and 2351244 (**17**) contain the supplementary crystallographic data for this paper. These data can be obtained free of charge via www.ccdc.cam.ac.uk/data_request/cif, or by emailing, or by contacting The Cambridge Crystallographic Data Centre, 12 Union Road, Cambridge CB2 1EZ, UK; fax: +44 1223 336033.

**Supplementary Table 1.** Comparison between bond lengths (in Å) from known crystal structures of dihydrosydnone *N*-oxide and its resemblants.[7–10]

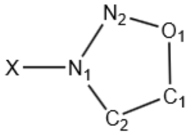


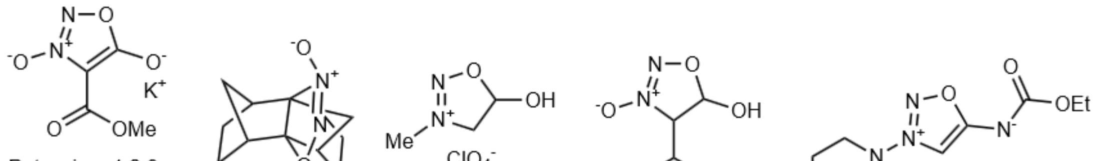

| | Potassium 1,2,3-oxadiazole-3-oxido-4-methylcarboxylate-5-hydroxylate | 4,5-dihydro-1,2,3-oxadiazole-3-oxide | 5-hydroxy-3-methyloxadiazolinium | Dihydrosydnone *N*-oxide | Molsidomine |
|---|---|---|---|---|---|
| X-$N_1$ | 1.262(3) | 1.257(8) | 1.459 | 1.2730(10) | 1.375 |
| $N_1$-$N_2$ | 1.324(3) | 1.260(9) | 1.238 | 1.2661(13) | 1.301 |
| $N_2$-$O_1$ | 1.386(3) | 1.408(8) | 1.319 | 1.3835(12) | 1.369 |
| $O_1$-$C_1$ | 1.367(3) | 1.488(7) | 1.515 | 1.4755(12) | 1.372 |
| $C_1$-$C_2$ | 1.405(3) | 1.528(7) | 1.496 | 1.5148(13) | 1.377 |
| $C_2$-$N_1$ | 1.380(3) | 1.501(7) | 1.464 | 1.4869(11) | 1.348 |

**Supplementary Table 2.** Summary of feeding experiments using $^{15}$N-labelled intermediates in various mutants.

| | Production of $^{15}$N-labelled compound | | Average retention time±standard error (min) | |
|---|---|---|---|---|
| | (±)–valdiazen | (*R*)–fragin | (±)–valdiazen | (*R*)–fragin |
| *ΔhamA* + $^{15}$N-**1** | Yes | Yes | 0.509±0.0284 | 6.02±0.0153 |
| *ΔhamB* + $^{15}$N-**1** | Yes | Yes | 0.497±0.0300 | 6.02±0.0115 |
| *ΔhamC* + $^{15}$N-**1** | Yes | Yes | 0.659±0.00808 | 5.76±0.0404 |
| *ΔhamD* + $^{15}$N-**1** | Yes | Yes | 0.654±0.00173 | 5.88±0.0404 |
| *ΔhamE* + $^{15}$N-**1** | Yes | Yes | 0.483±0.00808 | 6.02±0.02 |
| *ΔhamG* + $^{15}$N-**1** | No | Yes | n/a | 5.88±0.0265 |
| *ΔhamCG* + $^{15}$N-**4** | Yes | No | 0.654±0.00100 | n/a |
| *ΔhamDG* + $^{15}$N-**4** | Yes | No | 0.654±0.00153 | n/a |

Note: Since each run in the MS has different amount of total data points and measurements were scattered over different time points among repeats, it was impossible to average data points without curve fitting or approximation, either of which will compromise the hight quality data obtained from the state-of-art HRMS. To assess how reproducible among the triplicates, calculating mean and standard error (SE) of the time point where maximum intensity is reached makes more sense.

**Supplementary Table 3.** Crystallographic data for the dihydrosydnone *N*-oxide (**1**).

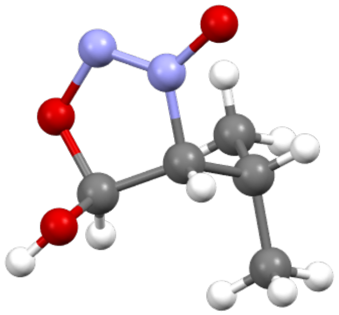

| | |
|---|---|
| Crystallised from | Ether and cyclopentane (antisolvent) |
| Empirical formula | $C_5H_{10}N_2O_3$ |
| Formula weight | 146.15 |
| Temperature/K | 160.0(1) |
| Crystal system | orthorhombic |
| Space group | Pbca |
| a/Å | 11.57870(10) |
| b/Å | 9.31170(10) |
| c/Å | 13.18000(10) |
| α/° | 90 |
| β/° | 90 |
| γ/° | 90 |
| Volume/Å$^3$ | 1421.03(2) |
| Z | 8 |
| $\rho_{calc}$g/cm$^3$ | 1.366 |
| μ/mm$^{-1}$ | 0.965 |
| F(000) | 624.0 |
| Crystal size/mm$^3$ | 0.26 × 0.19 × 0.12 |
| Radiation | Cu Kα (λ = 1.54184) |
| 2Θ range for data collection/° | 13.44 to 154.51 |
| Index ranges | -13 ≤ h ≤ 14, -11 ≤ k ≤ 10, -15 ≤ l ≤ 16 |
| Reflections collected | 9864 |
| Independent reflections | 1507 [$R_{int}$ = 0.0166, $R_{sigma}$ = 0.0106] |
| Data/restraints/parameters | 1507/0/98 |
| Goodness-of-fit on $F^2$ | 1.065 |
| Final R indexes [I>=2σ (I)] | $R_1$ = 0.0310, $wR_2$ = 0.0804 |
| Final R indexes [all data] | $R_1$ = 0.0316, $wR_2$ = 0.0809 |
| Largest diff. peak/hole / e Å$^{-3}$ | 0.27/-0.18 |

**Supplementary Table 4.** Crystallographic data for the amine analogue (**4**).

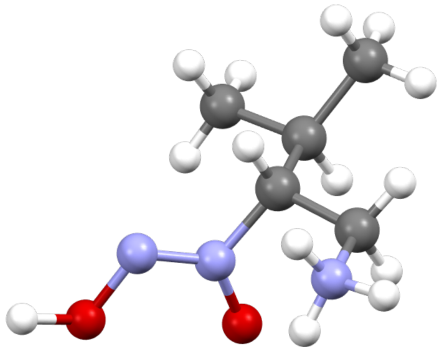

| | |
|---|---|
| Crystallised from | Methanol and ether (antisolvent) |
| Empirical formula | $C_{12}H_{35}ClN_6O_6$ |
| Formula weight | 394.91 |
| Temperature/K | 160.0(1) |
| Crystal system | monoclinic |
| Space group | $P2_1/c$ |
| a/Å | 9.9830(2) |
| b/Å | 21.2959(6) |
| c/Å | 11.0843(3) |
| α/° | 90 |
| β/° | 106.801(3) |
| γ/° | 90 |
| Volume/Å$^3$ | 2255.90(10) |
| Z | 4 |
| $\rho_{calc}$g/cm$^3$ | 1.163 |
| μ/mm$^{-1}$ | 1.809 |
| F(000) | 856.0 |
| Crystal size/mm$^3$ | 0.14 × 0.07 × 0.04 |
| Radiation | Cu Kα (λ = 1.54184) |
| 2Θ range for data collection/° | 8.30 to 154.72 |
| Index ranges | $-12 \le h \le 12$, $-26 \le k \le 25$, $-14 \le l \le 14$ |
| Reflections collected | 25904 |
| Independent reflections | 4767 [$R_{int}$ = 0.0434, $R_{sigma}$ = 0.0320] |
| Data/restraints/parameters | 4767/0/268 |
| Goodness-of-fit on $F^2$ | 1.058 |
| Final R indexes [I>=2σ (I)] | $R_1$ = 0.0471, $wR_2$ = 0.1349 |
| Final R indexes [all data] | $R_1$ = 0.0606, $wR_2$ = 0.1455 |
| Largest diff. peak/hole / e Å$^{-3}$ | 0.57/-0.31 |

**Supplementary Table 5.** Crystallographic data for the diol (**17**).

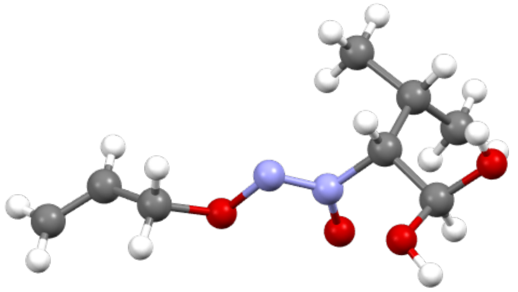

| | |
|---|---|
| Crystallised from | Pentane and diethyl ether |
| Empirical formula | $C_8H_{16}N_2O_4$ |
| Formula weight | 204.23 |
| Temperature/K | 160(1) |
| Crystal system | monoclinic |
| Space group | $P2_1/c$ |
| a/Å | 9.7461(2) |
| b/Å | 10.1007(4) |
| c/Å | 10.8278(2) |
| α/° | 90 |
| β/° | 90.358(2) |
| γ/° | 90 |
| Volume/Å3 | 1065.89(5) |
| Z | 4 |
| ϱcalcg/cm3 | 1.273 |
| μ/mm-1 | 0.860 |
| F(000) | 440.0 |
| Crystal size/mm3 | 0.25 × 0.21 × 0.02 |
| Radiation | CuKα (λ = 1.54184) |
| 2Θ range for data collection/° | 9.07 to 146.00 |
| Index ranges | -12 ≤ h ≤ 12, -12 ≤ k ≤ 11, -13 ≤ l ≤ 12 |
| Reflections collected | 18546 |
| Independent reflections | 2108 [$R_{int}$ = 0.0239, $R_{sigma}$ = 0.0101] |
| Data/restraints/parameters | 2108/0/138 |
| Goodness-of-fit on F2 | 1.043 |
| Final R indexes [I>=2σ (I)] | R1 = 0.0316, wR2 = 0.0821 |
| Final R indexes [all data] | R1 = 0.0335, wR2 = 0.0843 |
| Largest diff. peak/hole / e Å-3 | 0.20/-0.19 |

**Supplementary Table 6. Bacterial strains and plasmids used in this study.**

| Strain or plasmid | Characteristics | Source/Reference |
|---|---|---|
| ***Burkholderia cenocepacia*** | | |
| H111 | Cystic Fibrosis isolate, (Germany) | [11] |
| H111 *ΔhamC* | Unmarked *hamC* deletion mutant | [1] |
| H111 *ΔhamD* | Unmarked *hamD* deletion mutant | [1] |
| H111 *ΔhamG* | Unmarked *hamG* deletion mutant | This study |
| H111 *ΔhamCG* | Double deletion mutant of *hamC* and *hamG* | This study |
| H111 *ΔhamDG* | Double deletion mutant of *hamD* and *hamG* | This study |
| ***Escherichia coli*** | | |
| Top 10 | F-mcrAΔ(mrr-hsdRMS-mcrBC) φ80lacZΔM15 ΔlacX74 nupG recA1 araD139 Δ(ara-leu)7697 galE15 galK16 rpsL(StrR ) endA1 λ- | Invitrogen |
| CC118 | λpir Δ(ara,leu)7697araD139 ΔlacX74 galEgalKphoA20 thi-1rpsErpoB(RFR ) argE(am) recA1λpir+ | [12] |
| **Plasmids** | | |
| pSU11 | Promoter probe vector for lacZ fusion, GmR | [13] |
| pGPI-SceI::TetAR | Suicide plasmid with oriR6K, mob+, I-SceI restriction site; TpR TcR | [14] |
| pDAIGm-SceI | pDA17 plasmid carrying the I-SceI nuclease gene; GmR | [15] |
| phamA-lacZ | pSU11 containing the *hamA* promoter | [1] |

**Supplementary Fig 1.** Extracted-ion chromatogram (EIC) traces demonstrating no fragin was produced when feeding *ΔhamCG* and *ΔhamDG* with $^{15}$N-**1**, stacked with positive result from *ΔhamC* + $^{15}$N-**1**.

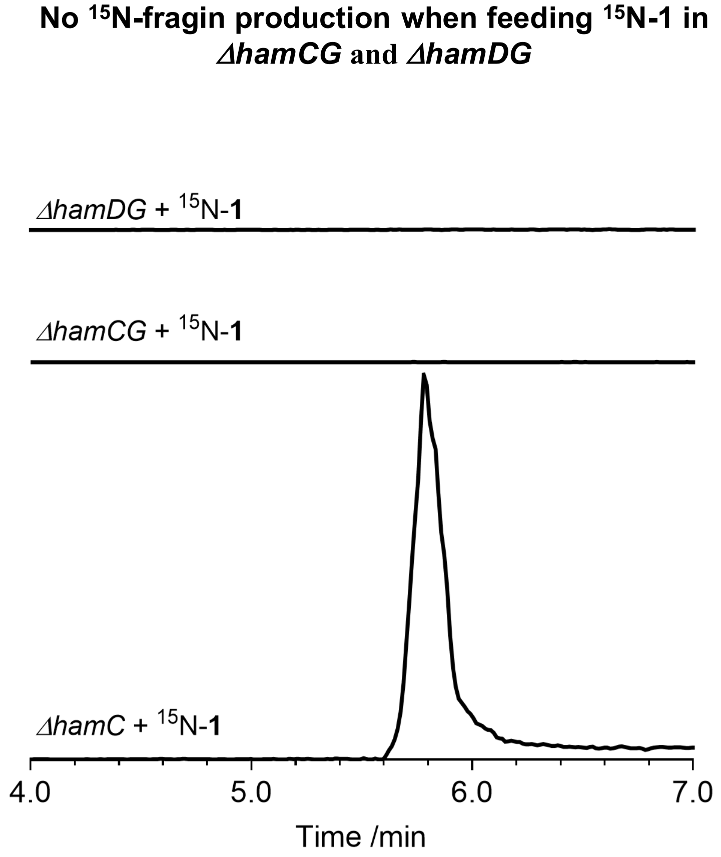

**Supplementary Fig 2.** EIC traces of fragins from supernatants spiked with (*R*) or (*S*)–fragins.

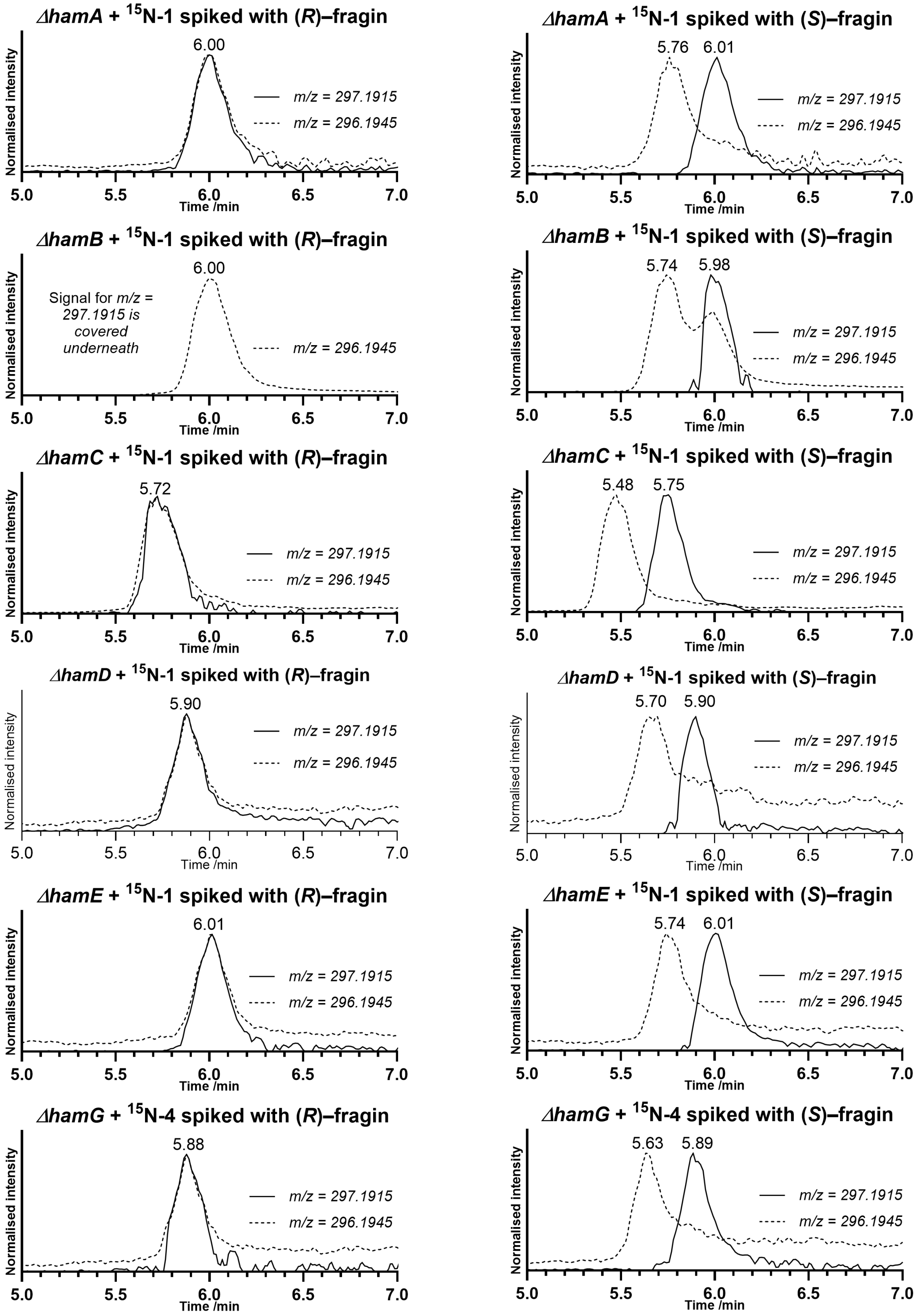

**Supplementary Fig 3.** EIC traces demonstrating no valdiazen was produced when feeding *ΔhamG* with $^{15}$N-**4**, stacked with positive result from *ΔhamC* + $^{15}$N-**1**.

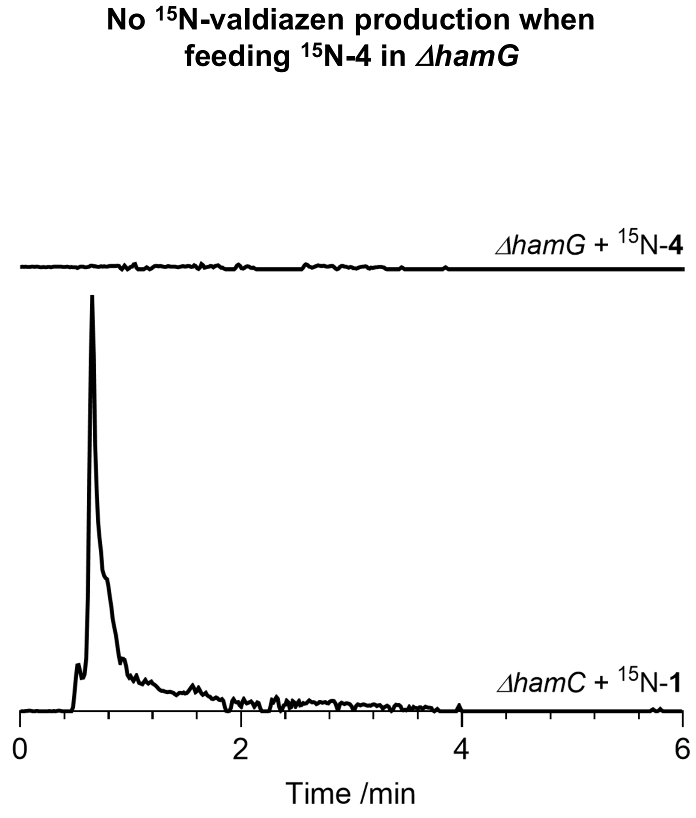

**Supplementary Fig 4.** Waterfall plots of EIC traces demonstrating the production of $^{15}$N-labelled fragin and valdiazen from *ΔhamA*, *ΔhamB and ΔhamE*.

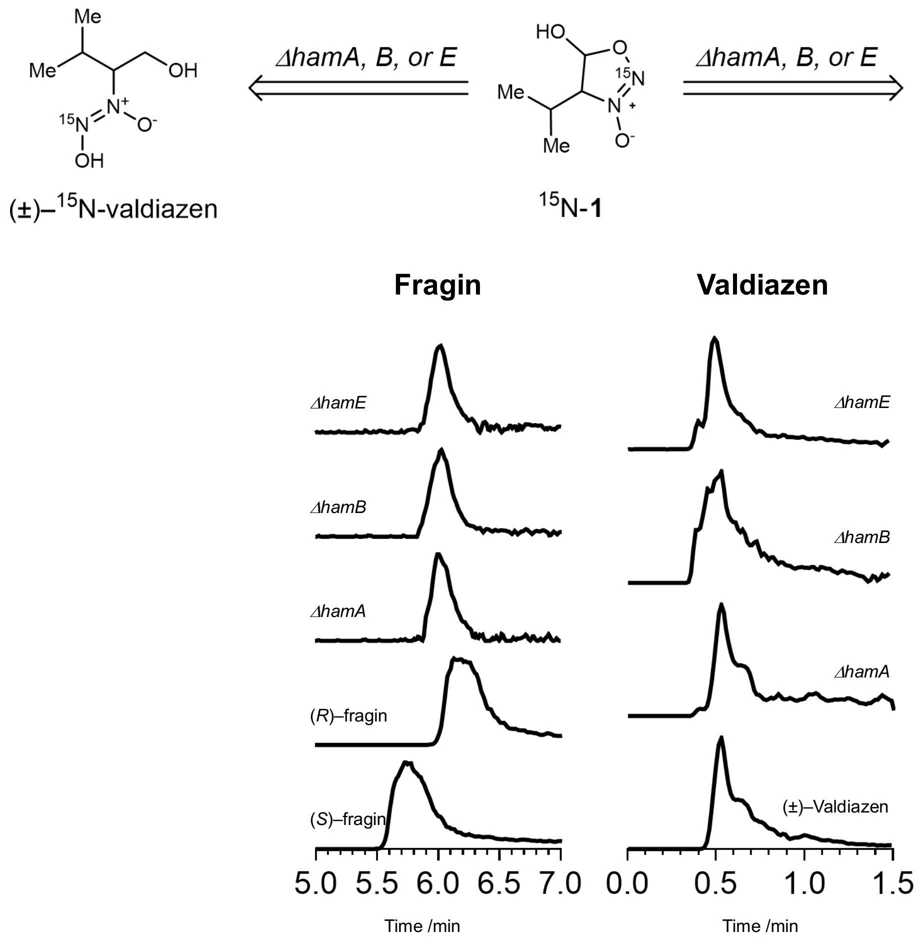

**Synthesis of 1,1-dimethoxy-3-methylbutan-2-one (7)**

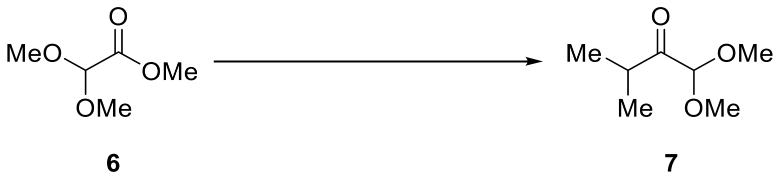

Under $N_2$, 2 M $AlMe_3$ in hexanes (20.4 mL, 40.8 mmol, 2.5 eq.) was slowly added to *N*,*O*–dimethylhydroxylamine hydrochloride (3.98 g, 40.8 mmol, 2.5 eq.) in dry DCM (22 mL) at 0 °C, the reaction mixture was allowed to warm up to r.t. and stirred for 1 hour at r.t.. The reaction mixture was cooled to 0 °C, a solution of methyl dimethoxyacetate (2.00 mL, 16.3 mmol, 1 eq.) in dry DCM (17.5 mL) was slowly added over 2 hours. The reaction mixture was allowed to warm up to r.t. and stirred for another 2 hours at this temperature. TLC in 50% EtOAc in hexanes showed full consumption of the starting material. The reaction mixture was then slowly poured into ice-cold 0.5 M aq. HCl (22 mL) and stirred at 0 °C for 30 min before the layers were separated. The aqueous phase was extracted with ether (2 × 20 mL). The combined organic phases were dried over anhydrous $Na_2SO_4$, filtered, and concentrated *in vacuo* (minimum 200 mbar, 40 °C) to give a Weinreb amide (2.65 g) as a light-yellow oil. TLC in neat EtOAc ($R_f$ = 0.6, faint under UV, oxidisable with $KMnO_4$) showed a very clean profile with minor impurity above the desired product spot. The compound is volatile under high stream of nitrogen gas.

Under $N_2$, to a solution of the above crude Weinreb amide (2.65 g) in dry THF (55 mL) at −78 °C, was added dropwise 3 M isopropylmagnesium bromide in 2-MeTHF (7.60 mL, 22.7 mmol, 1.3 eq.). The solution was then warmed to r.t. and stirred for 2 hours. TLC in neat EtOAc and 20% EtOAc in hexanes showed the completion of the reaction after 2 hours. The reaction was then cooled to 0 °C and quenched with sat. aq. $NH_4Cl$ (20 mL). The reaction was warmed back to r.t. and extracted with ether (3 × 50 mL). The combined organic layers were washed with brine (50 mL), dried over anhydrous $Na_2SO_4$, filtered, and concentrated *in vacuo* to give a colourless oil. The crude (2.11 g) was loaded onto silica gel (250 mL) and eluted with 20% ether in hexanes (2 L) to give 1,1-dimethoxy-3-methylbutan-2-one (1.34 g, 56% over 2 steps) as a colourless oil. The compound was found to be volatile and evaporated below 50 mbar at 40 °C on rotary evaporator. The analytics agree with the literature.[16]

**$R_f$** = 0.50 ($SiO_2$, 20% ether in hexanes, $KMnO_4$ stain).

**$^1$H NMR** (400 MHz, $CDCl_3$) $\delta$ = 4.61 (d, $J$ = 0.4 Hz, 1H), 3.39 (d, $J$ = 0.4 Hz, 6H), 3.01 (pd, $J$ = 6.9, 0.4 Hz, 1H), 1.10 (d, $J$ = 0.4 Hz, 3H), 1.09 (d, $J$ = 0.5 Hz, 3H).

**$^{13}$C NMR** (101 MHz, $CDCl_3$) $\delta$ = 209.25, 103.12, 54.60, 35.99, 18.37.

**ESI-HRMS** (MeOH + NaI): $m/z$ 169.08352 ($C_7H_{14}NaO_3^+$; [$M$+Na]$^+$; calc. 169.08352).

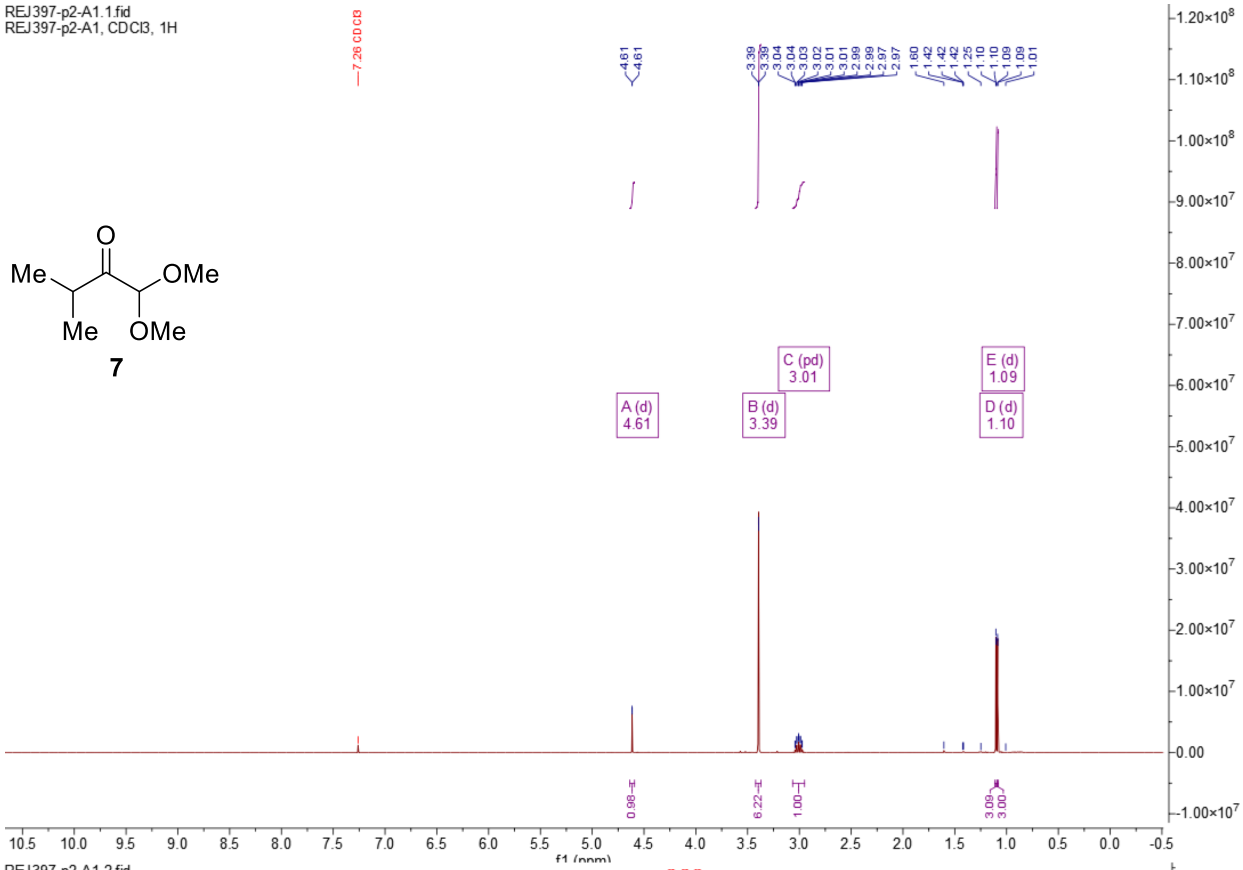
REJ397-p2-A1.1.fid
REJ397-p2-A1, CDCl3, 1H
7

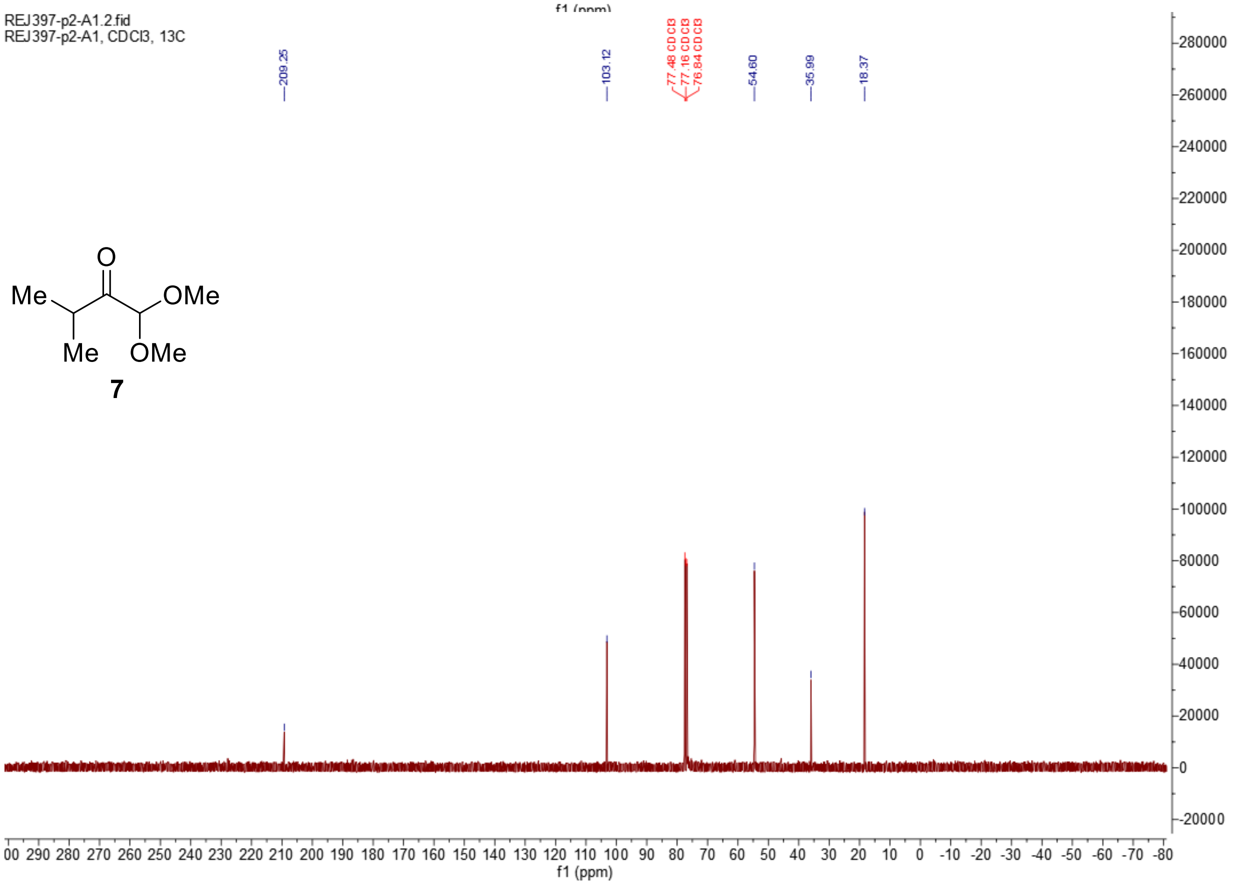
REJ397-p2-A1.2.fid
REJ397-p2-A1, CDCl3, 13C
7

**Synthesis of (*E*) and (*Z*)–1,1-dimethoxy-3-methylbutan-2-one oximes (8)**

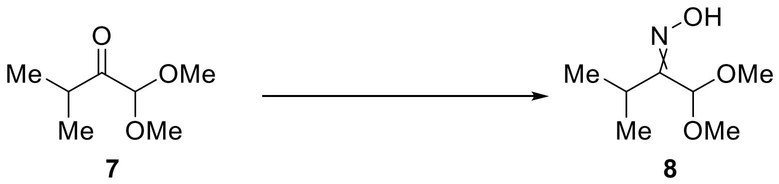

To a mixture of hydroxylamine hydrochloride (359 mg, 5.17 mmol, 1.2 eq.) and NaOAc (707 mg, 8.62 mmol, 2 eq.) in EtOH (30 mL) and water (13 mL), was added ketone **7** (630 mg, 4.31 mmol, 1 eq.) in a single portion. The mixture was then heated to 70 °C for 3 hours. TLC in 20% EtOAc in hexanes showed full consumption of the starting material and two oxidisable spots by $KMnO_4$ stain. The mixture was cooled to r.t., concentrated *in vacuo* and diluted with water (10 mL). The aqueous layer was extracted with EtOAc (3 × 20 mL). The combined organic phases were dried over anhydrous $Na_2SO_4$, filtered, and concentrated *in vacuo* to give a colourless oil. The crude (760 mg) was loaded onto silica gel (50 mL) and the compounds were eluted with 20% EtOAc in hexanes to give the corresponding oximes as a colourless oil consisting of a mixture of *E*/*Z* isomers (545 mg, 78%).

$\mathbf{R_f}$ = 0.57 and 0.36 ($SiO_2$, 20% EtOAc in hexanes, $KMnO_4$ stain).

**$^1$H NMR** (400 MHz, $CDCl_3$) $\delta$ = 5.50 (s, 1H), 4.67 (s, 3H), 3.45 (s, 6H), 3.37 (s, 17H), 3.07 (p, *J* = 7.1 Hz, 3H), 2.79 (p, *J* = 6.9 Hz, 1H), 1.22 (d, *J* = 7.1 Hz, 16H), 1.14 (d, *J* = 6.9 Hz, 6H).

**$^{13}$C NMR** (101 MHz, $CDCl_3$) $\delta$ = 162.22, 160.19, 104.00, 97.95, 55.72, 54.42, 28.36, 26.55, 21.20, 18.69.

**FTIR $\tilde{\nu}$ (cm$^{-1}$)** = 3282m, 2967m, 2935m, 2877m, 2833m, 1701w, 1547w, 1453m, 1383m, 1360m, 1214m, 1190m, 1155w, 1107s, 1068s, 1014m, 931s, 901s, 828m, 787m, 764m, 596w, 568w.

**ESI-HRMS** (MeOH): *m*/*z* 184.09470 ($C_7H_{15}O_3NNa^+$; [*M*+Na]$^+$; calc. 184.09441).

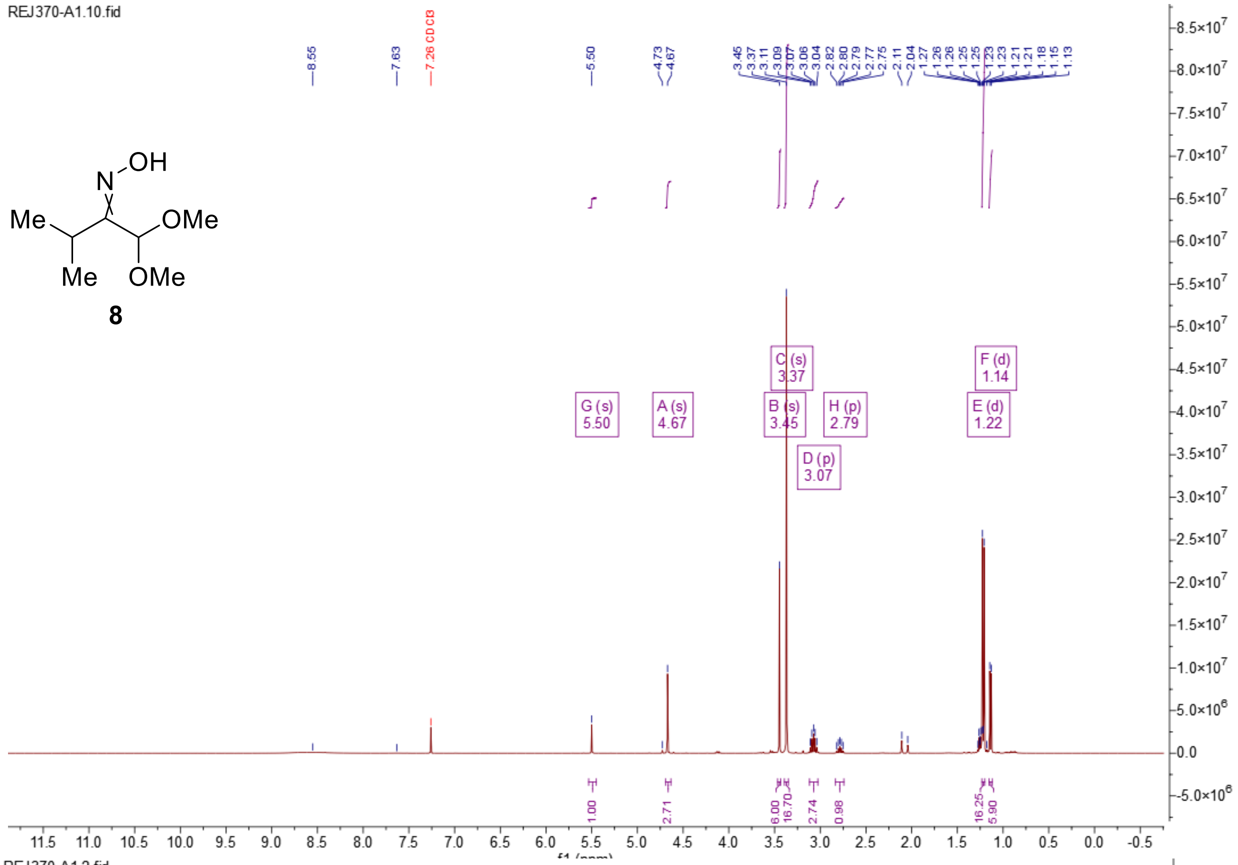
REJ370-A1.10.fid
8
8.55
7.63
7.26 CDCl3
5.50
4.73
4.67
3.45
3.37
3.11
3.09
3.07
3.06
3.04
2.82
2.80
2.79
2.77
2.75
2.11
2.04
1.27
1.26
1.26
1.25
1.23
1.23
1.21
1.18
1.15
1.13
G (s) 5.50
A (s) 4.67
C (s) 3.37
B (s) 3.45
D (p) 3.07
H (p) 2.79
F (d) 1.14
E (d) 1.22
1.00
2.71
6.00
16.70
2.74
0.98
16.25
5.90
f1 (ppm)

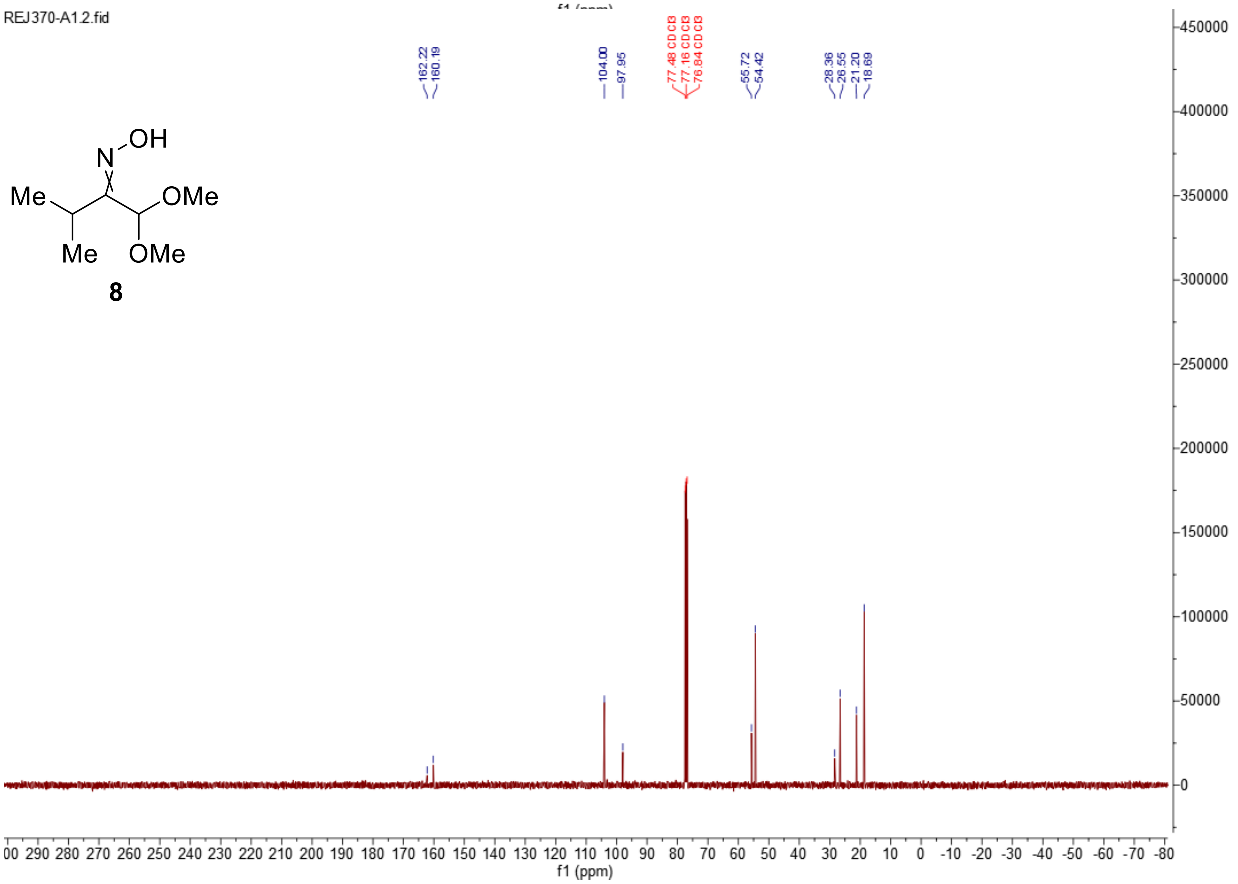
REJ370-A1.2.fid
8
162.22
160.19
104.00
97.95
77.48 CDCl3
77.16 CDCl3
76.84 CDCl3
55.72
54.42
28.36
26.55
21.20
18.69
f1 (ppm)

**Synthesis of (*Z*)–1-(1,1-dimethoxy-3-methylbutan-2-yl)-2-hydroxydiazene 1-oxide-2-$^{15}N$ (9)**

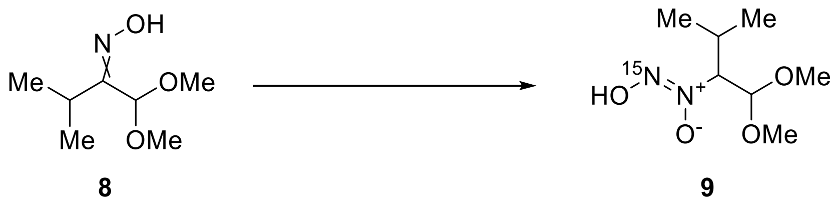

To a stirred solution of oximes **8** (1.06 g, 6.58 mmol, 1 eq.) in dry ethanol (21 mL) under $N_2$ at 0 °C, $NaCNBH_3$ (522 mg, 0.744 mmol, 1.2 eq.) was added followed by dropwise addition of 1.25 M HCl in ethanol (6.40 mL, 8.00 mmol, 1.2 eq.). The reaction mixture was then stirred at r.t. for 1 hour. UHPLC-MS showed full conversion. The crude mixture was concentrated *in vacuo*, treated with water (20 mL) and EtOAc (20 mL) before neutralising with sat. aq. $Na_2CO_3$ (2 mL). The aqueous layer was extracted with EtOAc (5 × 20 mL). The combined organic layers were dried over anhydrous $Na_2SO_4$, filtered, and concentrated *in vacuo* to give a crude hydroxylamine (1.09 g, quant.) as a light pink oil. UHPLC-MS showed the desired mass.

To a solution of the above hydroxylamine (200 mg, 1.23 mmol, 1 eq.) in 1:1 EtOH/$H_2O$ (2 mL) at 0 °C, was added 1 M aq. HCl (1.40 mL, 1.40 mmol, 1.1 eq.) dropwise. The mixture was degassed with argon for 10 mins. In a separate flask, a solution of $Na^{15}NO_2$ (95.0 mg, 1.36 mmol, 1.1 eq.) in water (1 mL) was also degassed with argon for 10 mins before being added dropwise to the hydroxylamine solution at 0 °C. The reaction was then stirred at 0 °C for 30 minutes. UHPLC-MS showed full conversion of the starting material and the formation of the desired product. The mixture was treated with sat. aq. $NaHCO_3$ (4 mL) and freshly distilled ether (5 mL). The aqueous phase was washed with freshly distilled ether (3 × 5 mL) before being acidified with 1 M aq. HCl (6 mL). The aqueous phase was then extracted with ether (5 × 10 mL). The combined organic phases were dried over anhydrous $Na_2SO_4$, filtered, and concentrated *in vacuo* to give NONOate **9** (130 mg, 55% over 2 steps) as a light-yellow oil with a characteristic smell.

**$^1$H NMR** (400 MHz, Methanol-*d₄*) $\delta$ = 4.85 (d, $J$ = 7.9 Hz, 1H), 4.16 (ddd, $J$ = 7.9, 6.0, 1.9 Hz, 1H), 3.44 (s, 3H), 3.38 (s, 3H), 2.27 (pd, $J$ = 6.9, 5.9 Hz, 1H), 1.03 (d, $J$ = 6.9 Hz, 3H), 0.99 (d, $J$ = 7.0 Hz, 3H).

**$^{13}$C NMR** (101 MHz, Methanol-$d_4$) $\delta$ = 103.62, 78.92, 55.73, 54.99, 29.47, 19.73, 18.24.

**FTIR $\tilde{\nu}$ (cm$^{-1}$)** = 2968m, 2939m, 2838w, 1457m, 1392m, 1371m, 1281m, 1192m, 1150m, 1119m, 1055s, 992m, 971m, 945m, 909m, 855w, 829m, 781m, 702m, 668m, 593m, 531w, 479w.

**ESI-HRMS** (MeCN): *m/z* 216.09728 ($C_7H_{16}O_4N^{15}NNa^+$; $[M+Na]^+$; calc. 216.09726).

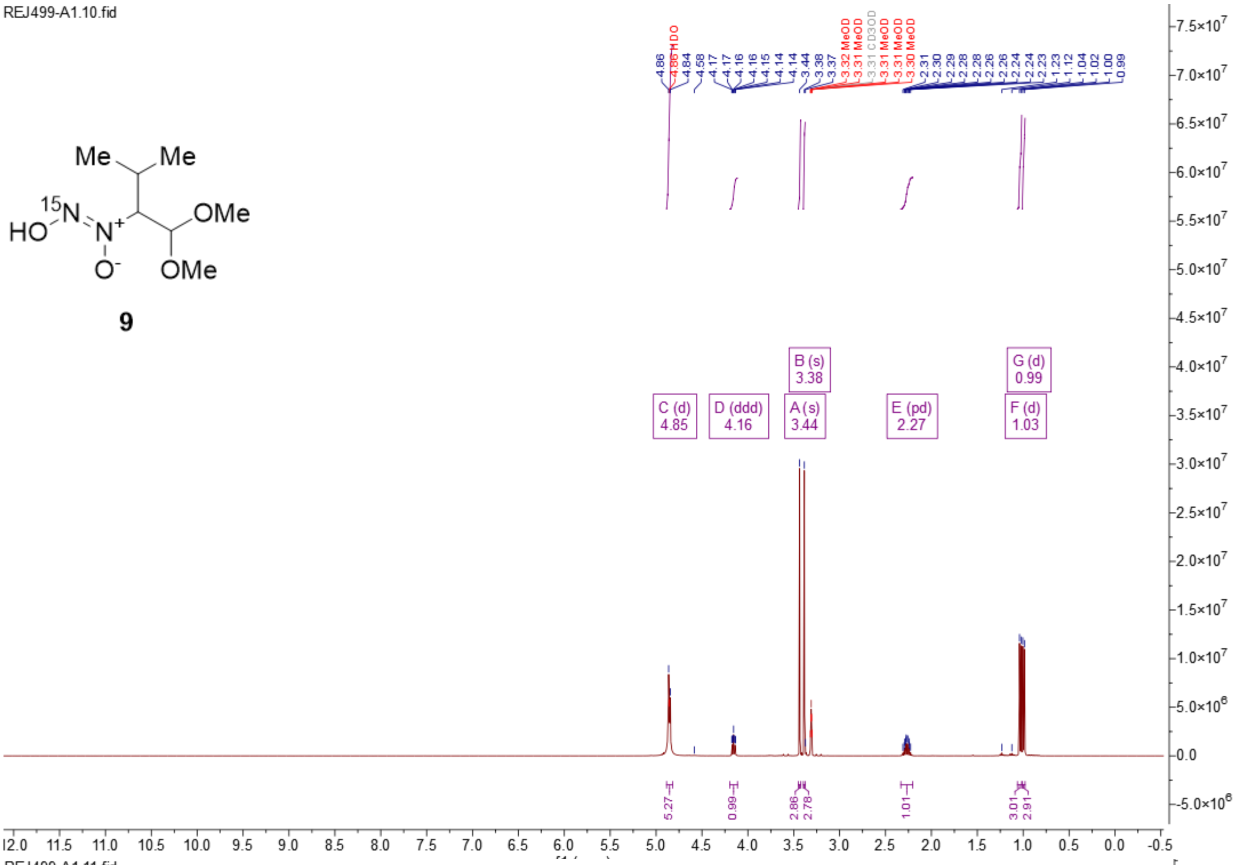
REJ499-A1.10.fid
Me
Me
15N
HO
N+
O-
OMe
OMe
9
C (d) 4.85
D (ddd) 4.16
A (s) 3.44
B (s) 3.38
E (pd) 2.27
F (d) 1.03
G (d) 0.99
5.27
0.99
2.86
2.78
1.01
3.01
2.91

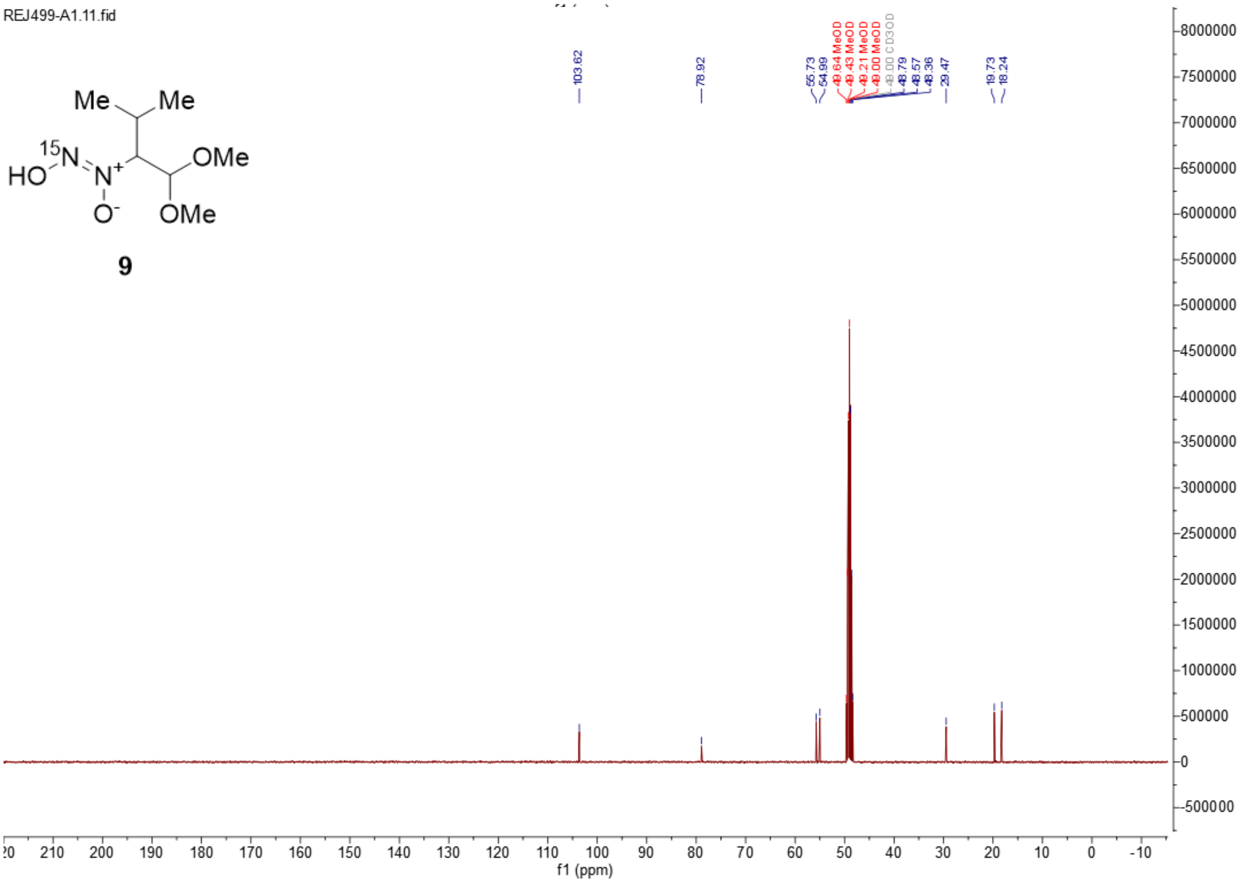
REJ499-A1.11.fid
Me
Me
15N
HO
N+
O-
OMe
OMe
9
103.62
78.92
55.73
54.99
29.47
19.73
18.24
f1 (ppm)

**Synthesis of 5-hydroxy-4-isopropyl-4,5-dihydro-1,2,3-oxadiazole 3-oxide-2-$^{15}$*N* ($^{15}$N-1)**

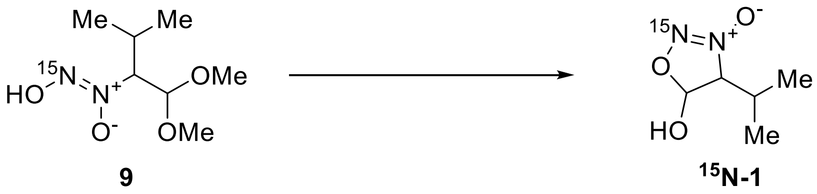

The protected NONOate **9** (49.0 mg, 0.254 mmol, 1 eq.) was treated with 10% aq. HCl (1.20 mL, 3.81 mmol, 15 eq.) and stirred at 50 °C for 3 hours. The mixture was reduced to less than 0.5 mL before being purified by preparative HPLC (*Phenomenex Synergi™* 10 µm Hydro-RP 80 Å, 250 mm × 21.2 mm, 20 mL/min, 0%–50% MeCN + 0.1% formic acid in MiliQ water + 0.1% formic acid, 30 minutes, $R_t$ = 15.0 min.) to give dihydrosydnone *N*-oxide $^{15}$N-**1** (17.0 mg, 46%) as a sticky fluffy white solid. The solid was recrystallised from ether/cyclopentane for single crystal X-ray crystallography.

**m.p.=** 55.8–60.2 °C

**$^1$H NMR** (500 MHz, Methanol-*d$_4$*) $\delta$ = 5.95 (d, *J* = 2.8 Hz, 1H), 4.15 (dd, *J* = 4.4, 2.8 Hz, 1H), 2.43 (pd, *J* = 7.0, 4.4 Hz, 1H), 1.09 (d, *J* = 7.0 Hz, 4H), 0.98 (d, *J* = 6.9 Hz, 3H).

**$^{13}$C NMR** (126 MHz, Methanol-*d$_4$*) $\delta$ = 101.29, 85.54, 29.18, 18.01, 17.02.

**FTIR $\tilde{\nu}$ (cm$^{-1}$)** = 3153m, 2973w, 1441m, 1397m, 1376w, 1343m, 1305m, 1246m, 1147m, 1130m, 1093m, 1008w, 976m, 959s, 933m, 903m, 843m, 807s, 765m, 694m, 612w, 576m, 501m, 477w.

**ESI-HRMS** (MeCN): *m/z* 146.05906 ($C_5H_9O_3N^{15}N^-$; [*M*–H]$^-$; calc. 146.05890).

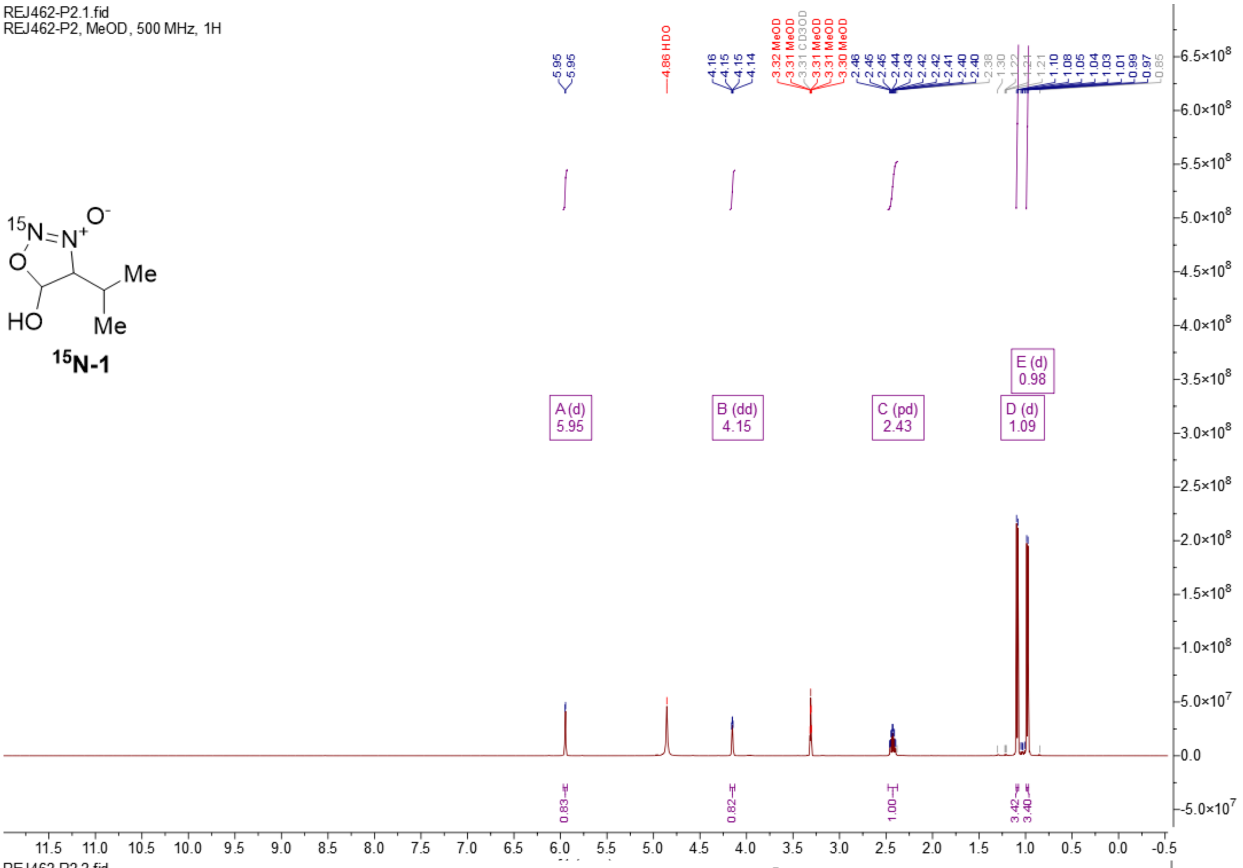
REJ462-P2.1.fid
REJ462-P2, MeOD, 500 MHz, 1H
15N-1
A (d) 5.95
B (dd) 4.15
C (pd) 2.43
D (d) 1.09
E (d) 0.98
f1 (ppm)

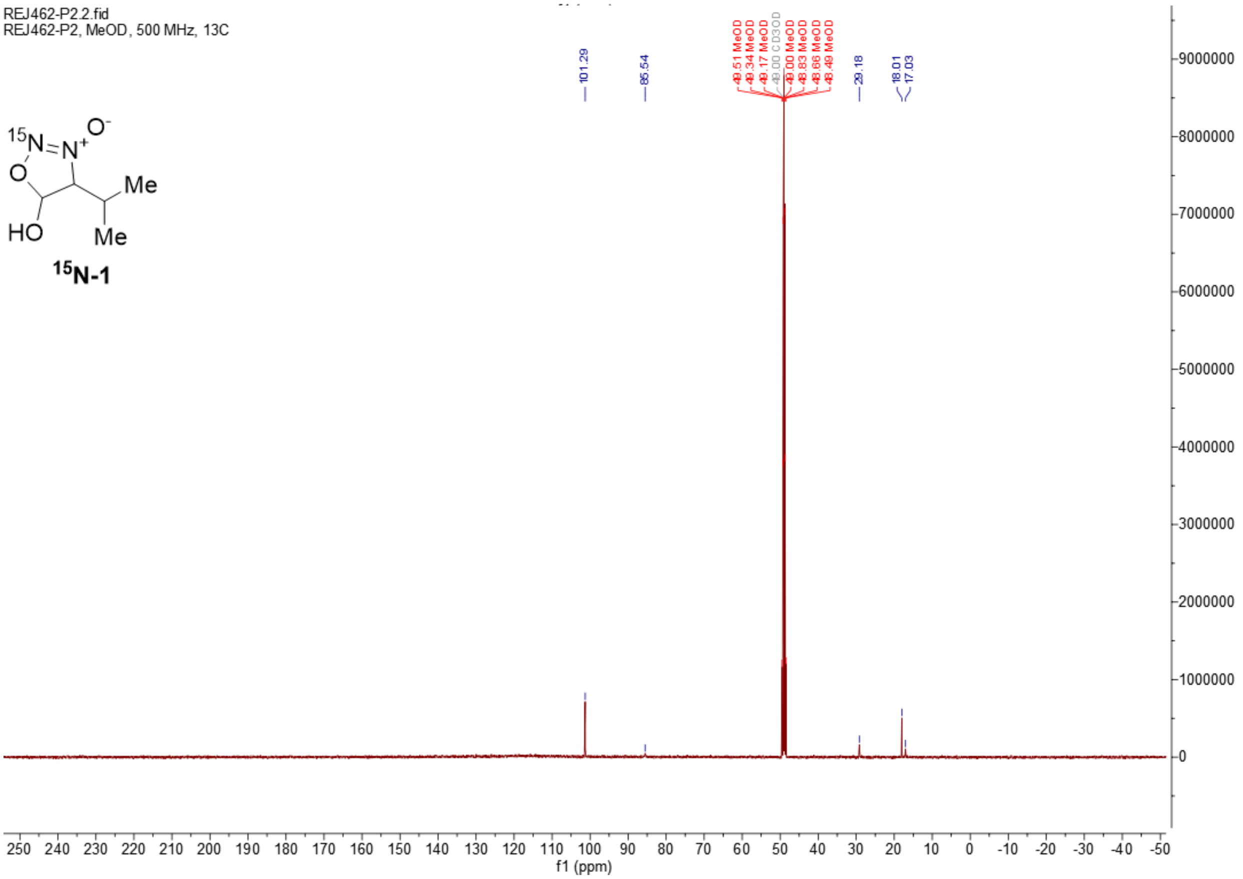
REJ462-P2.2.fid
REJ462-P2, MeOD, 500 MHz, 13C
15N-1
101.29
85.54
29.18
18.01
17.03
f1 (ppm)

**Synthesis of *tert*-butyl (3-methyl-2-oxobutyl) carbamate (11)**

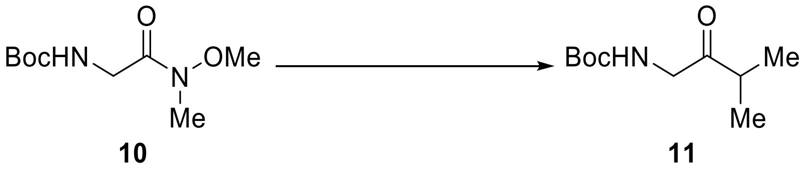

Under $N_2$, to a solution of Weinreb amide **10** (2.00 g, 9.16 mmol, 1 eq.) in dry THF (29 mL) at −78 °C, was added dropwise 3 M isopropylmagnesium bromide in 2-MeTHF (4.00 mL, 12.0 mmol, 1.3 eq.). The reaction was then warmed to r.t. and stirred for 2 hours. TLC in neat EtOAc and 20% EtOAc in hexanes were used for reaction control. Upon completion, the reaction then was cooled to 0 °C and quenched with sat. aq. $NH_4Cl$ (10 mL). The reaction was warmed back to r.t. and extracted with ether (3 × 50 mL). The combined organic layers were washed with brine (20 mL), dried over $Na_2SO_4$, filtered, and concentrated *in vacuo* to give the ketone **11** (889 mg, 49%) as a colourless oil. The analytics agree with literature.[17]

**$R_f$** = 0.50 ($SiO_2$, 20% EtOAc in hexanes, $KMnO_4$ stain).

**$^1$H NMR** (400 MHz, $CDCl_3$) $\delta$ = 5.24 (s, 1H), 4.09 (d, $J$ = 4.6 Hz, 2H), 2.64 (hept, $J$ = 6.9 Hz, 1H), 1.45 (s, 9H), 1.15 (s, 3H), 1.13 (s, 3H).

**$^{13}$C NMR** (101 MHz, $CDCl_3$) δ 209.49, 79.90, 48.56, 38.87, 28.48, 18.36.

**ESI-HRMS** (MeCN): $m/z$ 224.12581 ($C_{10}H_{19}O_3NNa^+$; $[M+Na]^+$; calc. 224.12571).

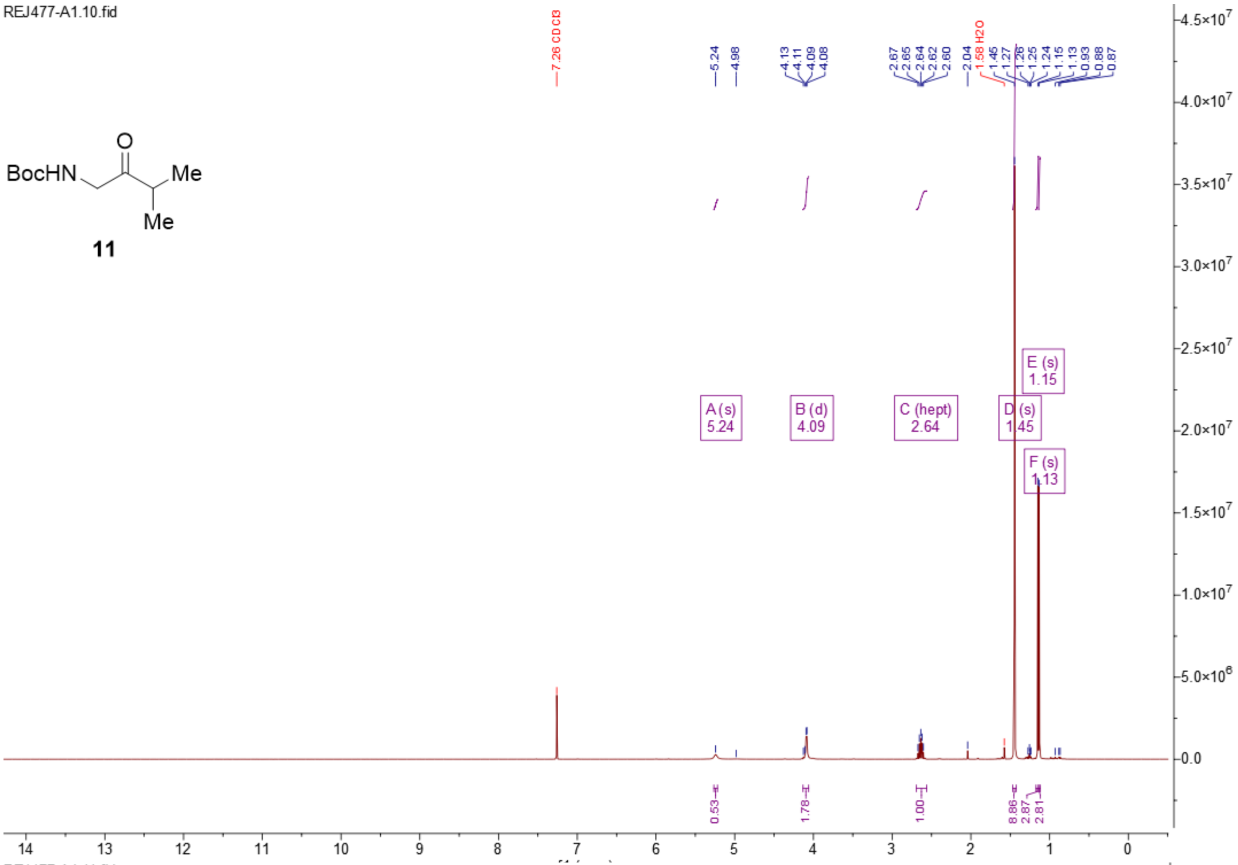
REJ477-A1.10.fid
BocHN
O
Me
Me
11
7.26 CDCl3
5.24
4.98
4.13
4.11
4.09
4.08
2.67
2.65
2.64
2.62
2.60
2.04
1.58 H2O
1.45
1.27
1.26
1.25
1.24
1.15
1.13
0.93
0.88
0.87
A (s)
5.24
B (d)
4.09
C (hept)
2.64
D (s)
1.45
E (s)
1.15
F (s)
1.13
0.53
1.78
1.00
8.86
2.87
2.81
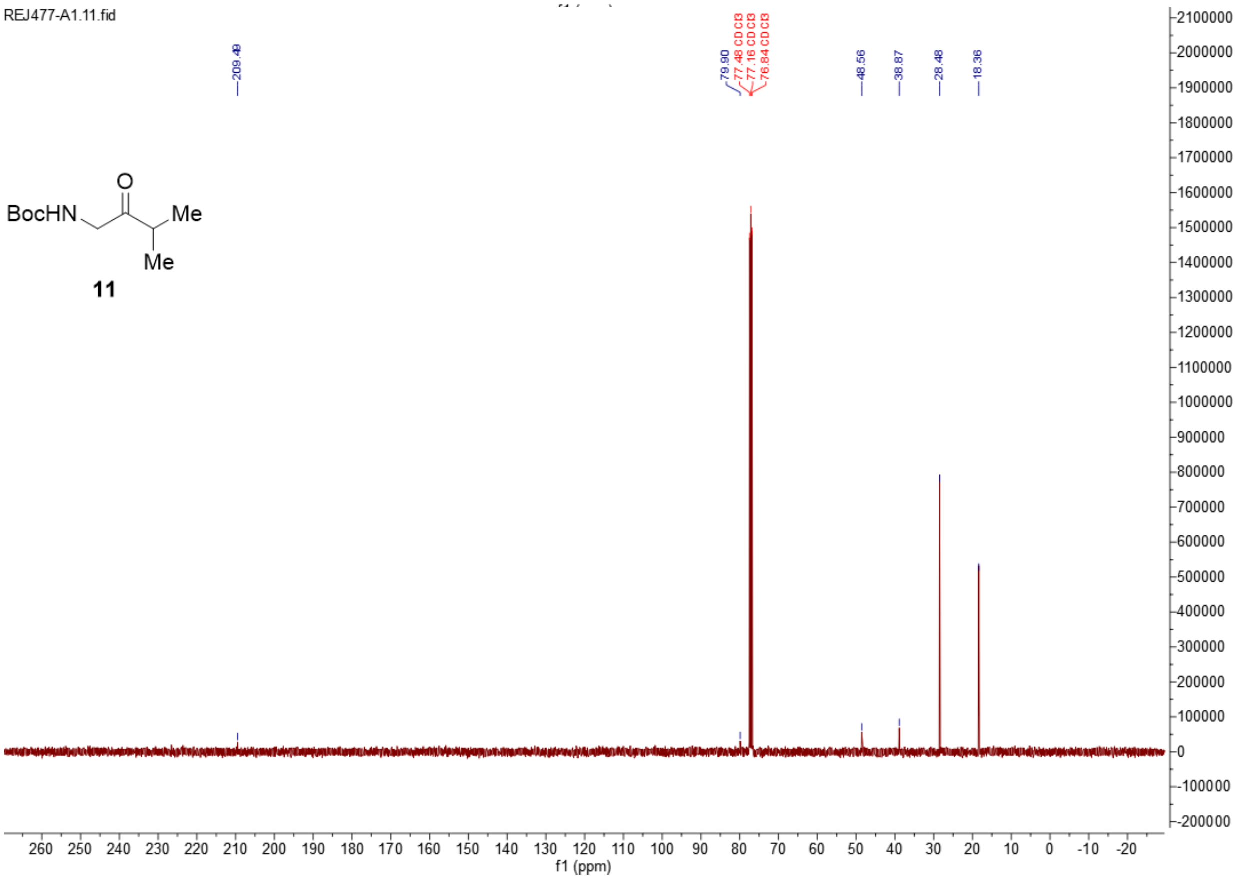
REJ477-A1.11.fid
BocHN
O
Me
Me
11
209.49
79.90
77.48 CDCl3
77.16 CDCl3
76.84 CDCl3
48.56
38.87
28.48
18.36
f1 (ppm)

**Synthesis of *tert*-butyl (2-(hydroxyimino)-3-methylbutyl) carbamate (12)**

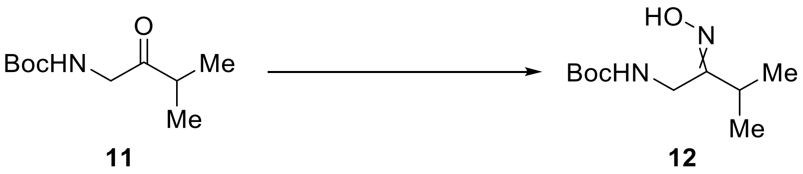

To a mixture of hydroxylamine hydrochloride (380 mg, 5.30 mmol, 1.2 eq.) and KOAc (868 mg, 8.84 mmol, 2 eq.) in EtOH (30 mL) and water (13 mL), was added ketone **11** (889 mg, 4.42 mmol, 1 eq.) in a single portion. The mixture was then heated to 70 °C for 3 hours. TLC in 20% EtOAc in hexanes showed full consumption of the starting material and two oxidisable spots by $KMnO_4$ stain. The mixture was cooled to r.t., concentrated *in vacuo*, and diluted with water (10 mL). The aqueous layer was extracted with EtOAc (3 × 20 mL). The combined organic phases were dried over anhydrous $Na_2SO_4$, filtered, and concentrated *in vacuo* to give a colourless oil. The crude (1.22 g) was loaded onto silica gel (120 mL) and eluted with 20% EtOAc in hexanes to give the corresponding oximes as a colourless oil consisting of a mixture of *E*/*Z* isomers (850 mg, 89%) as a white solid.

**$R_f$** = 0.38 and 0.30 ($SiO_2$, 20% EtOAc in hexanes, $KMnO_4$ stain).

**m.p.**= 94.5–98.2 °C

**$^1$H NMR** (400 MHz, $CDCl_3$) $\delta$ = 8.26 (s, 1H), 7.59 (s, 1H), 5.24 (s, 1H), 5.15 (s, 1H), 3.97 (d, $J$ = 6.3 Hz, 3H), 3.90 (d, $J$ = 5.0 Hz, 2H), 3.39 (p, $J$ = 7.1 Hz, 1H), 2.59 (hept, $J$ = 6.9 Hz, 2H), 1.45 (s, 23H), 1.12 (d, $J$ = 6.8 Hz, 10H), 1.09 (d, $J$ = 7.0 Hz, 6H).

**$^{13}$C NMR** (101 MHz, $CDCl_3$) $\delta$ = 163.26, 155.77, 79.75, 40.06, 36.39, 32.90, 28.52, 25.86, 19.62, 18.74.

**FTIR $\tilde{\nu}$ (cm$^{-1}$)** = 3323m, 2982w, 2965w, 1704w, 1681s, 1539s, 1454m, 1434m, 1390m, 1365m, 1341w, 1279s, 1249s, 1166s, 1147s, 1110m, 1038m, 1020m, 951m, 937s, 912m, 899w, 887m, 858m, 771w, 752m, 722m, 677s, 597m.

**ESI-HRMS** (MeCN): *m/z* 217.15459 ($C_{10}H_{21}O_3N_2^+$; [*M*+H]$^+$; calc. 217.15467).

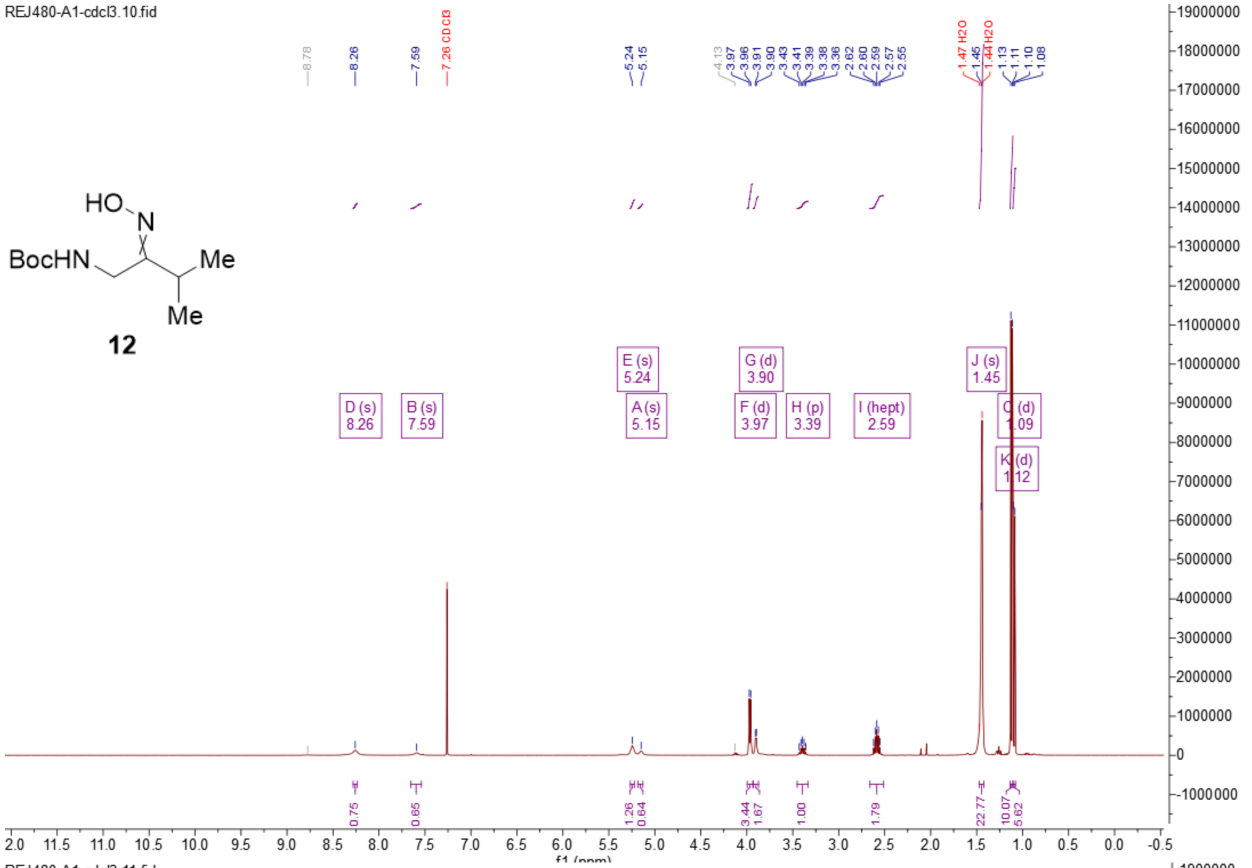
REJ480-A1-cdcl3.10.fid
HO
N
BocHN
Me
Me
12

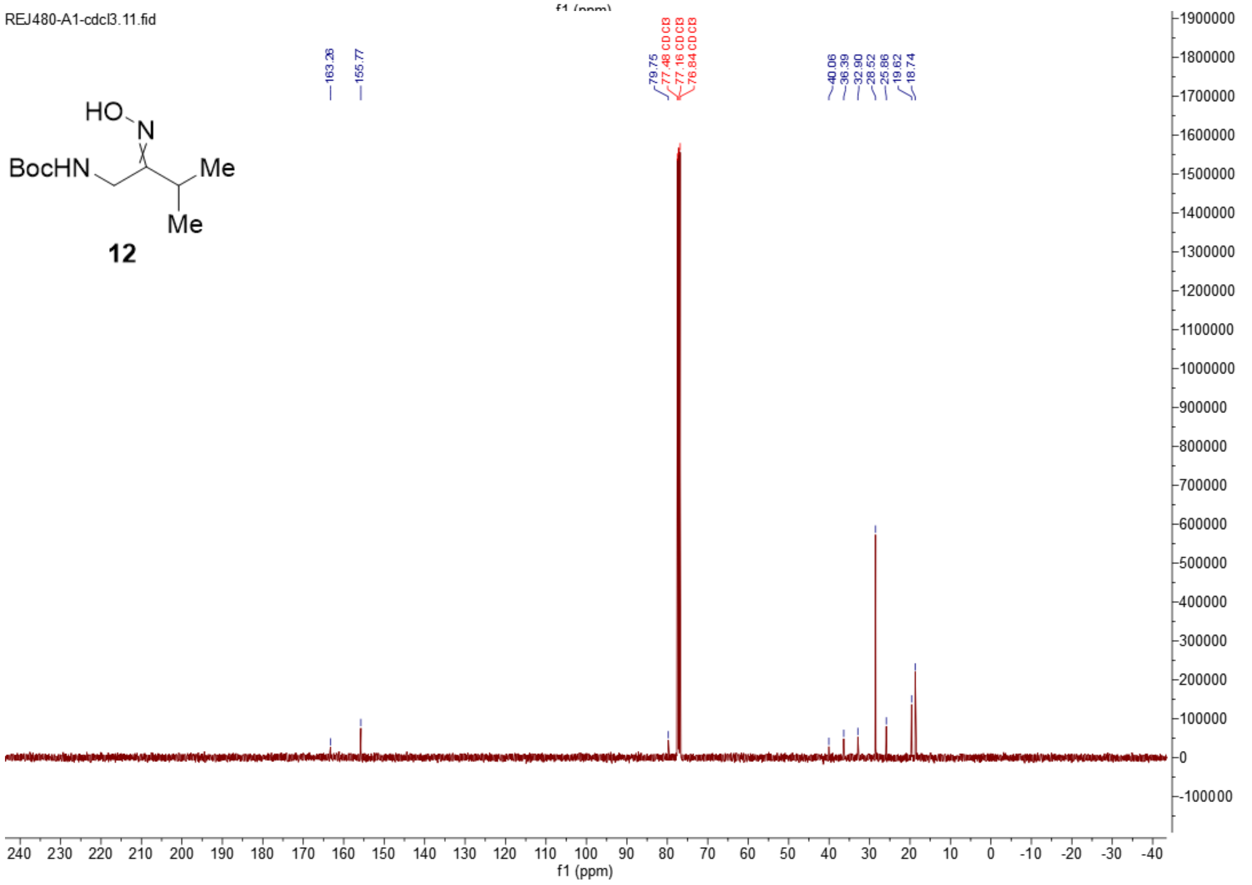
REJ480-A1-cdcl3.11.fid
HO
N
BocHN
Me
Me
12
f1 (ppm)

**Synthesis of (*Z*)–1-(1-amino-3-methylbutan-2-yl)-2-hydroxydiazene 1-oxide-2-$^{15}N$ ($^{15}$N-4)**

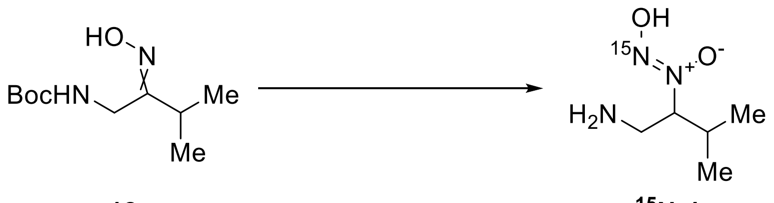


To a stirred solution of oxime **12** (207 mg, 0.957 mmol) in dry ethanol (3.00 mL) under $N_2$ at 0 °C, $NaCNBH_3$ (180 mg, 2.87 mmol, 3 eq.) was added followed by dropwise addition of 1.25 M ethanolic HCl (2.30 mL, 2.88 mmol) at 0 °C. The reaction was then warmed to r.t. and stirred for 1 hour. All starting material was consumed based on TLC at 20% EtOAc in hexanes. UHPLC-MS showed full consumption of the starting material and the formation of the desired product mass. The mixture was treated with EtOAc (10 mL) and sat. aq. $Na_2CO_3$ (3 mL). The aqueous layer was extracted with EtOAc (4 × 10 mL). The combined organic layers were washed with brine (10 mL), dried over anhydrous $Na_2SO_4$, filtered, and concentrated *in vacuo* to give the respective hydroxylamine (201 mg) as a colourless gum.

To a solution of the above hydroxylamine (201 mg, 0.921 mmol, 1 eq.) in DCM (1.70 mL) at 0 °C, was added TFA (711 µL, 9.57 mmol, 10 eq.). The reaction was stirred for 1 hour at r.t.. UHPLC-MS showed full consumption of the starting material and the formation of the desired product as a TFA salt. The mixture was concentrated *in vacuo*, reconstituted in water (1 mL) and filtered through DSC-18 cartridge (5 g bed) with the aid of water to give the corresponding amine (358 mg) as a TFA salt.

To a solution of the above amine (358 mg, 1.03 mmol, 1 eq.) in 1:1 EtOH/$H_2O$ (3.00 mL) at 0 °C, was added 1 M aq. HCl (2.20 mL, 2.20 mmol, 2.2 eq.) dropwise. The mixture was degassed with argon for 10 minutes. In a separate flask, a solution of $Na^{15}NO_2$ (81.0 mg, 1.16 mmol, 1.1 eq.) in water (1.00 mL) was degassed with argon for 10 minutes before being added dropwise to the hydroxylamine solution at 0 °C. The reaction was then stirred at 0 °C for 30 minutes. UHPLC-MS showed full conversion of the starting material and the formation of the desired product eluting at the solvent front. The mixture was concentrated *in vacuo* and treated with acetone (1 mL). The suspension was filtered, and the filtrate was concentrated *in vacuo*. The crude was purified by preparative HPLC (*Phenomenex Synergi*$^{TM}$ 10 μm Hydro-RP 80 Å, 250 mm × 21.2 mm, 20 mL/min, 100% MiliQ water + 0.1% formic acid, 10 minutes, $R_t$ = 6.5 minutes) to give β-amino NONOate $^{15}$N-**4** (30.0 mg, 21%) as a white solid. The solid was recrystallised from methanol/ether for single crystal X-ray crystallography.

**m.p.** = 50.2–56.2 °C

**$^1$H NMR** (400 MHz, Methanol-$d_4$) $\delta$ = 4.29 (t, $J$ = 9.4 Hz, 1H), 3.57 (dd, $J$ = 13.8, 9.7 Hz, 1H), 3.36 (dd, $J$ = 13.7, 2.6 Hz, 1H), 2.20 (h, $J$ = 6.8 Hz, 1H), 1.06 (d, $J$ = 6.7 Hz, 3H), 0.94 (d, $J$ = 6.5 Hz, 3H).

**$^{13}$C NMR** (101 MHz, Methanol-$d_4$) $\delta$ = 39.99, 30.53, 19.16, 19.09.

**FTIR $\tilde{\nu}$ (cm$^{-1}$)** = 2973br m, 1668s, 1537m, 1432m, 1377m, 1304w, 1273w, 1179s, 1131s, 1068m, 976m, 925m, 900m, 837m, 799m, 722s, 598w, 518m.

**ESI-HRMS** ($H_2O$): *m/z* 149.10495 ($C_5H_{14}O_2N_2{}^{15}N^+$; [$M$+H]$^+$; calc. 149.10509).

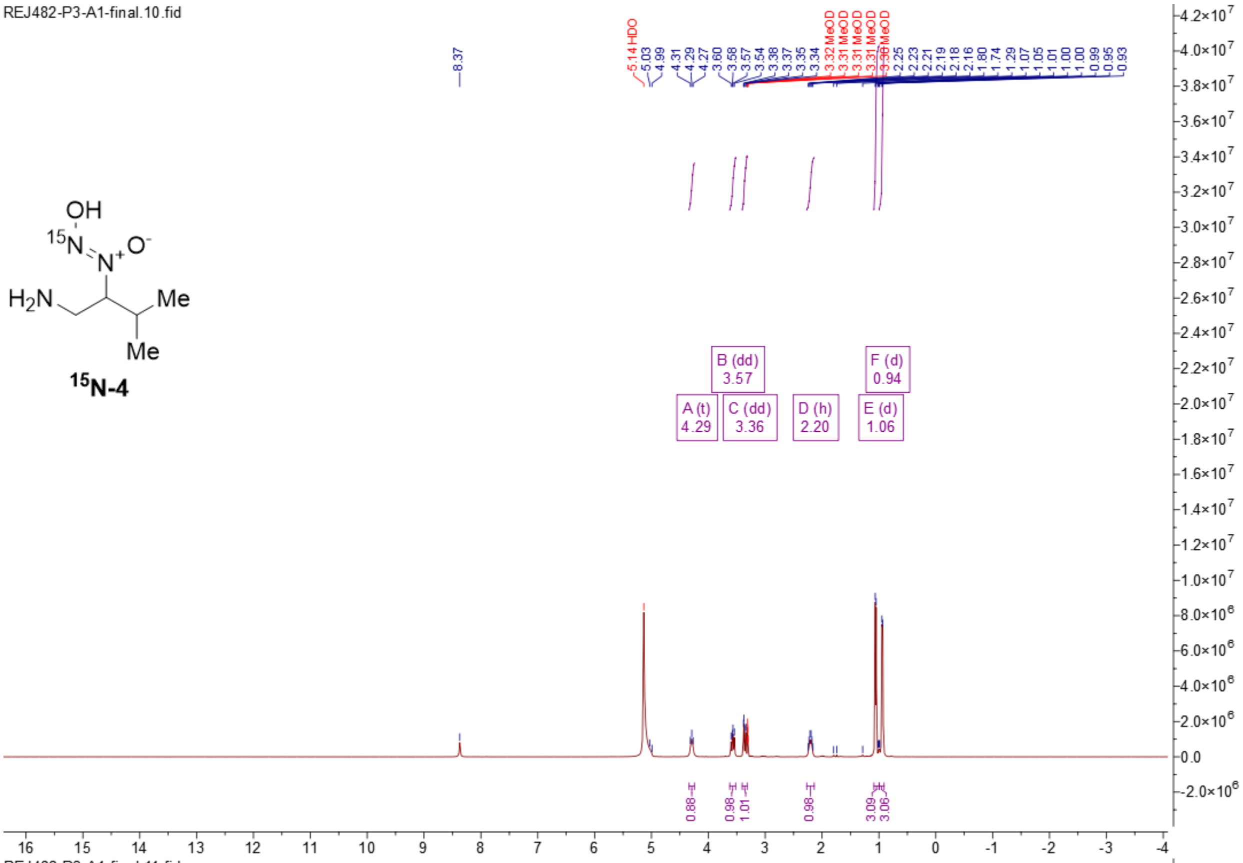
REJ482-P3-A1-final.10.fid
8.37
5.14 HDO
5.03
4.99
4.31
4.29
4.27
3.60
3.58
3.57
3.54
3.38
3.37
3.35
3.34
3.32 MeOD
3.31 MeOD
3.31 MeOD
3.31 MeOD
3.30 MeOD
2.25
2.23
2.21
2.19
2.18
2.16
1.80
1.74
1.29
1.07
1.05
1.01
1.00
0.99
0.95
0.93
OH
15N
N+
O-
H2N
Me
Me
15N-4
B (dd)
3.57
F (d)
0.94
A (t)
4.29
C (dd)
3.36
D (h)
2.20
E (d)
1.06
0.88
0.98
1.01
0.98
3.09
3.06

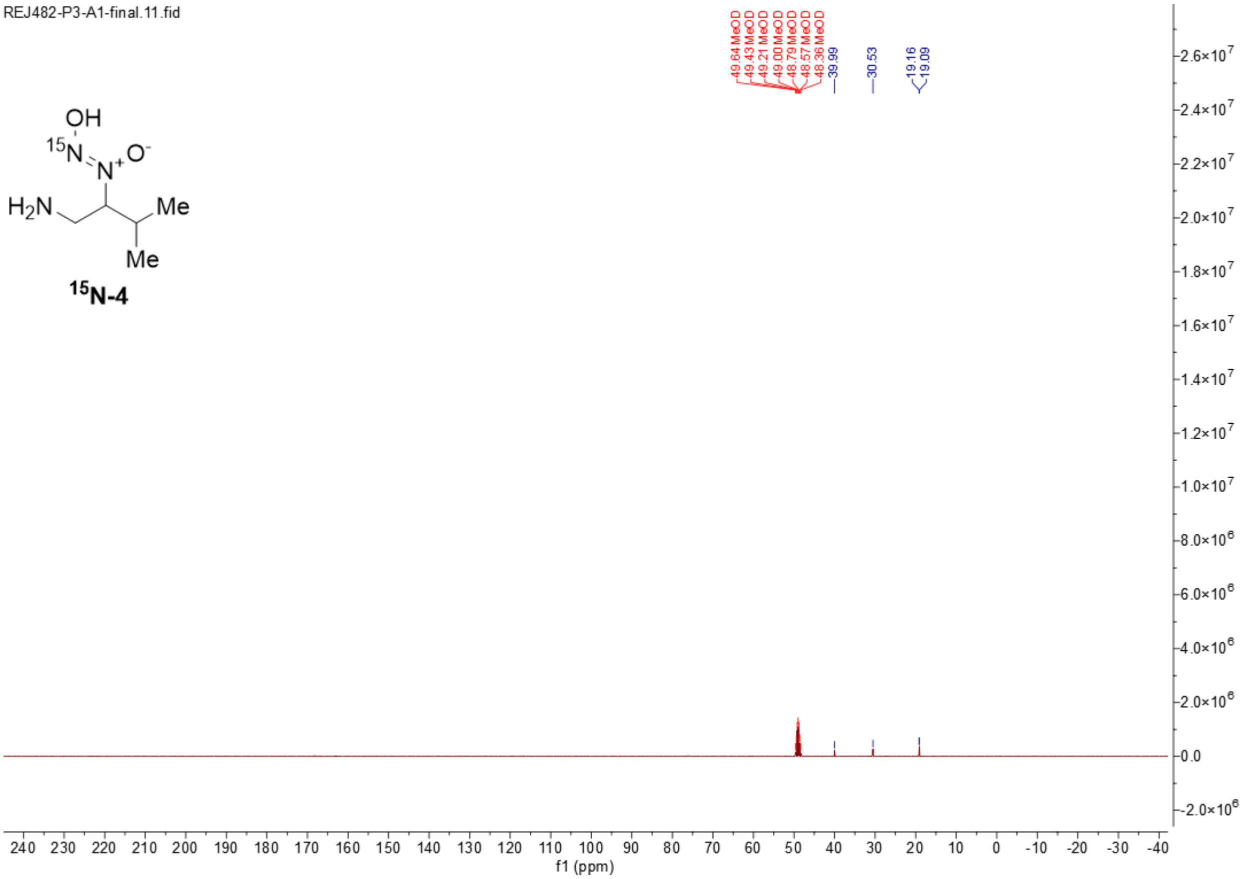
REJ482-P3-A1-final.11.fid
49.64 MeOD
49.43 MeOD
49.21 MeOD
49.00 MeOD
48.79 MeOD
48.57 MeOD
48.36 MeOD
39.99
30.53
19.16
19.09
OH
15N
N+
O-
H2N
Me
Me
15N-4
f1 (ppm)

**Synthesis (*S*)–2-((benzoyloxy)amino)-3-methylbutan-1-ol (((*S*)–13)**

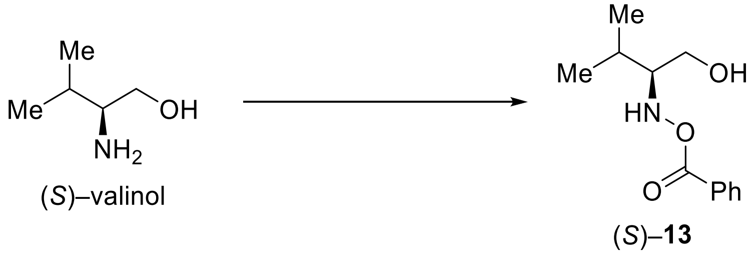


To a suspension of dibenzoylperoxide (611 mg, 2.52 mmol, 1.3 eq.) and $K_2HPO_4$ (541 mg, 3.10 mmol, 1.6 eq.) in dry THF (5.00 mL) at r.t., was added a solution of (*S*)–valinol (200 mg, 1.94 mmol, 1 eq.) in dry THF (2.30 mL) dropwise. The mixture was stirred at r.t. for 16 hours. The mixture was filtered through a short pad of Celite® with the aid of THF. The mixture was concentrated *in vacuo* to give a gummy colourless crude. The crude (655 mg) was loaded onto silica gel (70 mL) and eluted with 10% EtOAc in hexanes (250 mL) then 30% EtOAc in hexanes (200 mL) to give (*S*)–**13** (317 mg, 73%) as a colourless gum. The analytics agree with literature.[18]

$\mathbf{R_f}$ = 0.35 ($SiO_2$, 30% EtOAc in hexanes, UV and $KMnO_4$ stain).

**$^1$H NMR** (400 MHz, $CDCl_3$) $\delta$ = 8.01 (dd, $J$ = 8.0, 1.5 Hz, 2H), 7.64 – 7.55 (m, 1H), 7.46 (t, $J$ = 7.7 Hz, 2H), 3.80 (dd, $J$ = 11.6, 3.5 Hz, 1H), 3.64 (dd, $J$ = 11.6, 7.1 Hz, 1H), 2.87 (td, $J$ = 7.3, 3.5 Hz, 1H), 2.00 – 1.87 (m, 1H), 1.11 (d, $J$ = 6.9 Hz, 4H), 1.03 (d, $J$ = 6.8 Hz, 3H).

**$^{13}$C NMR** (101 MHz, $CDCl_3$) $\delta$ = 167.28, 133.64, 129.54, 128.74, 128.36, 68.60, 60.18, 27.41, 19.83, 19.55.

**ESI-HRMS** (MeCN): $m/z$ 224.12814 ($C_{12}H_{18}O_3N^+$; $[M$+H$]^+$; calc. 224.12812).

**Optical rotation**: $[\alpha]_D^{25}$ = −11.618 (c = 0.90, $CHCl_3$).

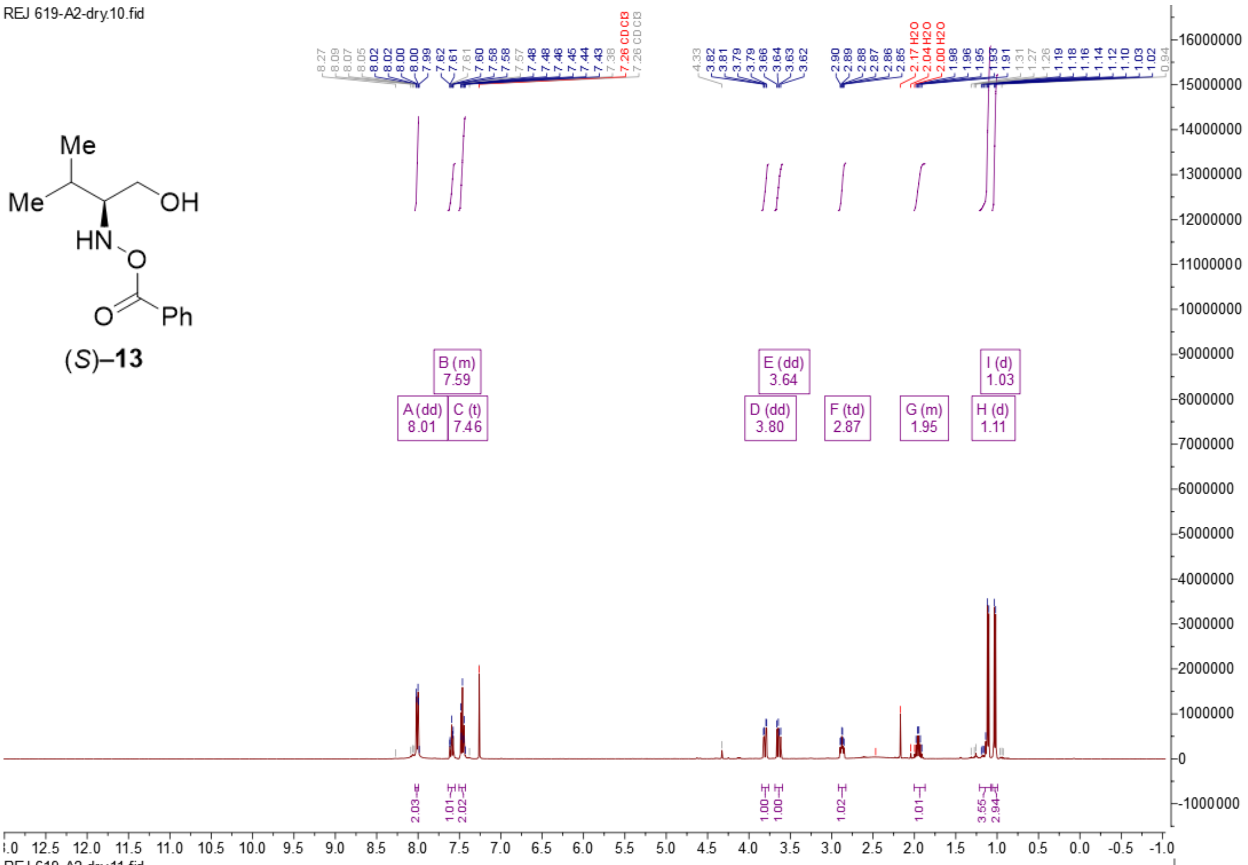
REJ 619-A2-dry.10.fid
Me
Me
OH
HN
O
O
Ph
(S)–13
A (dd) 8.01
B (m) 7.59
C (t) 7.46
D (dd) 3.80
E (dd) 3.64
F (td) 2.87
G (m) 1.95
H (d) 1.11
I (d) 1.03

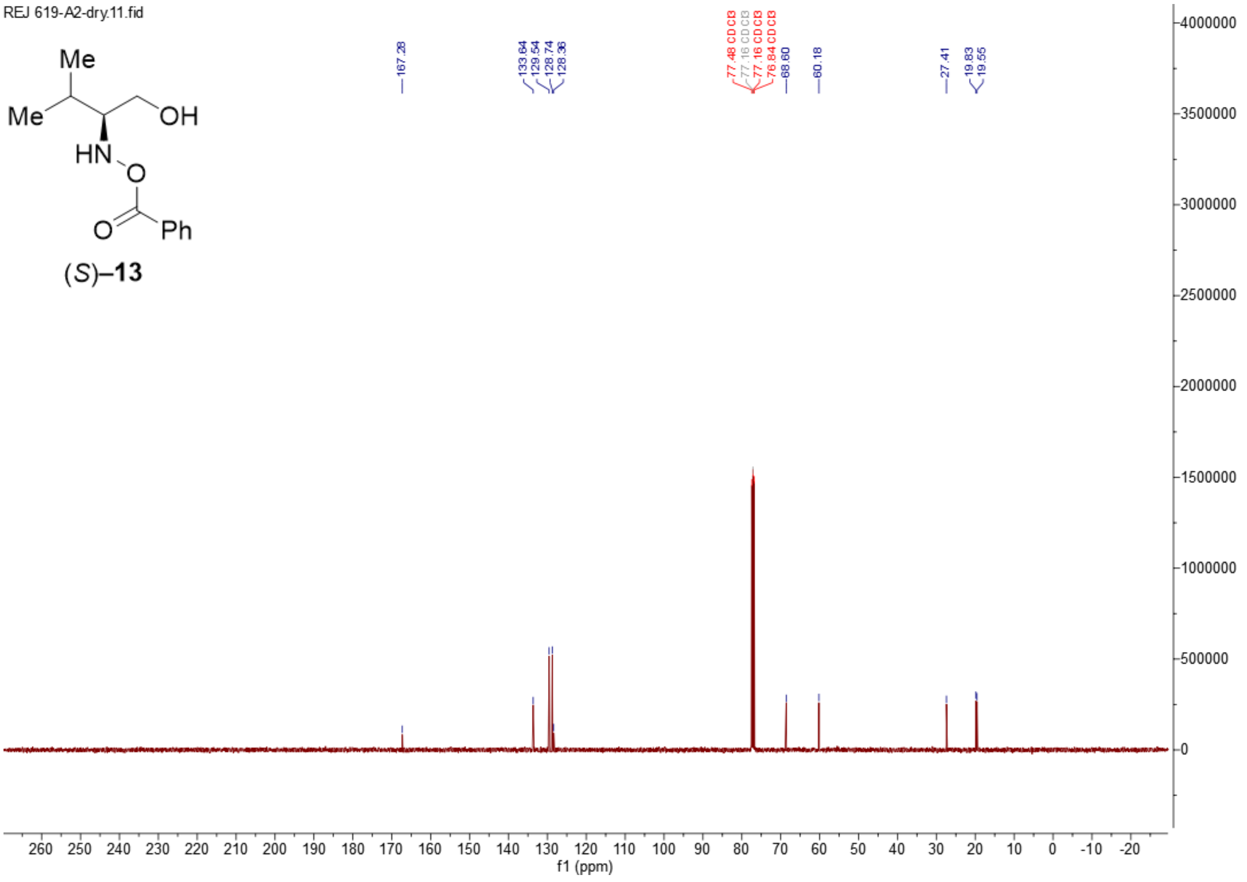
REJ 619-A2-dry.11.fid
Me
Me
OH
HN
O
O
Ph
(S)–13
167.28
133.64
129.54
128.74
128.36
68.60
60.18
27.41
19.83
19.55
f1 (ppm)

**Synthesis of (*S*,*Z*)–1-(1-amino-3-methylbutan-2-yl)-2-hydroxydiazene 1-oxide ((*S*)–4)**

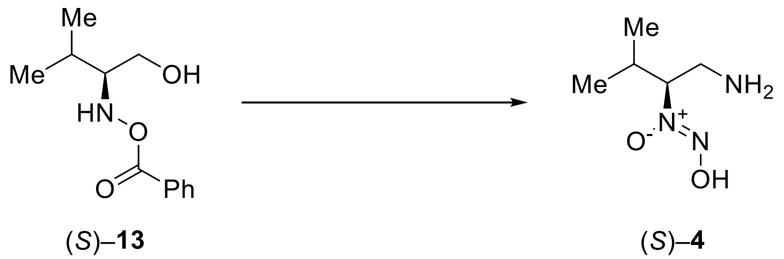

To a solution of alcohol (*S*)–**13** (791 mg, 3.54 mmol, 1 eq.) in dry THF (20.0 mL) under $N_2$ at 0 °C, $PPh_3$ (2.32 g, 8.85 mmol, 2.5 eq.) and *N*-Boc-*tert*-butylcarbamate (1.92 g, 8.85 mmol, 2.5 eq.) were added at 0 °C. The mixture was stirred for 5 minutes before the addition of DEAD (40% in toluene, 3.5 mL, 2.5 eq.) at 0 °C. The reaction was then warmed to r.t. and stirred for 18 hours. TLC in 30% EtOAc in hexanes showed full consumption of the starting material and the formation of many non-polar UV-active spots. The mixture was concentrated *in vacuo* to give a light-yellow crude (2.50 g). The crude was loaded onto silica gel (250 mL) and eluted with 10% EtOAc in hexanes to give an inseparable mixture containing mono-*N*-Boc and bis-*N*-Boc products (330 mg). $^1$H NMR suggested the ratio of bis:mono is about 1:2.

To a solution of the above mixture (330 mg) in MeOH (2.40 mL), was added $K_2CO_3$ (160 mg, 1.16 mmol). The reaction was stirred at r.t. for 2 hours. HPLC-MS showed full deprotection of the benzoyl group. The suspension was filtered with the aid of acetone and the filtrate was concentrated *in vacuo* to give a light yellow solid (524 mg).

To the crude above in DCM (3.7 mL) at 0 °C, was added TFA (1.60 mL, 21.5 mmol). The reaction was stirred at r.t. for 1 hour. HPLC-MS showed full conversion to the desired mass at the solvent front. The mixture was treated with water (5 mL) and separated. The aqueous layer was washed with DCM (2 × 8 mL) and was then concentrated *in vacuo* to give a colourless oil (560 mg). The crude was reconstituted in MiliQ water (0.5 mL) and filtered through DSC-18 cartridge (2 g bed) with the aid of MiliQ water (3 column volumes). The filtrate was concentrated *in vacuo* to give the corresponding β-amino hydroxylamine potentially as a bis-TFA salt (372 mg).

To a solution of the β-amino hydroxylamine TFA salt above (372 mg) in 1:1 EtOH/$H_2O$ (3.60 mL) at 0 °C, was added 1 M aq. HCl (2.30 mL, 2.30 mmol). The mixture was degassed with argon for

10 minutes. In a separate flask, a solution of $NaNO_2$ (82.0 mg, 1.19 mmol) in water (1.00 mL) was degassed with argon for 10 minutes before being added to the solution of hydroxylamine at 0 °C. The mixture was stirred at 0 °C for 30 minutes. HPLC-MS showed full conversion into the desired product mass eluting at the solvent front. The mixture was concentrated *in vacuo* to give a white gummy crude, which was treated with acetone (2 × 0.5 mL), sonicated, and filtered to give a light-yellow gum (400 mg). The gum was reconstituted in MiliQ water (1 mL) and filtered through DSC-18 cartridge (5 g bed) with the aid of MiliQ water (3 column volumes). The filtrate was concentrated *in vacuo* to give a colourless gum (294 mg). The gum was purified by preparative HPLC (*Phenomenex Synergi™* 10 µm Hydro-RP 80 Å, 250 mm × 21.2 mm, 20 mL/min, 100% MiliQ water + 0.1% formic acid, 10 minutes, $R_t$ = 6.5 minutes) to give (*S*)–**4** (48.0 mg, 9% over 4 steps) as a white solid.

**m.p.** = 50.5–54.5 °C

**$^1$H NMR** (400 MHz, Methanol-*d*$_4$) $\delta$ = 4.26 – 4.18 (m, 1H), 3.53 (dd, *J* = 13.8, 9.6 Hz, 1H), 3.36 – 3.32 (m, 1H), 2.21 (dp, *J* = 8.6, 6.7 Hz, 1H), 1.06 (d, *J* = 6.7 Hz, 3H), 0.95 (d, *J* = 6.6 Hz, 3H).

**$^{13}$C NMR** (126 MHz, Methanol-*d*$_4$) $\delta$ = 76.00, 40.09, 30.43, 19.24, 19.10.

**FTIR $\tilde{\nu}$ (cm$^{-1}$)** = 2971br m, 1672m, 1532m, 1433m, 1302w, 1273m, 1183s, 1134s, 1066m, 976m, 924m, 898m, 837m, 798m, 722m, 699m, 598w, 518m, 443w, 406w.

**ESI-HRMS** (MeOH): *m/z* 148.10798 ($C_5H_{14}O_2N_3^+$; $[M+H]^+$; calc. 148.10805).

**Optical rotation**: $[\alpha]_D^{25} = -7.018$ (c = 0.79, methanol).

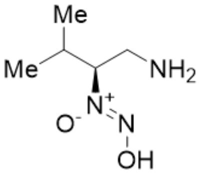

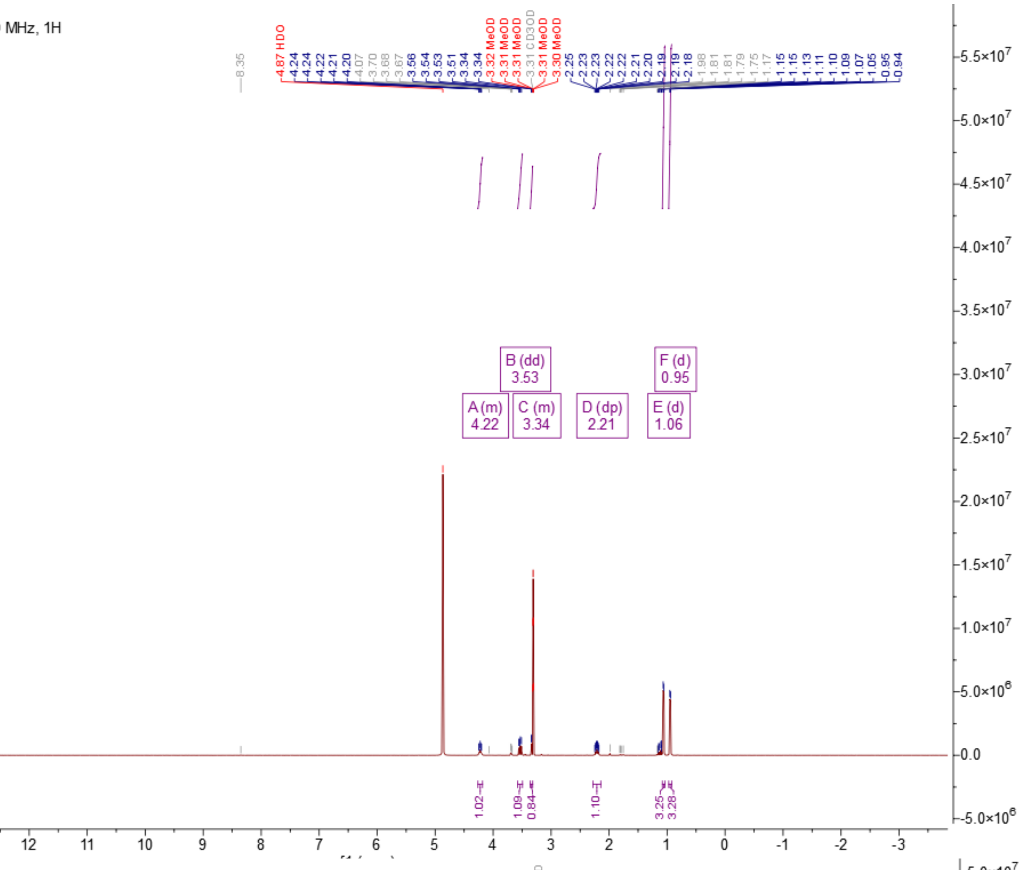

REJ691-P3-A1-prep.1.fid
REJ691-P3-A1-prep, MeOD, 500 MHz, 1H
A (m) 4.22
B (dd) 3.53
C (m) 3.34
D (dp) 2.21
E (d) 1.06
F (d) 0.95


(*S*)–**4**

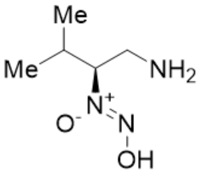

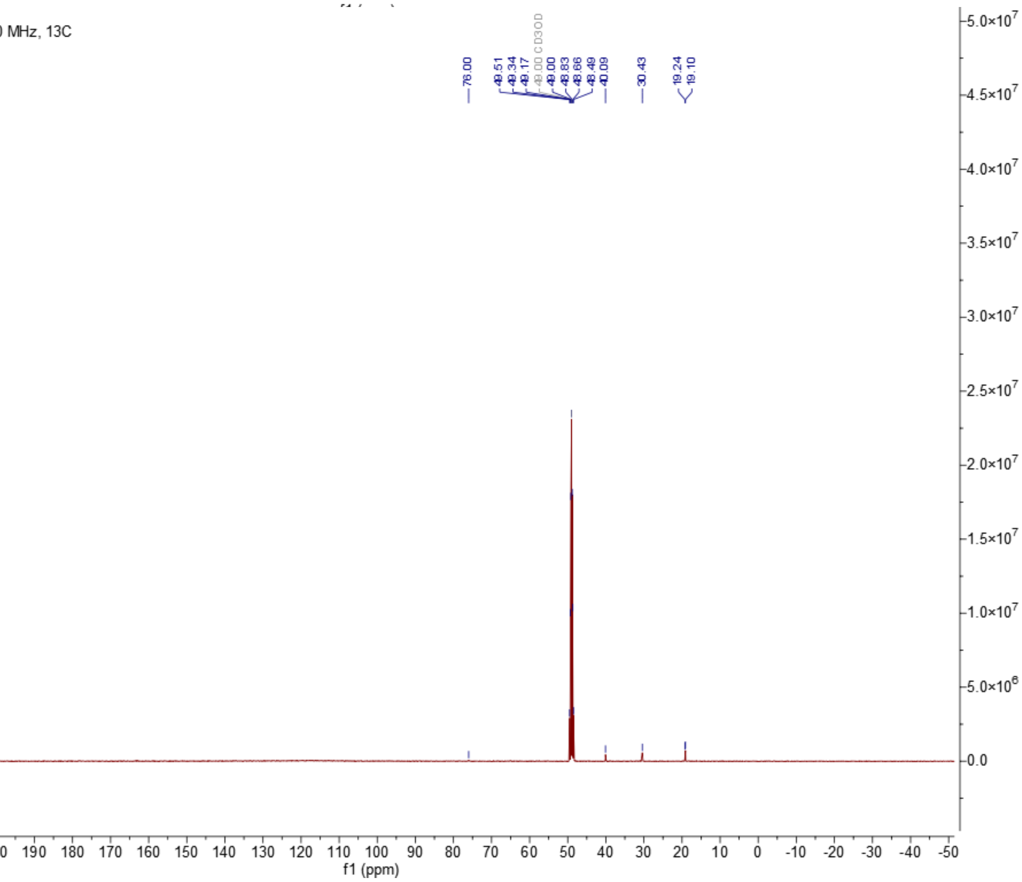

REJ691-P3-A1-prep.3.fid
REJ691-P3-A1-prep, MeOD, 500 MHz, 13C
76.00
40.09
30.43
19.24
19.10
f1 (ppm)


(*S*)–**4**

**Synthesis (*R*)-2-((benzoyloxy)amino)-3-methylbutan-1-ol ((*R*)–13)**

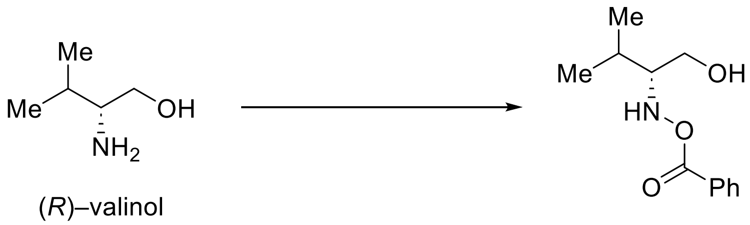

(*R*)–valinol (*R*)–**13**

To a suspension of dibenzoylperoxide (1.53 g, 6.32 mmol, 1.3 eq.) and $K_2HPO_4$ (1.36 g, 7.81 mmol, 1.6 eq.) in dry THF (12.5 mL) at r.t., was added a solution of (*R*)–valinol (500 mg, 4.85 mmol, 1 eq.) in dry THF (5.80 mL) dropwise. The mixture was stirred at r.t. for 16 hours. The mixture was filtered through a short pad of Celite® with the aid of THF. The mixture was concentrated *in vacuo* to give a gummy colourless crude. The crude (1.50 g) was loaded onto silica gel (200 mL) and eluted with 30% EtOAc in hexanes (1 L) to give (*R*)–**13** (824 mg, 76%) as a colourless gum.

$\mathbf{R_f}$ = 0.35 ($SiO_2$, 30% EtOAc in hexanes, UV and $KMnO_4$ stain).

**$^1$H NMR** (400 MHz, $CDCl_3$) $\delta$ = 8.05 – 7.97 (m, 2H), 7.64 – 7.55 (m, 1H), 7.51 – 7.42 (m, 2H), 3.80 (dd, $J$ = 11.7, 3.6 Hz, 1H), 3.64 (dd, $J$ = 11.6, 7.0 Hz, 1H), 2.87 (td, $J$ = 7.3, 3.5 Hz, 1H), 2.04 – 1.87 (m, 1H), 1.11 (d, $J$ = 6.8 Hz, 3H), 1.03 (d, $J$ = 6.9 Hz, 3H).

**$^{13}$C NMR** (101 MHz, $CDCl_3$) $\delta$ = 167.28, 133.64, 129.54, 128.74, 128.36, 68.60, 60.19, 27.41, 19.83, 19.55.

**FTIR $\tilde{\nu}$ (cm$^{-1}$)** = 3243br w, 3065w, 2963m, 2875m, 1720m, 1600m, 1574m, 1451m, 1388m, 1315m, 1270s, 1177m, 1112m, 1069m, 1026m, 975w, 923w, 836w, 781w, 709s, 688m, 542w.

**ESI-HRMS** (MeOH): *m/z* 224.12810 ($C_{12}H_{18}O_3N^+$; [*M*+H]$^+$; calc. 224.12812).

**Optical rotation**: $[\alpha]_D^{25}$ = +11.375 (c = 0.80, $CHCl_3$).

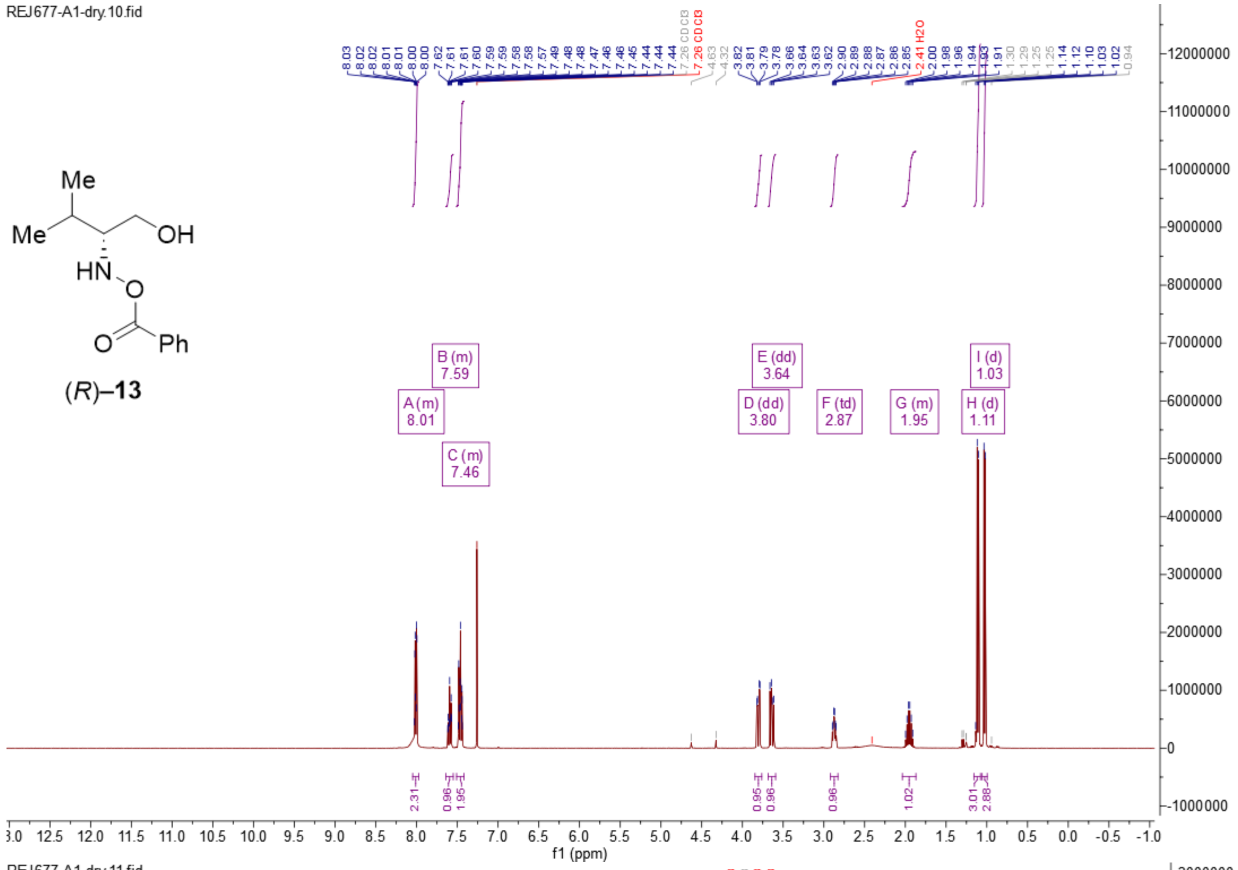
REJ677-A1-dry.10.fid
(R)–13
A (m)
8.01
B (m)
7.59
C (m)
7.46
D (dd)
3.80
E (dd)
3.64
F (td)
2.87
G (m)
1.95
H (d)
1.11
I (d)
1.03
f1 (ppm)

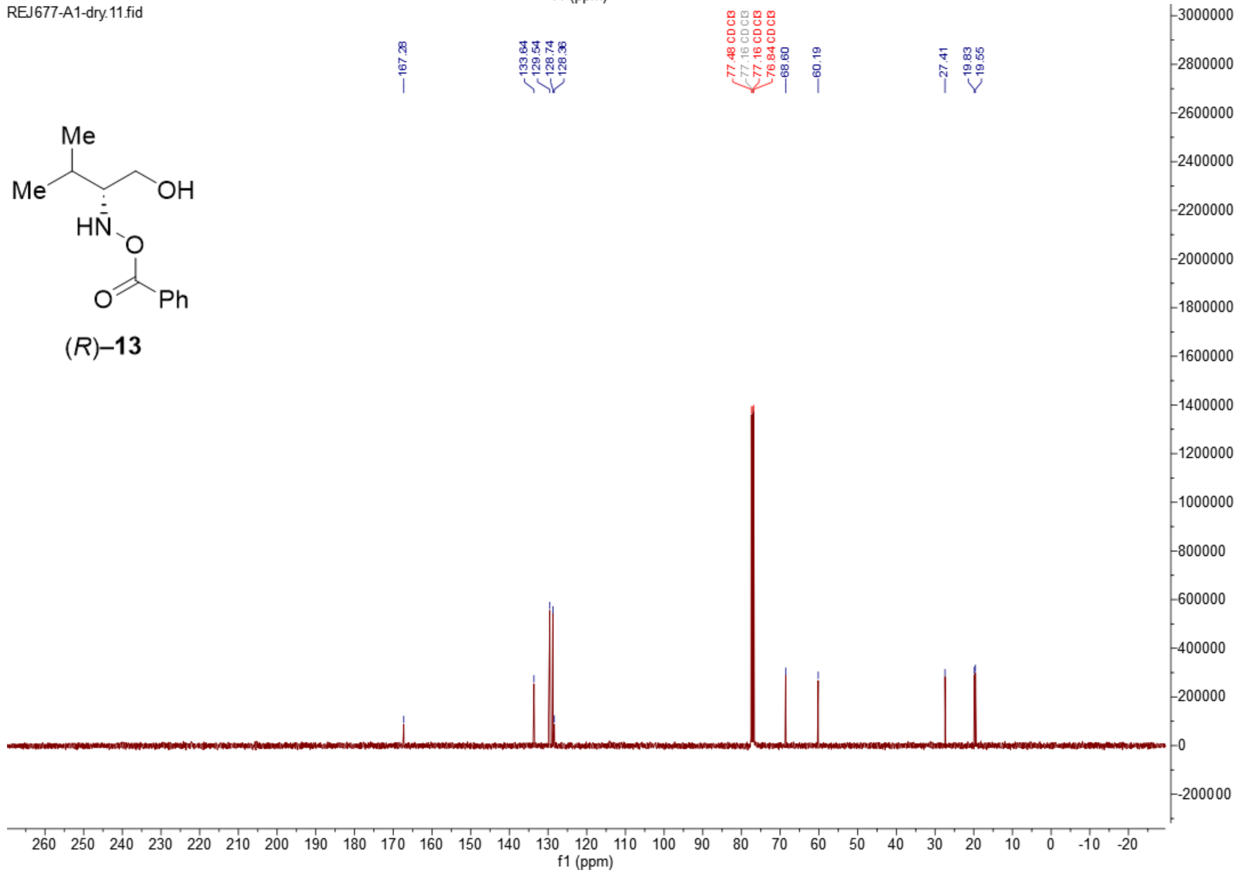
REJ677-A1-dry.11.fid
(R)–13
167.28
133.64
129.54
128.74
128.36
68.60
60.19
27.41
19.83
19.55
f1 (ppm)

**Synthesis of (*R*,*Z*)-1-(1-amino-3-methylbutan-2-yl)-2-hydroxydiazene 1-oxide (((*R*)–4)**

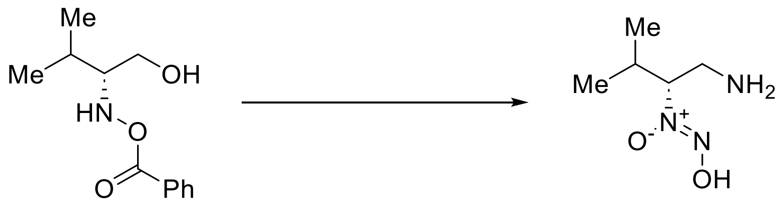

(*R*)–**13** (*R*)–**4**

To a solution of alcohol (*R*)–**13** (750 mg, 3.36 mmol, 1 eq.) in dry THF (19.0 mL) under $N_2$ at 0 °C, $PPh_3$ (2.20 g, 8.40 mmol, 2.5 eq.) and *N*-Boc-*tert*-butylcarbamate (1.83 g, 8.40 mmol, 2.5 eq.) were added at 0 °C. The mixture was stirred for 5 minutes before the addition of DEAD (40% in toluene, 3.3 mL, 2.5 eq.) at 0 °C. The reaction was then warmed to r.t. and stirred for 18 hours. TLC in 30% EtOAc in hexanes showed full consumption of the starting material and the formation of many non-polar UV-active spots. The mixture was concentrated *in vacuo* to give a light-yellow crude (2.10 g). The crude was loaded onto silica gel (200 mL) and eluted with 10% EtOAc in hexanes to give an inseparable mixture containing mono-*N*-Boc and bis-*N*-Boc products (376 mg).

To a solution of the above mixture (376 mg) in MeOH (2.4 mL), was added $K_2CO_3$ (182 mg, 1.32 mmol). The reaction was stirred at r.t. for 2 hours. HPLC-MS showed full deprotection of the benzoyl group. The suspension was filtered with the aid of acetone and the filtrate was concentrated *in vacuo* to give a light yellow solid (507 mg).

To the crude above in DCM (3.70 mL) at 0 °C, was added TFA (1.80 mL, 24.2 mmol). The reaction was stirred at r.t. for 1 hour. HPLC-MS showed full conversion to the desired mass at the solvent front. The mixture was treated with water (5 mL) and separated. The aqueous layer was washed with DCM (2 × 8 mL) and was then concentrated *in vacuo* to give a colourless oil (540 mg). The crude was reconstituted in MiliQ water (0.5 mL) and filtered through DSC-18 cartridge (2 g bed) with the aid of MiliQ water (3 column volumes). The filtrate was concentrated *in vacuo* to give the corresponding β-amino hydroxylamine potentially as a bis-TFA salt (438 mg).

To a solution of the β-amino hydroxylamine TFA salt above (438 mg) in 1:1 EtOH/$H_2O$ (4.20 mL) at 0 °C, was added 1 M aq. HCl (2.70 mL, 2.70 mmol). The mixture was degassed with argon for 10 minutes. In a separate flask, a solution of $NaNO_2$ (97.0 mg, 1.40 mmol) in water (1.00 mL) was degassed with argon for 10 minutes before being added to the solution of hydroxylamine at 0 °C. The mixture was stirred at 0 °C for 30 minutes. HPLC-MS showed full conversion into the desired product mass eluting at the solvent front. The mixture was concentrated *in vacuo* to give a white gummy crude, which was treated with acetone (2 × 0.5 mL), sonicated, and filtered to give a light-yellow gum (404 mg). The gum was reconstituted in MiliQ water (1 mL) and filtered through DSC-18 cartridge (5 g bed) with the aid of MiliQ water (3 column volumes). The filtrate was concentrated *in vacuo* to give a colourless gum (290 mg). The gum was purified by preparative HPLC (*Phenomenex Synergi*$^{TM}$ 10 µm Hydro-RP 80 Å, 250 mm × 21.2 mm, 20 mL/min, 100% MiliQ water + 0.1% formic acid, 10 minutes, $R_t$ = 6.5 minutes) to give (*R*)–**4** (54.0 mg, 11% over 4 steps) as a white solid.

**m.p.** = 51.2–55.7 °C

**$^1$H NMR** (400 MHz, Methanol-$d_4$) $\delta$ = 4.26 – 4.18 (m, 1H), 3.53 (dd, *J* = 13.8, 9.6 Hz, 1H), 3.35 – 3.32 (m, 1H), 2.21 (dp, *J* = 8.5, 6.6 Hz, 1H), 1.06 (d, *J* = 6.8 Hz, 3H), 0.95 (d, *J* = 6.6 Hz, 3H).

**$^{13}$C NMR** (126 MHz, Methanol-$d_4$) $\delta$ = 75.98, 40.10, 30.42, 19.24, 19.10.

**FTIR $\tilde{\nu}$ (cm$^{-1}$)** = 2971br m, 1672m, 1532m, 1432m, 1396m, 1302w, 1272m, 1181s, 1133s, 1066m, 976m, 924m, 897m, 836m, 798m, 722m, 699m, 598w, 517m, 443w, 411w.

**ESI-HRMS** (MeOH): *m/z* 148.10810 ($C_5H_{14}O_2N_3^+$; [*M*+H]$^+$; calc. 148.10805).

**Optical rotation**: $[\alpha]_D^{25}$ = +8.085 (c 0.80, MeOH).

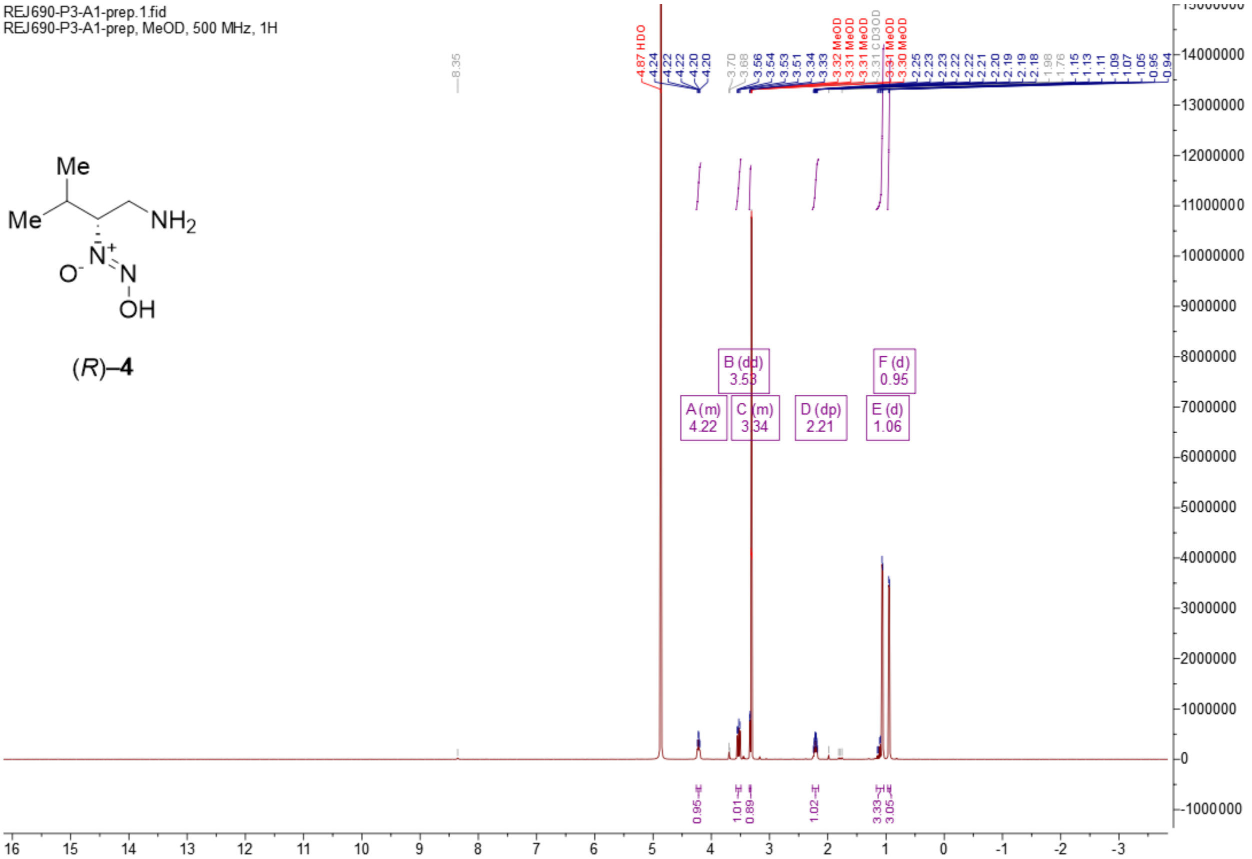
REJ690-P3-A1-prep.1.fid
REJ690-P3-A1-prep, MeOD, 500 MHz, 1H
Me
Me
NH2
N+
O-
N
OH
(R)–4
8.35
4.87 HDO
4.24 4.22 4.22 4.20 4.20
3.70 3.68 3.56 3.54 3.53 3.51 3.34 3.33
3.32 MeOD 3.31 MeOD 3.31 MeOD 3.31 CD3OD 3.31 MeOD 3.30 MeOD
2.25 2.23 2.23 2.22 2.22 2.21 2.20 2.19 2.19 2.18 1.98 1.76 1.15 1.13 1.11 1.09 1.07 1.05 0.95 0.94
A (m) 4.22
B (dd) 3.53
C (m) 3.34
D (dp) 2.21
E (d) 1.06
F (d) 0.95
0.95 1.01 0.89 1.02 3.33 3.05
16 15 14 13 12 11 10 9 8 7 6 5 4 3 2 1 0 -1 -2 -3
14000000 13000000 12000000 11000000 10000000 9000000 8000000 7000000 6000000 5000000 4000000 3000000 2000000 1000000 0 -1000000

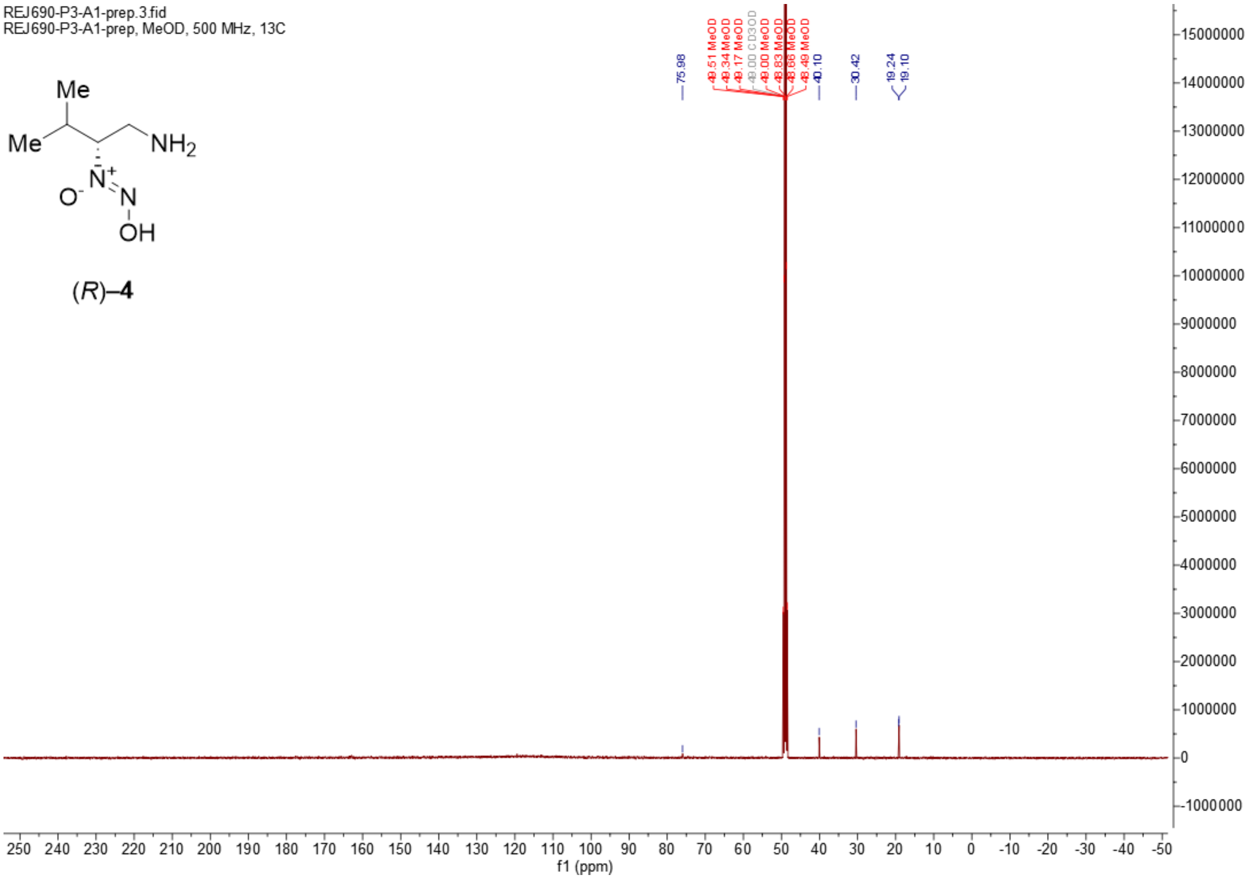
REJ690-P3-A1-prep.3.fid
REJ690-P3-A1-prep, MeOD, 500 MHz, 13C
Me
Me
NH2
N+
O-
N
OH
(R)–4
75.98
49.51 MeOD 49.34 MeOD 49.17 MeOD 49.00 CD3OD 49.00 MeOD 48.83 MeOD 48.66 MeOD 48.49 MeOD
40.10
30.42
19.24 19.10
250 240 230 220 210 200 190 180 170 160 150 140 130 120 110 100 90 80 70 60 50 40 30 20 10 0 -10 -20 -30 -40 -50
f1 (ppm)
15000000 14000000 13000000 12000000 11000000 10000000 9000000 8000000 7000000 6000000 5000000 4000000 3000000 2000000 1000000 0 -1000000

**Synthetic attempt towards enantiomerically pure aldehyde (2)**

**Synthesis of methyl hydroxy-L-valinate (15)**

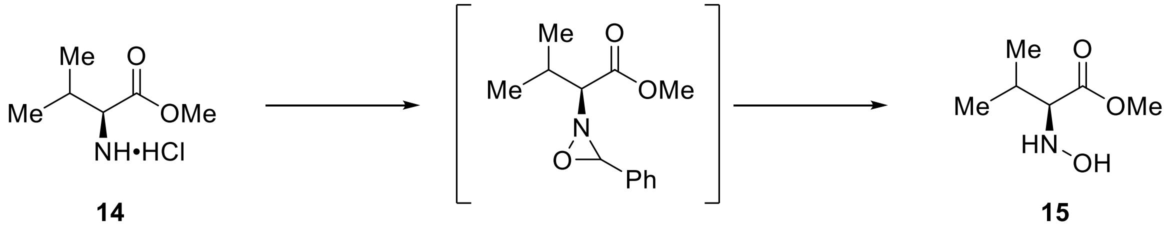

This experiment was conducted using a modified published procedure.[19] To a solution of L–valine methyl ester hydrochloride **14** (6.25 g, 37.3 mmol, 1 eq.) in dry methanol (120 mL), dry $Na_2CO_3$ (7.00 g, 55.9 mmol, 1.5 eq.) and *p*-anisaldehyde (4.60 mL, 37.0 mmol, 1 eq.) were added and the reaction mixture was stirred for 18 hours at r.t. followed by 1 hour at 40 °C. The reaction mixture was filtered through a short pad of Celite® and concentrated *in vacuo*. The residues were redissolved in $Et_2O$, filtered and concentrated *in vacuo* to give a slightly yellow crude product, which was used without further purification in the next step. The imine was dissolved in dry DCM (30 mL) and cooled to −5 °C in an ice/NaCl bath. A solution of *m*-CPBA (8.02 g, 37.3 mmol, 1 eq.) in DCM was dried over $MgSO_4$, filtered and added dropwise to the reaction mixture over 1 hour. The reaction was allowed to warm to r.t. and was stirred for 24 hours. The reaction mixture was filtered, washed with a sat. aq. solution of $NaHCO_3$ (3 × 80 mL) and brine (80 mL), dried over anhydrous $Na_2SO_4$, filtered, and concentrated *in vacuo* to afford the crude oxaziridine, which was used without further purification in the next step. The oxaziridine was dissolved in dry MeOH (80 mL) and treated with hydroxylamine hydrochloride (5.23 g, 74.6 mmol, 2 eq.) at r.t. for 24 hours. The solvent was removed under reduced pressure, and the residues were suspended in $H_2O$ and washed with $Et_2O$ (3 × 80 mL). The aqueous layer was neutralized with $NaHCO_3$ and extracted with $Et_2O$ (3 × 80 mL). The combined organic layers were dried over $Na_2SO_4$, filtered, and concentrated *in vacuo* to obtain the hydroxylamine **15** as a slightly yellow solid (3.16 g, 58% over 3 steps).

$\mathbf{R_f}$ = 0.22 ($SiO_2$, 50% EtOAc in pentane, $KMnO_4$ stain).

**m.p.** = 157–161 °C

$^{1}$**H NMR** (400 MHz, Methanol-*d₄*): $\delta$ = 3.74 (s, 3H), 1.93 – 1.78 (m, $J$ = 6.9 Hz, 1H), 0.99 (d, $J$ = 6.9 Hz, 3H), 0.92 (d, $J$ = 6.8 Hz, 3H).

$^{13}$**C NMR** (101 MHz, Methanol-$d_4$): $\delta$ = 175.82, 73.08, 52.00, 30.09, 19.94, 19.49.

**FTIR $\tilde{\nu}$ (cm$^{-1}$)** = 3259m, 3181br m, 2953m, 1742s, 1430m, 1370m, 1273m, 1185m, 1156s, 1030m, 992m, 764m, 683m, 524m, 533m.

**Optical rotation**: $[\alpha]_D^{25}$ = +9.200 (c 0.80, $CH_2Cl_2$).

**ESI-HRMS** (MeOH): *m/z* 148.09662 ($C_6H_{14}NO_3^+$; [*M*+H]$^+$; calc. 148.09682).

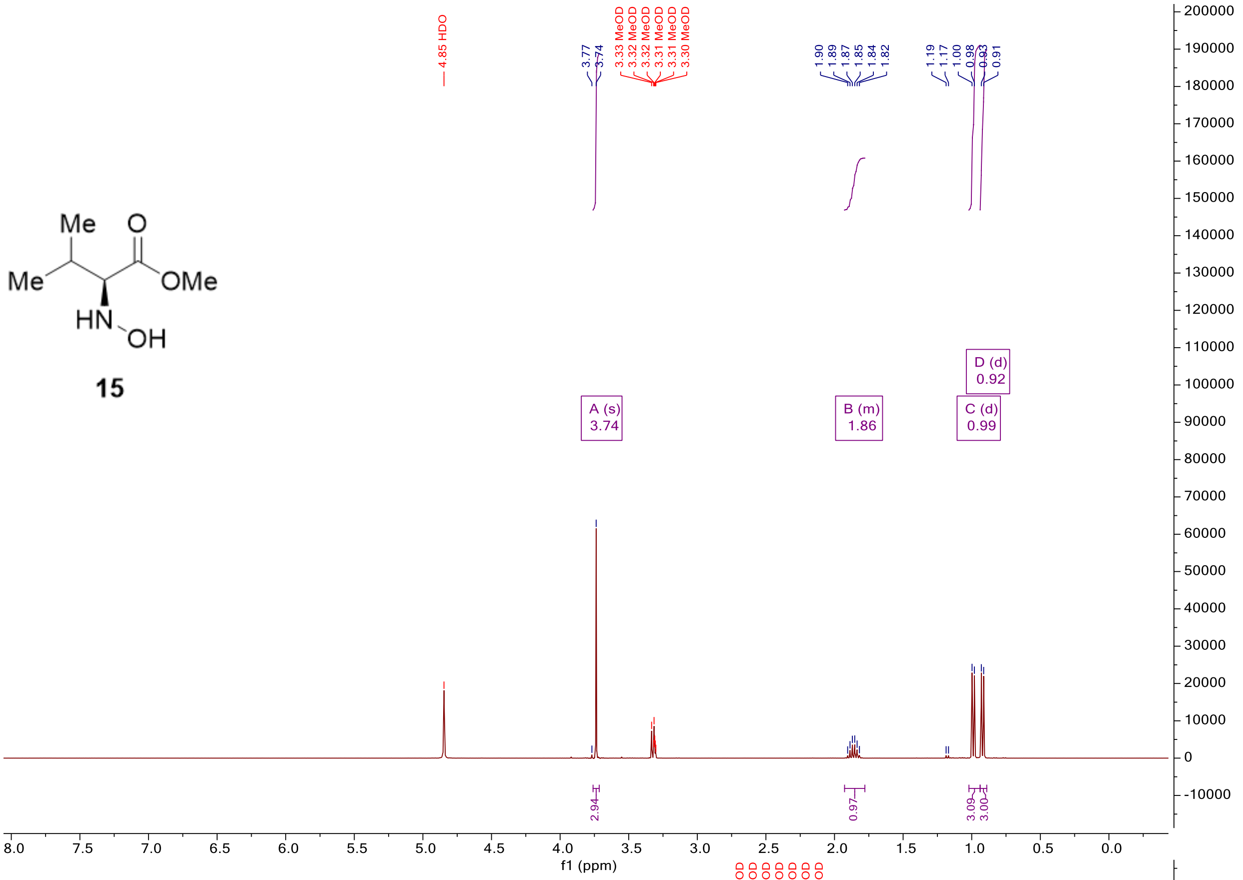
Me
O
Me
OMe
HN
OH
15
4.85 HDO
3.77
3.74
3.33 MeOD
3.32 MeOD
3.32 MeOD
3.31 MeOD
3.31 MeOD
3.30 MeOD
1.90
1.89
1.87
1.85
1.84
1.82
1.19
1.17
1.00
0.98
0.93
0.91
A (s)
3.74
B (m)
1.86
C (d)
0.99
D (d)
0.92
2.94
0.97
3.09
3.00
f1 (ppm)

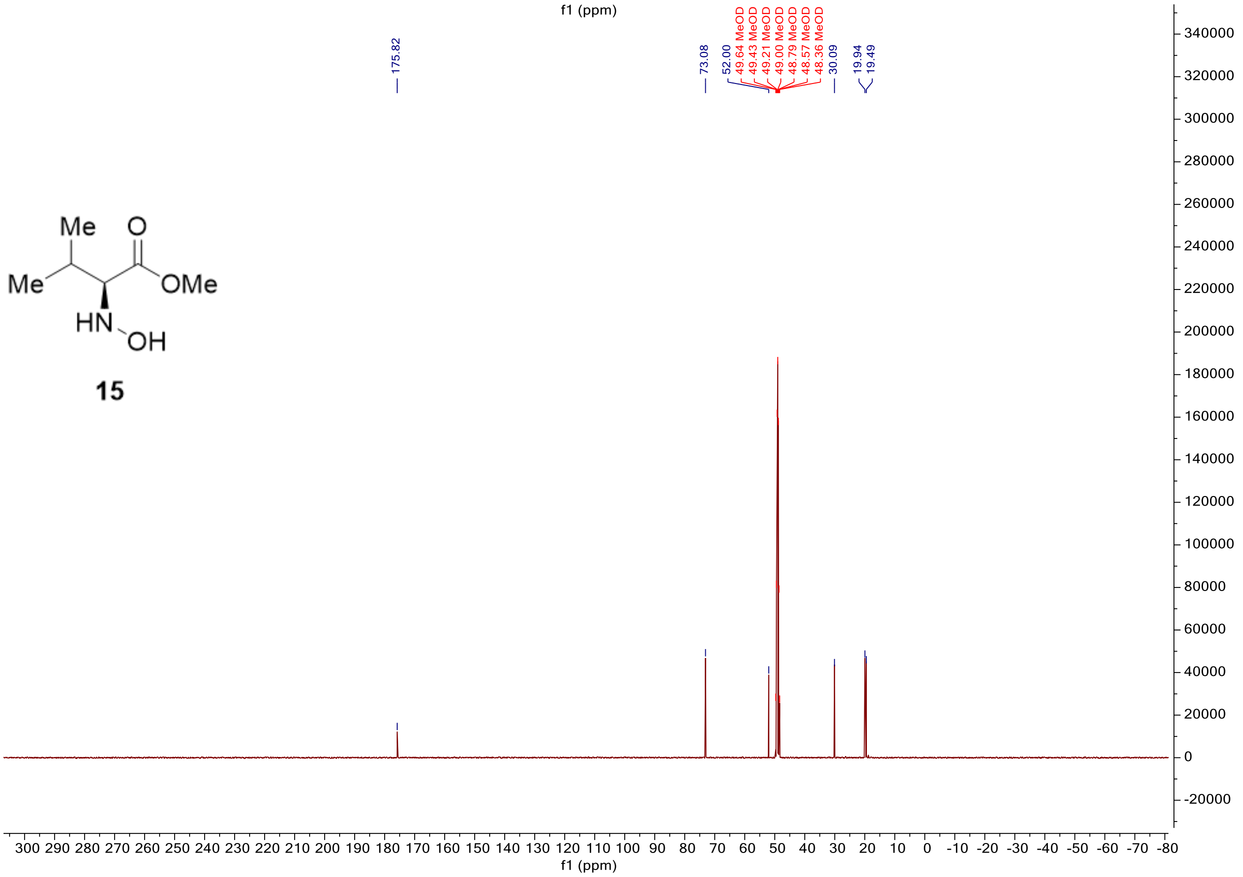
Me
O
Me
OMe
HN
OH
15
175.82
73.08
52.00
49.64 MeOD
49.43 MeOD
49.21 MeOD
49.00 MeOD
48.79 MeOD
48.57 MeOD
48.36 MeOD
30.09
19.94
19.49
f1 (ppm)

**Synthesis of (*Z*)-2-(allyloxy)-1-(3-methyl-1-oxobutan-2-yl)diazene 1-oxide (16) and (*Z*)-2-(allyloxy)-1-(1,1-dihydroxy-3-methylbutan-2-yl)diazene 1-oxide (17)**

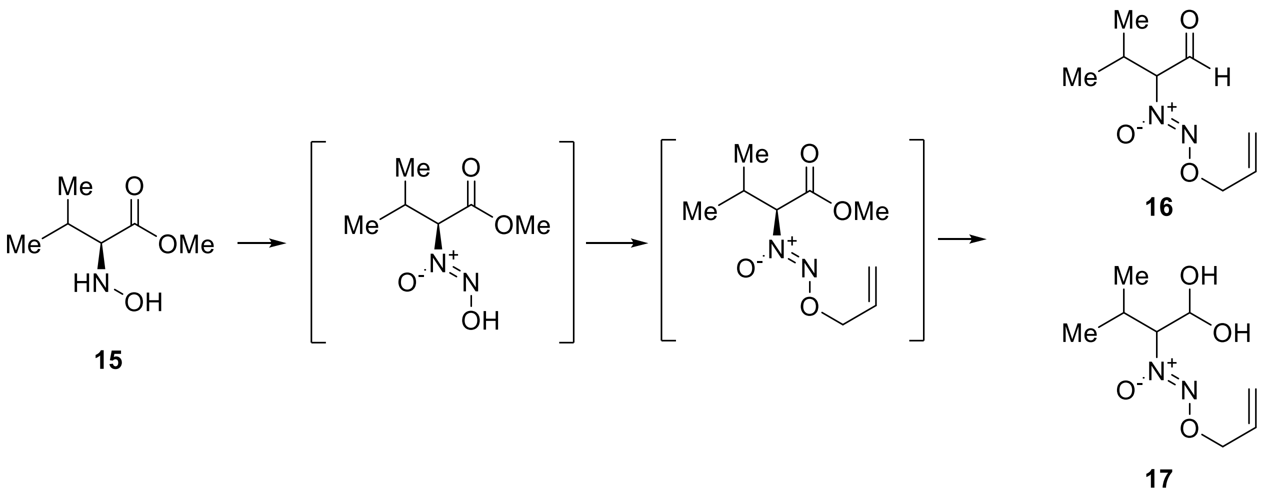


This experiment was conducted using a modified published procedure.[20,21] To a solution of (*S*)–methyl hydroxy valinate **15** (500 mg, 3.40 mmol, 1 eq.) in 1:1 EtOH/$H_2O$ (15.0 mL), was added 4 M aqueous HCl (850 µL, 3.40 mmol, 1 eq.) at 0 °C. The mixture was degassed with argon for 10 minutes. In a separate flask, a solution of $NaNO_2$ (235 mg, 3.40 mmol) in water (3.00 mL) was degassed with argon for 10 minutes before being added dropwise with a syringe pump over 45 minutes to the solution of hydroxylamine at 0 °C. The mixture was stirred at 0 °C for 30 minutes. The yellow solution was diluted with $H_2O$ (30 mL) and extracted with DCM (3 × 30 mL). The combined organic layers were washed with brine (30 mL), dried over anhydrous $Na_2SO_4$, filtered, and concentrated *in vacuo*. Full removal of the solvent was avoided, due to the observed spontaneous decomposition. The green crude oil was used in the next step without further purification. The crude diazeniumdiolate and $Na_2CO_3$ (324 mg, 3.40 mmol, 1 eq.) were dissolved in dry DMF (5.00 mL). Allyl bromide (186 µL, 2.15 mmol) was added at 0 °C and the reaction was allowed to warm to r.t. and stirred for 12 hours. The reaction mixture was filtered through a short pad of Celite® and concentrated *in vacuo* to obtain a mixture of two compounds diverging slightly in the H and C NMR signal for the isopropyl moiety similar $R_f$ on TLC. Unfortunately, those compounds (235 mg) were found as a yellow oil, could not be further separated and were directly used as starting material in the next step.

To a solution of the crude protected diazeniumdiolates (95.0 mg.) in dry DCM (12.0 mL) at –78 °C, was added 1 M DIBAL-H in hexanes (1.20 mL, 1.20 mmol, 2.7 eq.). The reaction was stirred at –78 °C for 1 hour and quenched with EtOAc (3 mL) followed by the addition of 1 M aq. NaOH (3 mL). The reaction was allowed to warm to r.t., treated with $H_2O$ (20 mL), and extracted with DCM (3 × 20 mL). The combined organic layers were dried over anhydrous $MgSO_4$, filtered, and concentrated *in vacuo*. The residue was purified by column chromatography with 20% – 50% ether in pentane to obtain a complex mixture of aldehydes **16** and aldehydes hydrate **17** as a yellow oil (83 mg). The aldehyde was identified by the characteristic peak at 9.64 ppm in the $^1$H NMR and at 195.3 ppm in the $^{13}$C NMR. The presence of the diol was determined by the presence of two doublets at 6.45 and 6.29 ppm standing for the two OH groups and with the down shielded carbon at 88.0 ppm in the $^{13}$C NMR. The identity of the diol was further corroborated by an X-ray analysis of a single crystal obtained by treating the compound with pentane/diethyl ether mixture at –20 °C. Due to the complexity of those mixtures, another route was designed for the project.

**Characterization of the 16 and 17 mixture**

**$R_f$** = 0.2 ($SiO_2$, 50% ether in pentane, $KMnO_4$ stain).

**$^1$H NMR** (500 MHz, DMSO-*d*$_6$) $\delta$ 9.64 (d, $J$ = 0.9 Hz, 1H), 6.29 (d, $J$ = 7.1 Hz, 1H), 6.20 (d, $J$ = 6.4 Hz, 1H), 6.01 – 5.88 (m, 2H), 5.40 – 5.22 (m, 4H), 5.11 (q, $J$ = 6.9 Hz, 1H), 4.85 (dd, $J$ = 8.0, 0.9 Hz, 1H), 4.77 (d, $J$ = 5.7 Hz, 2H), 4.68 (d, $J$ = 5.5 Hz, 1H), 3.79 (dd, $J$ = 7.5, 5.8 Hz, 1H), 2.49 – 2.42 (m, 1H), 2.18 (pd, $J$ = 6.9, 5.8 Hz, 1H), 1.05 (d, $J$ = 6.8 Hz, 3H), 0.95 – 0.89 (m, 9H).

**$^{13}$C NMR** (126 MHz, DMSO-*d*$_6$) $\delta$ 195.3, 133.0, 132.7, 119.1, 118.5, 88.0, 83.8, 81.8, 73.9, 73.2, 27.7, 27.6, 19.5, 18.4, 18.3, 17.6.

**Optical rotation**: $[\alpha]_D^{25}$ = +1.5 (c 0.84, $CH_2Cl_2$).

**ESI-HRMS** (MeOH): aldehydes (**16**): *m/z* 209.08965 ($C_8H_{14}N_2NaO_3^+$; [*M*+Na]$^+$; calc. 209.08966) and diols (**17**): *m/z* 227.10016 ($C_8H_{16}N_2NaO_4^+$; [*M*+Na]$^+$; calc. 227.10023)

**X-ray crystallography**: Structure obtained after analysis of a single crystal of the diol (see **Supplementary Table 5**)

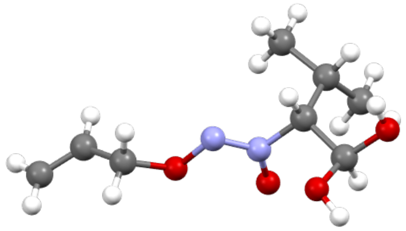

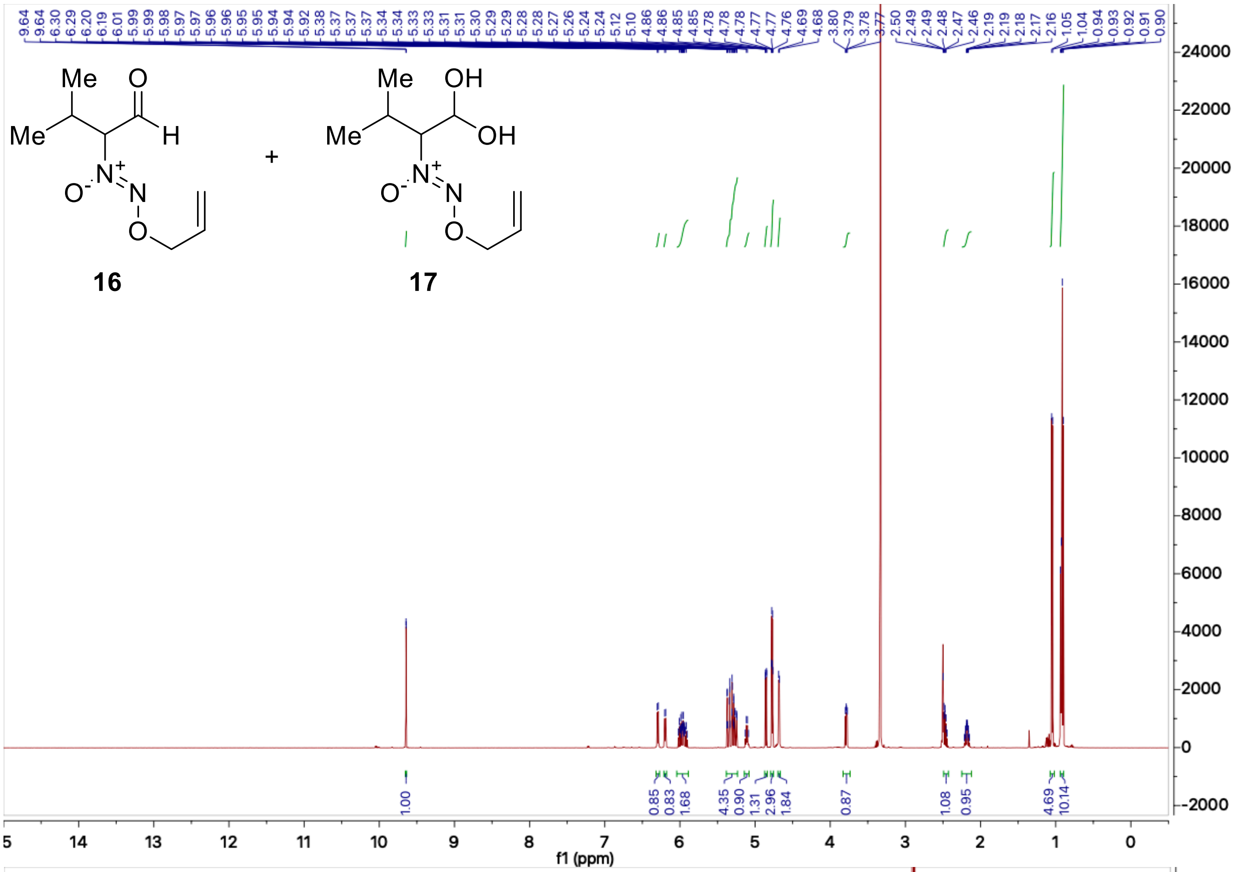

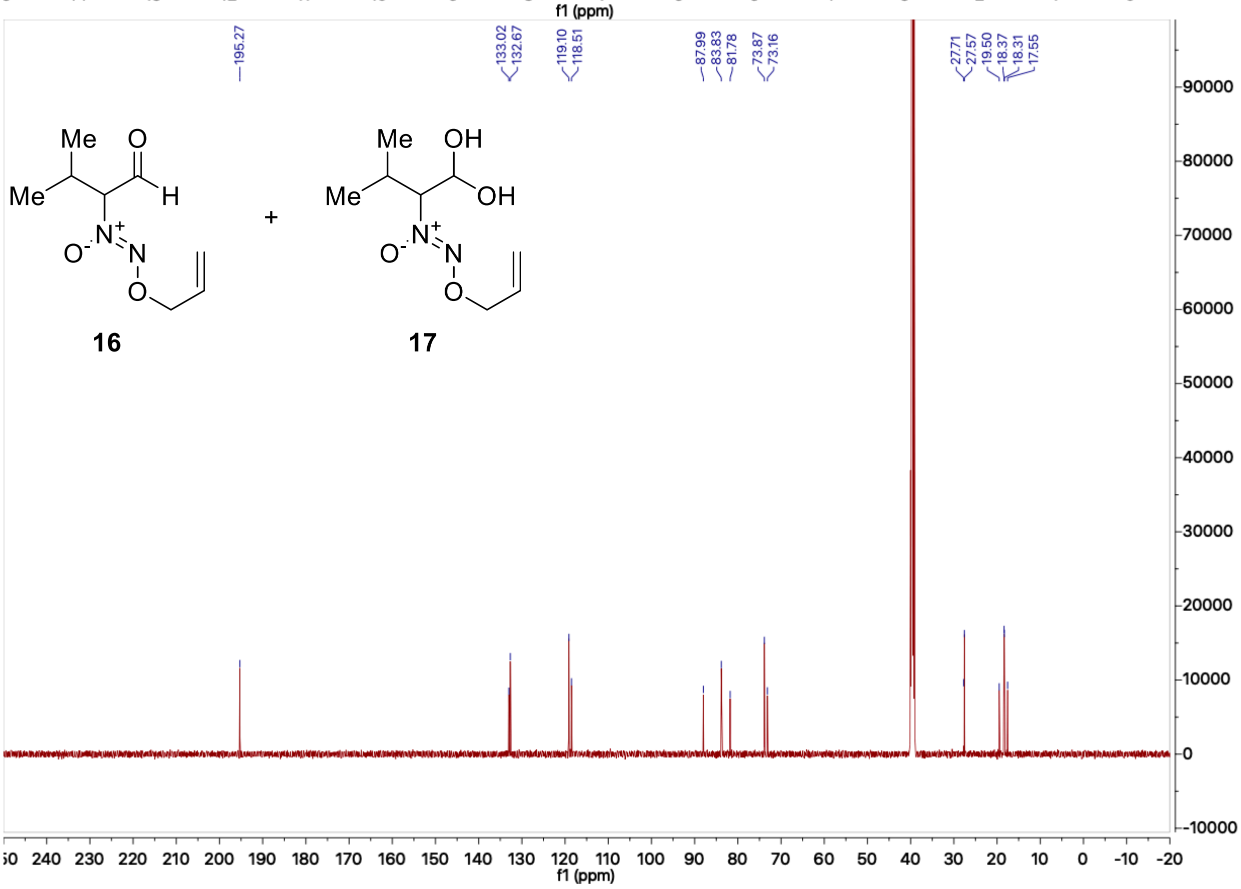
